# PICASSO: Ultra-multiplexed fluorescence imaging of biomolecules through single-round imaging and blind source unmixing

**DOI:** 10.1101/2021.01.27.428247

**Authors:** Junyoung Seo, Yeonbo Sim, Jeewon Kim, Hyunwoo Kim, In Cho, Young-Gyu Yoon, Jae-Byum Chang

## Abstract

Ultra-multiplexed fluorescence imaging of biomolecules is essential to studying heterogeneous biological systems. However, this is challenging due to fluorophores’ spectral overlap and variation of the emission spectra. Here, we propose a strategy termed PICASSO, which enables more than 15-colour multiplexed imaging of thick tissue slices through a single imaging process and blind unmixing without reference spectra measurement. We show that PICASSO can be used to achieve a high multiplexing capability in diverse applications, such as 3D protein imaging, expansion microscopy, tissue clearing, imaging of clinical specimens, and cyclic immunofluorescence imaging. PICASSO only requires an equal number of images as the number of fluorophores, enabling such a high level of multiplexed imaging even with bandpass filter-based microscopy. As such, PICASSO would be a useful tool for the study of cancer, the immune system, and the brain, as well as for the diagnosis of cancer, as it enables ultra-multiplexed imaging of diverse specimens with minimum instrumental requirements and experimental processes.

## INTRODUCTION

Simultaneous imaging of more than 10 proteins in a single specimen can reveal the molecular architecture of highly heterogeneous systems^1–5^. To achieve this kind of multiplexing with fluorescence microscopy, it is necessary to use spectrally overlapping fluorophores and decompose or unmix their signals into each one^6^. Despite recent advancements in spectral imaging and decomposition, exemplified by linear unmixing and blind source separation, the simultaneous imaging of more than 10 proteins has not yet been achieved with these two approaches^3–5^.

The biggest hurdle for multiplexing more than 10 proteins *via* spectral imaging and linear unmixing is the difficulty of measuring precise emission spectra, especially in immunohistochemistry imaging of highly heterogeneous specimens. Mathematically, linear unmixing is formulated as an inverse problem, specifically *D* = *M* × *F*, where *D* and *F* are the acquired mixed and resulting unmixed images, respectively, and *M* is the mixing matrix^7^. For *M*, the reported emission spectra cannot be used because they depend on multiple parameters including the instrumentation, image acquisition setting, and chemical environment involved (see **Supplementary Note 1** for details)^6,8–12^. Particularly in thick tissue, such as mouse brain, fluorophore emission spectra exhibit high levels of variation, presumably affected by the degree of photobleaching, the target proteins, and the subregions where the spectra were acquired, and such variation affects unmixing success (**Supplementary Figs. 1, 2, 3**). As a result, reference spectra need to be measured from all subregions of interest, either from single-fluorophore areas of the target specimen or from separate specimens that have been prepared in the exact same way but each with only one fluorophore. However, the first approach becomes challenging when the number of fluorophores increases beyond 10 and the protein expression patterns overlap, which happens frequently in immunofluorescence imaging of the brain. Reference measurements from separate specimens are also challenging when the number of fluorophores increases beyond 10 because the number of specimens required increases too. In addition, reference spectra measurements from singly labelled separate specimens is sometimes not possible due to limited availability of identical specimens.

Blind source separation, on the other hand, can unmix signals without reference spectra. In this approach, both *M* and *F* are determined from *D* based on *D* = *M* × *F* and can therefore be considered a matrix factorization problem. Unfortunately, this factorization problem is ill-posed since it does not have a unique solution, and so employing proper constraints on *M* and *F* is critical to finding satisfactory explication. Notably, non-negative matrix factorization (NMF) and its variants have shown remarkable success in hyperspectral data and bioinformatics, because *D, M*, and *F* are all non-negative^13,14^. However, its success in multiplexed fluorescence imaging has been limited. NMF does not have a unique solution and typically requires that the number of observations (i.e. images) is sufficiently larger than the number of source signals (i.e. fluorophores)^15,16^, which necessitates microscopy capable of capturing the spectral profiles of specimens. Even with spectral detectors, NMF requires human intervention for the reliable unmixing of four fluorophores^17^ and bleed-through has been observed in seven-colour multiplexed imaging of thin tissue slices^18^ (see **Supplementary Fig. 4** for our NMF unmixing results).

We therefore propose a new blind unmixing technique, PICASSO, which can robustly unmix *N* spectrally overlapping fluorophores without reference measurements and with a number of images equal to the number of fluorophores. By combining PICASSO with an antibody complex formation technique, we demonstrate 15-colour multiplexed imaging of the mouse brain in a single staining and imaging round without emission spectra measurement and not limited by the availability of host species of antibodies. We show that PICASSO can be used for multiplexed 3D imaging, mRNA imaging, super-resolution imaging through tissue expansion, tissue clearing, and for the multiplexed imaging of clinical specimens. We also show that PICASSO can improve the multiplexing capability of cyclic immunofluorescence imaging techniques by allowing them to use more fluorophores in one cycle. Lastly, we show that PICASSO can be implemented with emission filter-based microscopy because it requires an equal number of image acquisitions and fluorophores without the need to use spectral detectors.

## RESULTS

### Multiplexed fluorescence imaging *via* PICASSO

PICASSO can blindly unmix five or more spectrally overlapping fluorophores from the equal number of images acquired at different detection channels. In the experimental implementation of PICASSO, spectrally overlapping fluorophores that can be strongly excited by the same excitation source are selected from various commercial fluorophores. If *N* fluorophores are chosen for each of the *k* excitation lasers, the total number of proteins that can be imaged simultaneously is *k* × *NN*. Higher multiplexing can be achieved by adding large Stokes shift fluorophores to these fluorophores. To stain such a multitude of proteins with antibodies, not limited by the availability of host species, we used the primary antibody–Fab complex preformation technique, which enables the use of multiple primary antibodies from the same host species^19^. Each primary antibody is assembled with a Fab fragment of a secondary antibody bearing one of those fluorophores, and then all the assembled antibody complexes are applied together to specimens (**Fig. 1a**). We validated the staining pattern (**Supplementary Fig. 5**), the crosstalks between antibodies (**Supplementary Fig. 6**), the compatibility of antibodies (**Supplementary Fig. 7** and **Supplementary Table 1**), the compatibility of fluorophores **(Supplementary Table 2**), and the staining depth (**Supplementary Fig. 8**) of the antibody preformation technique. Then, images of specimens are acquired at different spectral ranges, each containing the emission peak of each fluorophore (**Fig. 1b**). Finally, the images are unmixed *via* PICASSO and separated into images of a single protein (**Fig. 1c**). Through this approach with 15 fluorophores (**Fig. 1d**; see **Supplementary Fig. 9** for the excitation spectrum of the fluorophores), we demonstrated the 15-colour multiplexed imaging of proteins, including cell-type markers, subcellular structures, and synaptic proteins in the mouse brain, as shown in **Fig. 1e–t**. PICASSO can blindly unmix two spectrally overlapping fluorophores with only an 8-nm separation between their emission peaks (**Supplementary Fig. 10**). We note that this unmixing scheme can be extended to more than 20-colour multiplexed imaging with commercially available fluorophores.

**Figure 1.**
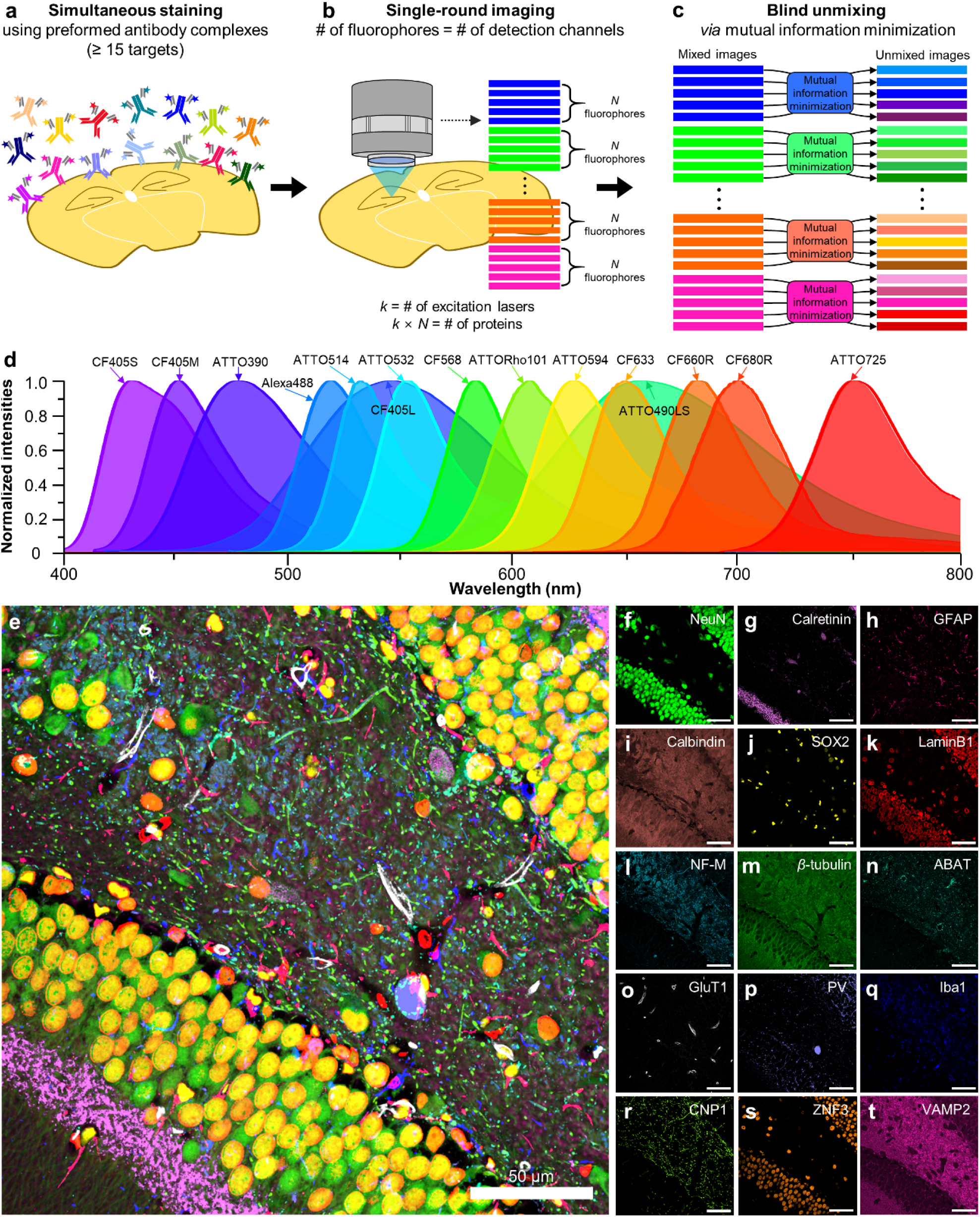
Schematic of the PICASSO process and 15-colour multiplexed imaging of the mouse brain *via* PICASSO. (**a–c**) Experimental process of multiplexed immunofluorescence imaging *via* PICASSO. (**a**) Simultaneous immunostaining of more than 15 proteins with primary antibody–Fab complexes conjugated with spectrally overlapping fluorophores. (**b**) Acquisition of images at different detection channels. The number of required images equals the number of fluorophores to be unmixed. For each of the *k* excitation lasers, *N* spectrally overlapping fluorophores are used, making the total number of fluorophores *kk* × *NN*. (**c**) Blind source separation of *N* mixed images into *N* images, each of which contains the signal of only one fluorophore through progressive mutual information minimization. (**d–t**) Fifteen-colour multiplexed imaging. Fifteen images were acquired at 15 detection channels and blindly unmixed *via* PICASSO. (**d**) Emission spectra of the 15 fluorophores used. (**e**) Fifteen-colour multiplexed imaging of the dentate gyrus of the mouse hippocampus. Target proteins are listed in (**f–t**). (**f–t**) Single-channel images of **e**. In **e**, the contrast of each channel was adjusted to clearly show all channels. In **f–t**, images were displayed without any contrast adjustments or thresholding.

### General working principle of PICASSO

The PICASSO unmixing algorithm takes *N*-channel mixed images of *N*-fluorophores as the input and obtains the unmixed images by iteratively subtracting scaled images from one another to minimize the mutual information (MI) between the images. The main assumption underlying PICASSO is that spectral mixing results in an increase of the MI between multiple channels, and therefore the unmixed images can be recovered *via* MI minimization. We first confirmed this assumption in a simple setting, with two spectrally overlapping fluorophores and two specific detection channels. We set two detection channels, such that the first detection channel contained the signal of only the first fluorophore, while the second detection channel contained the signal of both fluorophores. Therefore, we can express the relationship between the images as follows:

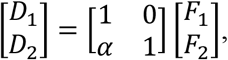

where *D*_1_ and *D*_2_ are the first and second images acquired in the first and the second detection channels, respectively; *F*_1_ and *F*_2_ are the images of the first and second fluorophores, respectively; and α is the ratio of the fluorescence intensity of the first fluorophore in the second detection channel to the fluorescence intensity of the first fluorophore in the first detection channel (**Supplementary Fig. 11**). Then, the ground-truth image of the second fluorophore (*F*_2_) can be obtained by simply subtracting the first image (*D*_1_) from the second image (*D*_2_) after multiplying the first image by α (i.e. *F*_2_ = *D*_2_ − α *D*_1_). We confirmed through both experiments and simulations (see **Supplementary Note 2** and **Supplementary Figs. 12–16** for details) that α can be accurately and robustly calculated by solving a simple optimization problem as follows:

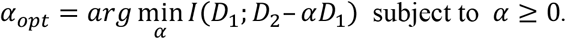

The PICASSO unmixing algorithm is a generalized version of this algorithm that can handle an arbitrary number of channels and has relaxed constraints on the acquired images. For *N*-channel unmixing, we can express the relationships among the images as follows:

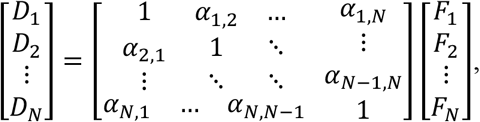

where α_*i,jj*_ refers to the relative leakage from the j_th_ fluorophore to the ith image. To undo the mixing, PICASSO starts by initializing the ith channel solution *X*_*i*(0)_ as *D*_*i*_ and progressively updates the solution based on the following equation:

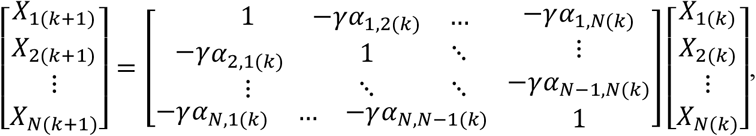

where the *k, γ*, and *X*_*i*(*k*)_ denote the iteration number, the update step size, and the i_th_channel image after k iterations, respectively, and α_*i,jj*(*k*)_ is calculated as 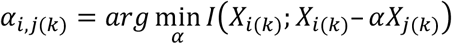. Conceptually, these iterative updates can be explained as follows. PICASSO starts by selecting a pair from multi-channel images and estimates the level of mixing between the two channels by measuring the MI. Based on the level of mixing, we subtract the scaled images from other images and repeat this process for all possible pairs. This procedure is iteratively performed to progressively minimize the MI (**Fig. 2a**; see **Supplementary Note 3** for a detailed description). A sufficient condition for PICASSO is that first, each fluorophore appears brightest in one channel compared to other channels (not compared to other fluorophores), and second, the determinants of the sub 2 × 2 matrices of the mixing matrix are positive.

**Figure 2.**
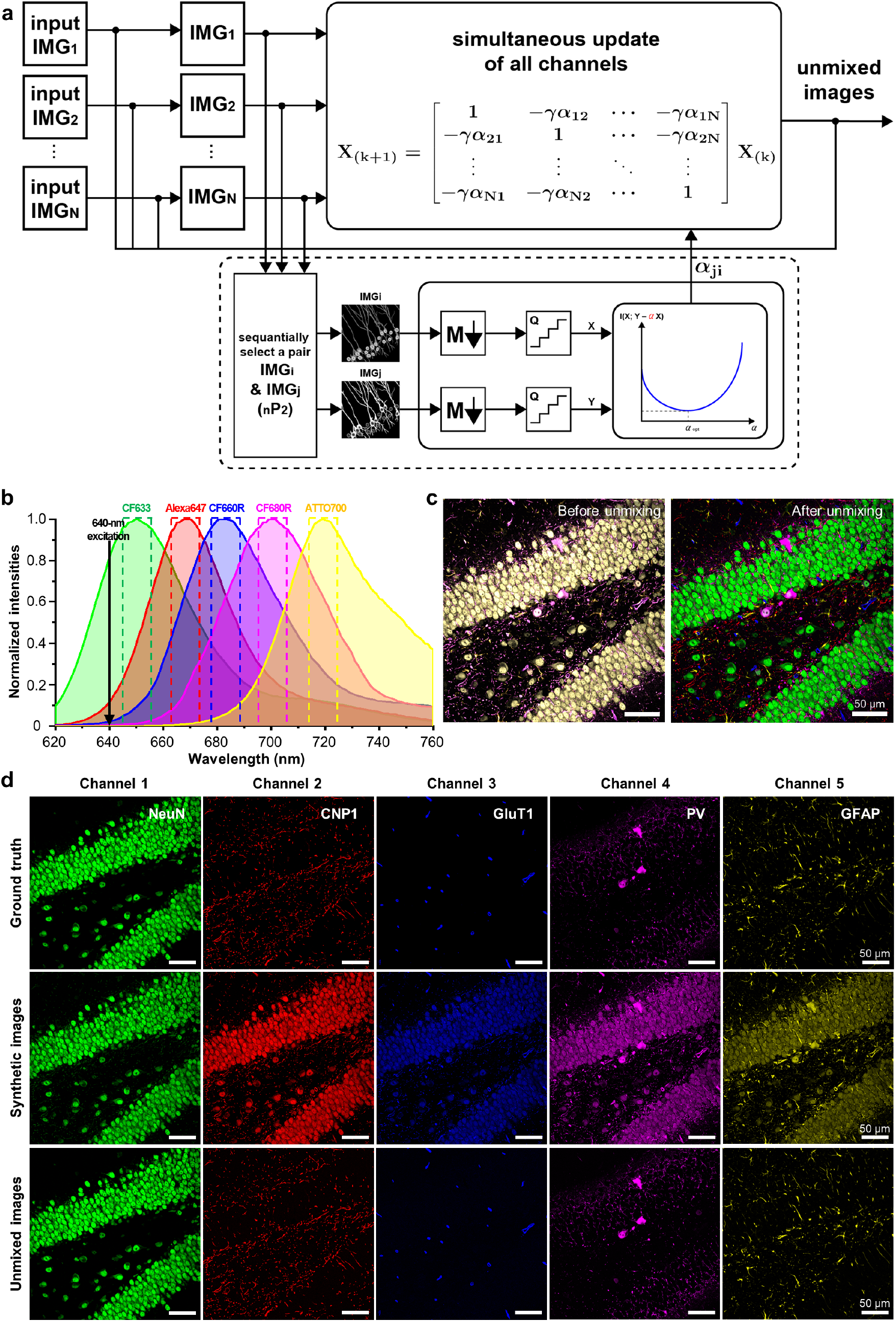
General working principle and simulation result of five-colour unmixing *via* PICASSO. (**a**) Schematic of PICASSO unmixing algorithm. The images are unmixed by progressively minimizing the mutual information between mixed images. (**b**) Emission spectra of five fluorophores and detection channels used in the simulation. Solid line: emission spectrum of each fluorophore. Dotted box: detection channel defined from -5 to +5 nm of the emission maximum of each fluorophore. (**c**) Result of five-colour unmixing simulation. Left: Overlay of the five synthetic mixed images. Synthetic mixed images were generated by mixing five single-channel images according to the mixing matrix calculated from the emission spectra of the fluorophores and the detection channels shown in **b**. Right: Overlay of five images after unmixing *via* PICASSO. (**d**) Single-channel images before mixing, which are ground-truth images (1^st^ row), synthetic mixed images (2^nd^ row), and unmixed images (3^rd^ row).

We verified the capability of the algorithm by performing a five-colour unmixing simulation. We first chose five spectrally overlapping fluorophores (CF633, Alexa Fluor 647, CF660R, CF680R, and ATTO700) that can be simultaneously excited using a 640-nm laser. Then, the mixing matrix was generated by calculating the contribution of the fluorophores in five detection channels, each of which was defined from -5 to +5 nm of the emission maximum of each fluorophore (**Fig. 2b**). Ten-nm-wide detection channels centered at the emission maximum of each fluorophore were used, but broader, narrower, or shifted detection channels would also work. The synthetic mixed images were generated by mixing a five-colour source image of a mouse brain slice, in which each channel corresponded to neuronal nuclei (NeuN), 2’, 3’-cyclic-nucleotide 3’-phosphodiesterase (CNP1), glucose transporter 1 (GluT1), parvalbumin (PV), and glial fibrillary acidic protein (GFAP) with Gaussian additive noise (**Fig. 2c**). Then, the synthetic images were fed to the unmixing algorithm to obtain the unmixed images. The unmixed images were identical to the source ground-truth images (**Fig. 2d**) with an average Pearson correlation coefficient of 0.99 despite the high level of spectral overlap, verifying the blind unmixing capability of the PICASSO algorithm.

We also found that PICASSO could unmix different fluorophore combinations even when the same detection channels were used. We used the five detection channels shown in **Fig. 2b** and unmixed 32 different fluorophore combinations of 10 spectrally overlapping fluorophores. For all 32 tested combinations, PICASSO reliably unmixed the mixed images with the Pearson correlation of unmixing around 0.99 (**Supplementary Table 3**). This result indicates that multiple different fluorophore combinations can be unmixed with fixed detection channels, without the optimization of detection channels for each fluorophore combination. This result also suggests that the same detection channel could be used to unmix given fluorophores in diverse conditions, in which the emission spectra of the fluorophores could shift a few nanometers due to the environmental or chemical factors described in **Supplementary Note 1**. Due to the flexibility of PICASSO, it could be implemented with microscopy equipped with bandpass filters with fixed detection channels to unmix various fluorophore combinations, or to unmix given fluorophores in diverse conditions. We demonstrate the PICASSO unmixing with bandpass filters in the latter part of the manuscript. Also, for all 32 combinations, the determinants of their 2 × 2 matrices were positive (**Supplementary Table 3**). We further validated the performance of PICASSO by unmixing five highly overlapping fluorophores (CF488A, ATTO488, ATTO514, Alexa Fluor 514, and CF532) with only an 8-to 10-nm difference in their emission maxima. PICASSO successfully unmixed these highly overlapping fluorophores with a Pearson correlation coefficient of 0.99 (**Supplementary Fig. 17**).

Next, we experimentally validated this algorithm. A mouse brain slice was stained with three antibody complexes against PV, NeuN, and GFAP, conjugated with CF488A, ATTO514, and ATTO532, respectively. At the same time, the mouse brain slice was stained with two antibodies against NeuN and GFAP, conjugated with CF660R and CF405S, respectively. By using a 488-nm excitation laser, we acquired three mixed images of PV, NeuN, and GFAP at three detection ranges (**Fig. 3a, b**). The images were fed to our blind unmixing algorithm to obtain the unmixed images (**Fig. 3c, d**). For validation, we acquired ground-truth images of NeuN and GFAP by using 405-nm and 640-nm lasers (**Fig. 3e**), and the unmixed images showed excellent agreement with the ground-truth images.

**Figure 3.**
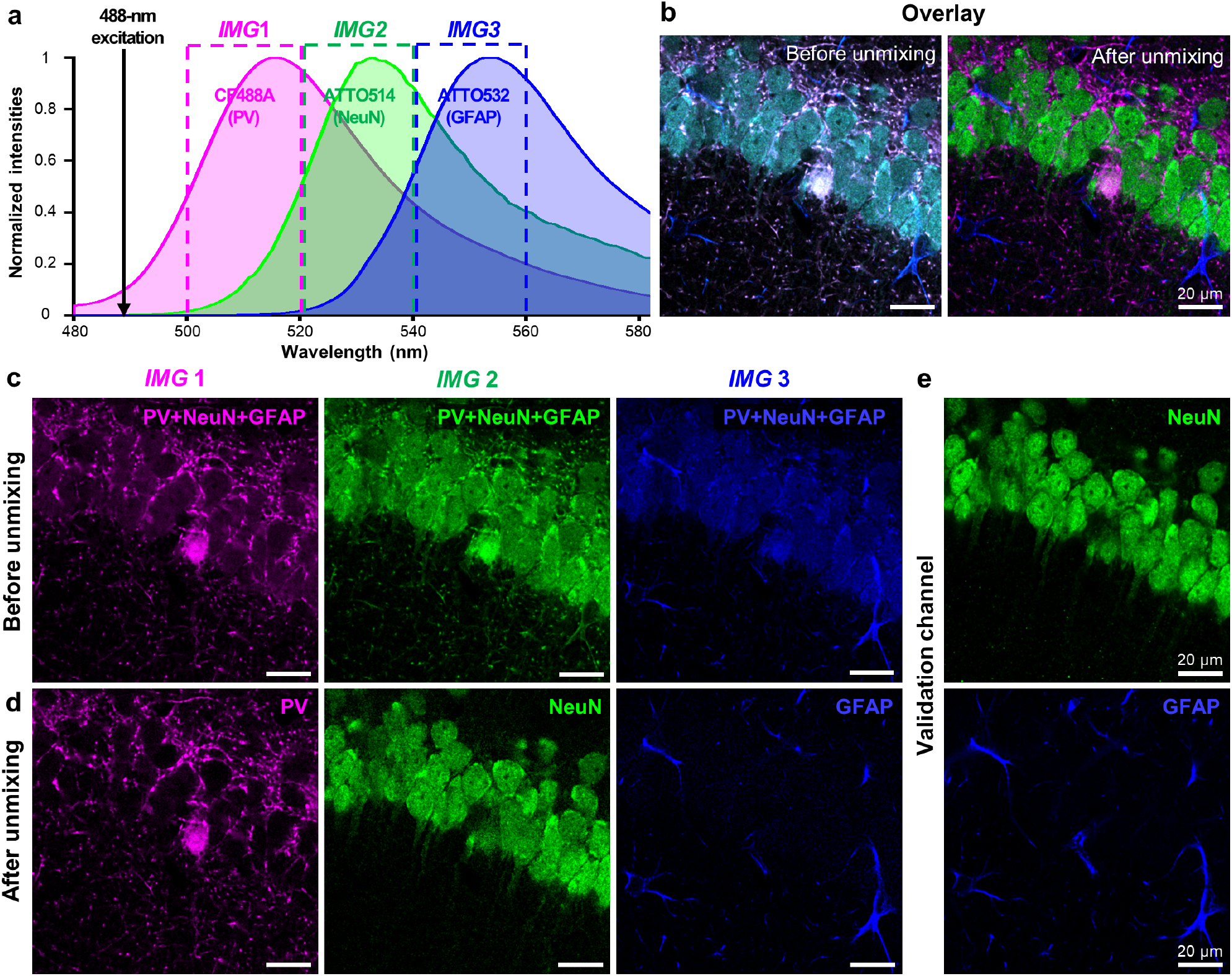
Experimental validation of the unmixing *via* PICASSO. (**a**) Emission spectra of three fluorophores and their target proteins for the validation experiment. PV was labelled with CF488A, NeuN was labelled with ATTO514, and GFAP was labelled with ATTO532. Solid line and shaded area: emission spectrum of each fluorophore. Dotted box: detection channel where images were acquired. (**b**) Overlays of the three-channel images before and after unmixing *via* PICASSO. Left: before unmixing. Right: after unmixing. (**c**) Three images acquired at the three detection channels shown in **a**. (**d**) Three images after unmixing. Each image contains the signal of only one protein. (**e**) Fluorescence images of the validation channels acquired through the simultaneous staining of NeuN and GFAP of the same specimen with other antibodies labelled with spectrally distinctive fluorophores. All scale bars: 20 μm.

### Demonstration of PICASSO with multiple excitation lasers

We then implemented PICASSO with multiple excitation lasers. Eight rabbit antibody complexes against PV, calbindin, NeuN, GFAP, GluT1, zinc finger protein 3 (ZNF3), laminin, and calretinin were simultaneously applied to a mouse brain slice (see **Supplementary Video 1** for the sample preparation procedure). An eight-colour mosaic image was acquired over a millimeter with a lateral resolution of 250 nm and then unmixed *via* PICASSO (**Fig. 4a**; see **Supplementary Fig. 18** for an enlarged image). The resulting image showed heterogeneous protein expression in the thalamus, as shown in **Fig. 4b**. For example, when the protein expression levels of the two cells in the dotted box in **Fig. 4b** were compared; the left cell highly expressed calbindin, calretinin, and ZNF3; by contrast, the right cell highly expressed NeuN and laminin (**Fig. 4c–h**). The eight-colour imaging also showed the cellular organization of the blood-brain barrier (BBB) from the same specimen (**Fig. 4i–m**). As previously reported^20^, endothelial cells were surrounded by a basal lamina and then wrapped by the end-feet of astrocytes in the BBB (**Fig. 4i**). We compared the spatial expression patterns of five proteins (calbindin, calretinin, ZNF3, NeuN, and PV) with literature reports^21–24^ and databases (rows 1–4 of **Supplementary Table 4**), and the protein expression patterns we observed matched these reports (see **Supplementary Note 4** for details and **Supplementary Fig. 19** for the individual images of these five proteins).

**Figure 4.**
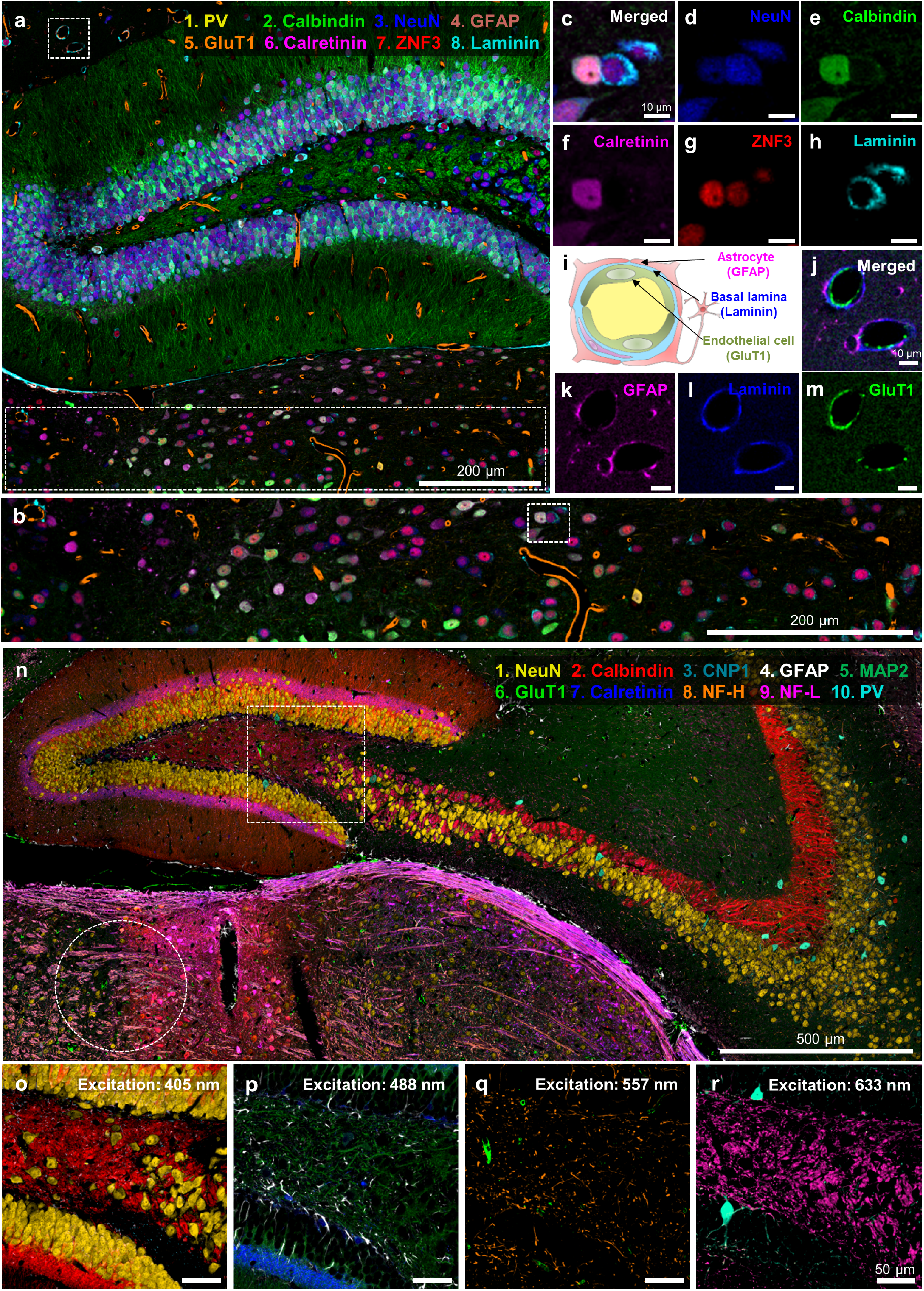
Multiplexed imaging of the mouse brain *via* PICASSO with multiple excitation lasers. (**a**) Eight-colour multiplexed imaging of the dentate gyrus of the mouse hippocampus. (**b**) Magnified view of the dotted boxed region at the bottom of **a**. (**c–h**) Magnified view of the dotted boxed region in **b** for individual labelled proteins. (**i**) Structure of the blood-brain barrier (BBB). (**j–m**) Magnified view of the dotted boxed region at the top left of **a** clearly showing the cellular structures of the BBB. (**n**) Ten-colour multiplexed imaging of the mouse hippocampus. (**o–r**) Magnified view of the dotted boxed region of **n**. Overlays of two or three channels acquired with a single excitation laser.

Next, we attempted 10-colour multiplexed imaging. A mouse brain slice was stained with 10 preformed antibody complexes against NeuN, CNP1, calbindin, calretinin, GFAP, microtubule-associated protein 2 (MAP2), neurofilament-H (NF-H), GluT1, neurofilament-L (NF-L), and PV. Then, 10 images were acquired at 10 detection channels and unmixed *via* MI minimization. As a result, the 10 proteins were successfully visualized over a few millimeters (**Fig. 4n–r**; see **Supplementary Fig. 20** for an enlarged image). The lateral posterior nucleus of the thalamus (the region in the right half of the dotted circle in **Fig. 4n**) showed high calbindin and calretinin expression (see **Supplementary Fig. 21** for details), matching the results reported in the various databases (row 1 of **Supplementary Table 4**) and the literature^22^. The use of PICASSO is not limited to multiplexed immunostaining; we showed that PICASSO could be applied to multiplexed mRNA fluorescence in-situ hybridization (FISH) (**Supplementary Fig. 22a–r**), the simultaneous imaging of proteins and mRNA (**Supplementary Fig. 22s–v**), the super-resolution imaging of tissue *via* physical expansion^25^ (**Supplementary Fig. 23a, b**), and tissue clearing^26^ (**Supplementary Fig. 23c, d**) to improve their multiplexing capability.

We attempted three demonstrations of PICASSO, which possibly maximized its potential. First, PICASSO enabled the 3D multiplexed imaging of a thick mouse brain slice (**Fig. 5a**) (see **Supplementary Video 2** for 3D video; see **Supplementary Fig. 24a** for individual channels; see **Supplementary Fig. 24b, c** for more 3D imaging demonstrations). Second, PICASSO successfully unmixed the signals of spectrally overlapping fluorescent proteins and organic fluorophores (**Supplementary Fig. 25**). Lastly, 10-colour multiplexed imaging *via* PICASSO was achieved with a microscope equipped with emission bandpass filters, which did not have a spectral imaging capability (**Fig. 5b**). Microscopy systems equipped with spectral detectors are still not commonly available in most research laboratories and hospitals; instead, most microscopy systems use bandpass filters to detect fluorescence signals in specific spectral ranges. Like most bandpass filter**-**based microscopes, our equipment was able to accommodate up to eight filters, so we used two filters twice for 10-colour imaging (see **Supplementary Table 5** for details of the filters and imaging conditions). A mouse brain slice was stained with 10 antibody complexes and imaged with microscopy equipped with eight bandpass filters, and the signals were unmixed *via* PICASSO (**Fig. 5b–f**). By using this instrumental setting, a 10-colour multiplexed image with a lateral resolution of 250 nm over a field of view of 320 × 320 μm was acquired in less than six seconds (with a 40× NA1.15 objective; see **Supplementary Video 3**). At this speed, the imaging of a 1-mm^2^ field of view would take approximately one minute. Five or six channels were selected and overlaid to better visualize the biological contexts (**Fig. 5g–i**).

**Figure 5.**
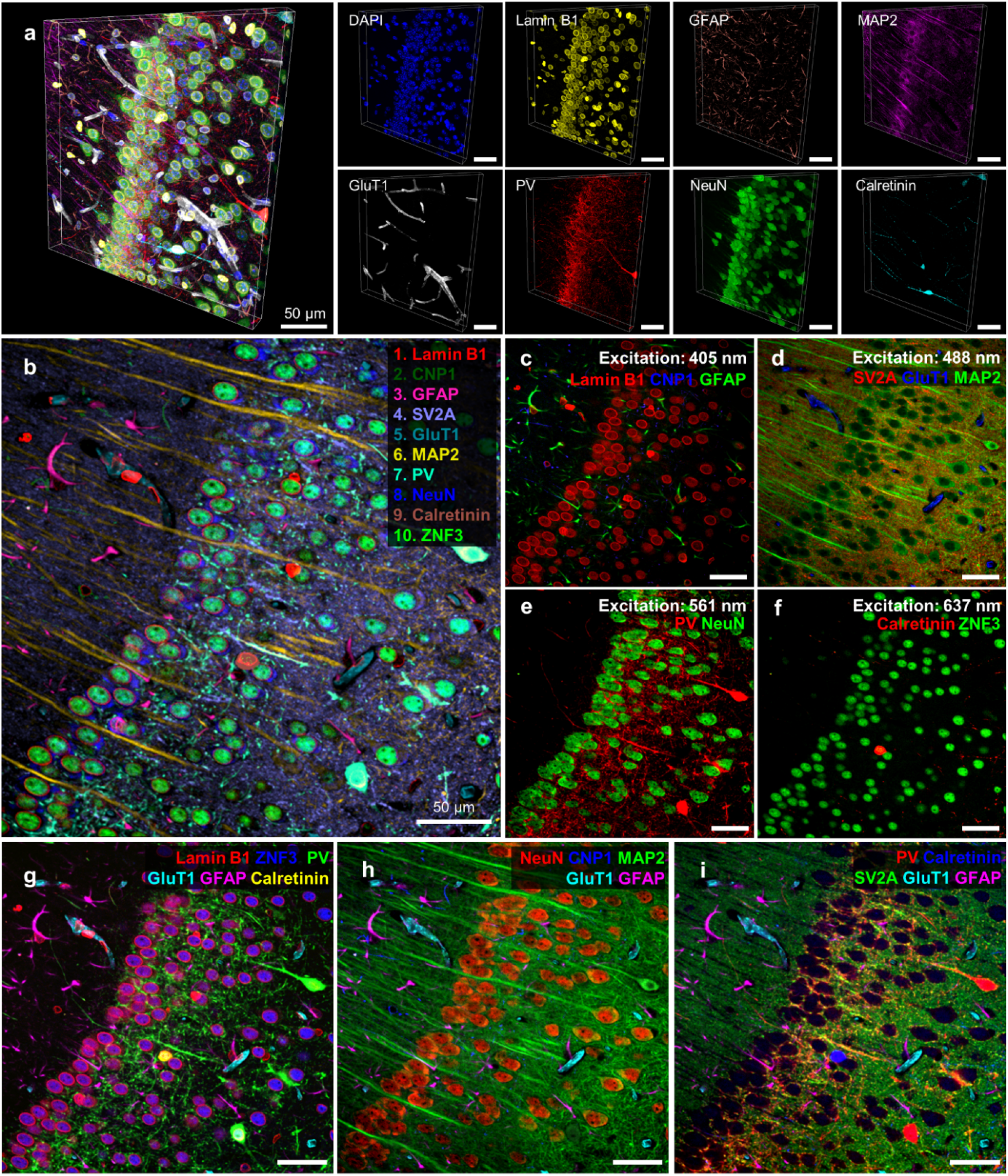
Three-dimensional multiplexed imaging *via* PICASSO and the demonstration of PICASSO with microscopy equipped with bandpass filters. (**a**) Three-dimensional view of a z-stack image acquired from a mouse brain slice stained, imaged, and unmixed *via* PICASSO. Left: Overlay of the eight channels. Right: Single-channel images. (**b**) Demonstration of the 10-colour multiplexed imaging of the mouse hippocampus CA1 *via* PICASSO with microscopy equipped with bandpass filters. The images were acquired using a confocal microscope equipped with eight bandpass filters and four excitation lasers. (**c–f**) Two or three channels of the image shown in **b** acquired by one of the four excitation lasers. The wavelengths of the excitation lasers were (**c**) 405 nm, (**d**) 488 nm, (**e**) 561 nm, and (**f**) 637 nm. (**g–i**) Overlay of five or six channels chosen from the image shown in **b** to clearly visualize different sets of biologically relevant channels together. Different display colours were used for proteins in each image to maximize visibility. (**a–i**) All primary antibodies were rabbit antibodies.

We tested whether PICASSO worked with clinical samples using a tissue microarray containing FFPE specimens from 12 human organs. We chose keratin 19, histone H3, vimentin, and cytochrome c oxidase subunit 4 (COX IV) as target proteins, as they are related to cancers and used as cancer markers^27–30^. We first attempted to image keratin 19 and histone H3 with two spectrally overlapping fluorophores. Keratin 19 and histone H3 are localized in different cellular compartments (in the cytoplasm and nucleoplasm, respectively; rows 5 and 6 of **Supplementary Table 4**); their expression patterns should be spatially separated if their signals are successfully unmixed. The obtained images clearly show the spatial separation of the two proteins (**Fig. 6a–l**), indicating that PICASSO successfully separated the signals of two spectrally overlapping fluorophores in clinical specimens. We then attempted the simultaneous imaging of keratin 19, histone H3, vimentin, and COX IV in clinical specimens. The simultaneous visualization of keratin and vimentin in a single specimen is also clinically important, as the vimenin:keratin ratio is related to the epithelial–mesenchymal transition status^31^. The four proteins were successfully visualized using two pairs of spectrally overlapping fluorophores and two excitation lasers in four different organs (**Fig. 6m–p**). In the imaging of clinical specimens, autofluorescence is sometimes a severe problem. To remove the autofluorescence from immunostaining signals, the emission profiles of autofluorescence need to be measured for each specimen before the staining and then used as a reference profile for unmixing. PICASSO was able to separate the autofluorescence from true staining signals without any autofluorescence measurement (**Supplementary Fig. 26**).

**Figure 6.**
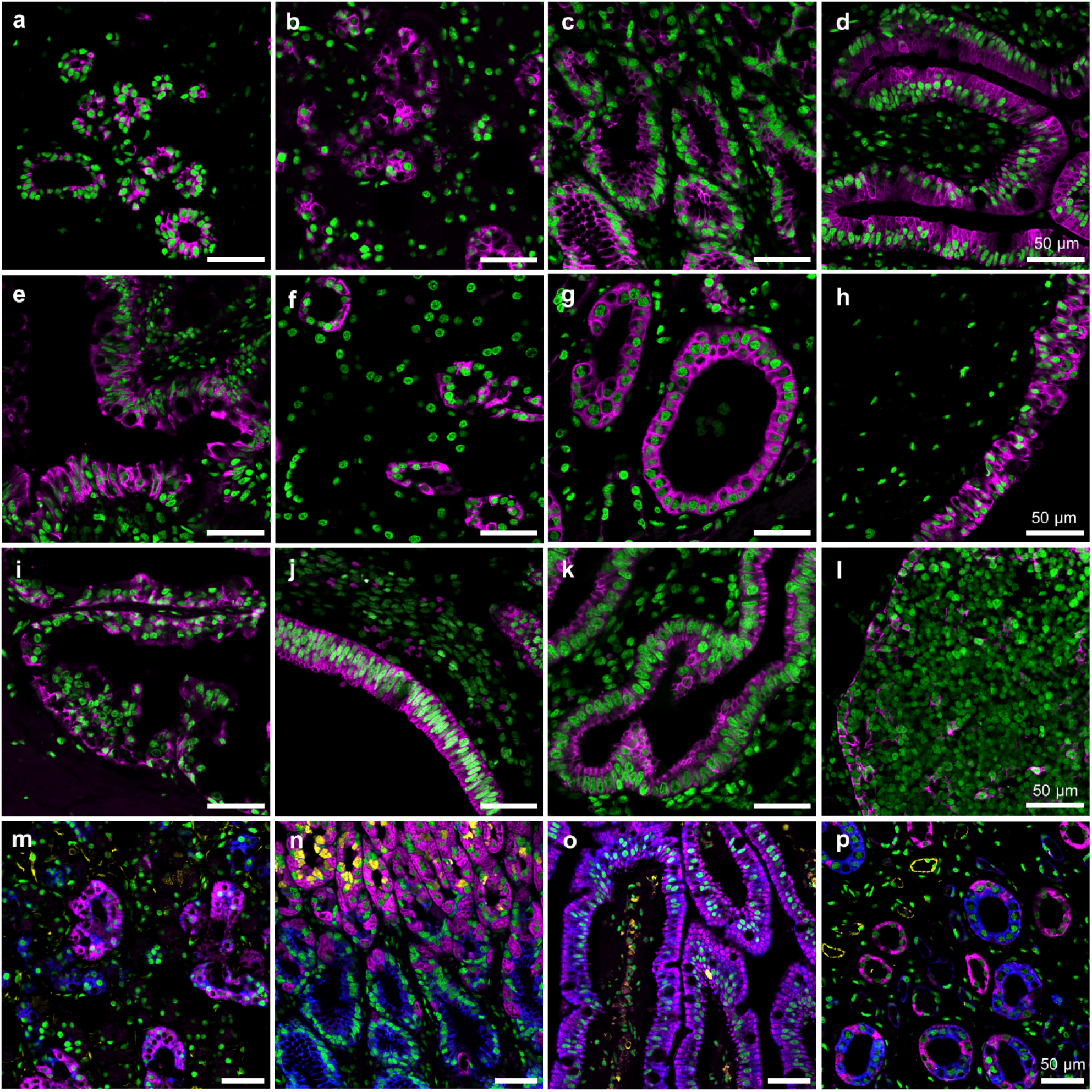
Multiplexed imaging of FFPE clinical samples *via* PICASSO. (**a–l**) Two-colour multiplexed imaging of 12 different human tissue types using one excitation laser enabled *via* PICASSO. Magenta, keratin 19 (CF488A); green, histone H3 (ATTO514). (**a**) Breast. (**b**) Salivary gland. (**c**) Stomach, body. (**d**) Small intestine, jejunum. (**e**) Colon. (**f**) Kidney, cortex. (**g**) Kidney, medulla. (**h**) Bladder. (**i**) Prostate. (**j**) Endometrium, proliferative. (**k**) Endometrium, secretory. (**l**) Thymus. (**m–p**) Four-colour multiplexed imaging of four of the same human tissue types using two excitation lasers enabled *via* PICASSO. Blue, keratin 19 (CF488A); green, histone H3 (ATTO514); magenta, COX IV (CF568); yellow, vimentin (ATTORho101). (**m**) Salivary gland. (**n**) Stomach, body. (**o**) Small intestine, jejunum. (**p**) Kidney, medulla. (**a–p**) All primary antibodies were rabbit antibodies.

Finally, we demonstrated 15-colour multiplexed imaging of the mouse hippocampus, where the expressions of these 15 proteins spatially overlapped, as shown in **Fig. 1e** (see **Supplementary Fig. 27** for more 15-colour imaging demonstrations). In this experiment, three spectrally overlapping fluorophores were used for each of 405-, 488-, 557-, and 640-nm excitation lasers. The fluorophores used were CF405S, CF405M, and ATTO390 for a 405-nm laser, Alexa Fluor 488, ATTO514, and ATTO532 for a 488 nm laser, CF568, ATTORho101, and ATTO594 for a 557-nm laser, and CF633, CF660R, and CF680R for a 640-nm laser. For the 405- and 488-nm lasers, large Stokes shift fluorophores (CF405L and ATTO490LS) were used along with the spectrally overlapping fluorophores. One more infrared fluorophore (ATTO725) was used for a 730-nm excitation laser, making the total number of fluorophores 15 (see **Supplementary Table 6** for the complete list of the fluorophores). In this experiment, we used a spectral detector with variable detection ranges to acquire images at 15 detection channels. To confirm that the same 15-colour unmixing could be achieved by using a linear detector array with predetermined detection channels, we defined consecutive 10 nm-wide detection channels and chose the ones containing the emission maxima of the above fluorophores. We then generated a mixing matrix by calculating the contribution of the fluorophores in each of the chosen detection channels, generated synthetic mixed images, and then unmixed them. PICASSO successfully unmixed those synthetic mixed images, indicating that it works with a linear detector array with predetermined detection channels (data not shown). As PICASSO can reliably unmix more than five fluorophores, as shown in **Fig. 2**, the simultaneous imaging of more than 27 fluorophores would be achievable with five excitation lasers, including large Stokes shift fluorophores.

PICASSO can be combined with cyclic immunofluorescence imaging techniques to image an unlimited number of proteins with a reduced number of cycles. A brain slice was stained with antibody complexes labelled with spectrally overlapping fluorophores, imaged, chemically bleached *via* tissue-based cyclic immunofluorescence (t-CyCIF) protocol^32^, re-stained with another two antibody complexes labelled with spectrally overlapping fluorophores, and then imaged again (**Supplementary Fig. 28**). PICASSO unmixed the signals of two spectrally overlapping fluorophores in both cycles, indicating that it can be combined with t-CyCIF to increase the number of fluorophores that can be used simultaneously in one cycle.

## DISCUSSION

We herein proposed PICASSO, which enables ultra-multiplexed fluorescence imaging of biological specimens through the blind unmixing of heavily mixed fluorophores. By simulation, we showed that PICASSO can reliably unmix five spectrally overlapping fluorophores with less than 10-nm spectral separation and that the number of fluorophores would not be limited to five. We demonstrated 15-colour multiplexed imaging of proteins, whose expression patterns overlapped using multiple excitation lasers. PICASSO does not need reference spectra; hence, 15 or even more proteins can be simultaneously imaged without the time-consuming, complicated, and even sometimes impossible reference measurement processes. PICASSO needs only one image per fluorophore, minimizing the number of required detection channels. As PICASSO needs a small number of images and its requirements for detection channels are not strict, it can be implemented with bandpass filter-based microscopy, which is currently available to most biology laboratories and hospitals. The use of a small number of images also has advantages in terms of signal-to-noise ratio, as the number of photons collected per channel can be maximized for the simultaneous imaging of a given number of fluorophores within a given spectral range^7^. As PICASSO does not need repeated staining and imaging processes up to 15-colour, multiplexed 3D imaging can be easily performed without a complicated 3D registration process. We showed that PICASSO can increase the multiplexing capability of mRNA imaging, the co-imaging of mRNA and proteins, expansion microscopy, tissue clearing, and even cyclic staining. PICASSO shares the practical limitations of spectral unmixing, such as the susceptibility to noise and difficulty in distinguishing spectrally identical fluorophores. Such fluorophores could be resolved by combining PICASSO with excitation unmixing.

PICASSO is a versatile tool for multiplexed biomolecule imaging of cultured cells, tissue slices, and clinical specimens. It is suitable for high-throughput protein imaging, as antibodies can be simultaneously applied to specimens during the staining process, and only a minimum number of images need to be collected. In addition, PICASSO does not need complicated optics or spectral detectors; instead, it can be implemented with a simple imaging system consisting of an objective, light sources such as a lamp or LED, a camera, and excitation/emission bandpass filters. For more than 10-colour imaging, custom multi-band pass filters could be used for such simple microscopy setups. PICASSO can be combined with live imaging^4^, tissue clearing^26,33^, or tissue expansion^25,34–36^ techniques to achieve multiplexed 3D super-resolution imaging or multiplexed whole-organ imaging. PICASSO can be combined with many forms of cyclic immunofluorescence imaging without any significant change^2,32,37–42^. Once combined, fewer cycles are needed to image a given number of proteins, greatly reducing the time and complexity of the whole imaging process. PICASSO is a strategy for distinguishing fluorophores and thus could be used for mRNA imaging^43–46^, bioassays^47^, and cell tracing^48^ to improve their multiplexing capabilities. It could also be combined with fluorescent barcoding techniques to increase the amount of information that a single barcode can encode^49–52^.

We anticipate that PICASSO will be useful for a broad range of applications for which the spatial information of biomolecules is important. PICASSO would be useful to reveal the cellular heterogeneities of tumour microenvironments, especially the heterogeneous populations of immune cells, which are closely related to cancer prognoses^1^ and the efficacy of cancer therapies^53^. In addition, the multi-scale 3D atlases of biomolecules^54–57^ have changed our understanding of biology^58^. In constructing these atlases, multiplexed fluorescence imaging is one of the major tools to map the spatial distribution of biomolecules. PICASSO is expected to enable the 3D visualization of more biomolecules simultaneously, providing information about the co-localization of different proteins or mRNA molecules, or how their expressions are correlated.

## METHODS

### Online methods

All the chemicals used in this study are listed in **Supplementary Table 7**.

#### PICASSO

PICASSO is an acronym of ‘**P**rocess of ultra-multiplexed **I**maging of biomole**C**ules *vi****A*** the unmixing of the **S**ignals of **S**pectrally **O**verlapping fluorophores’.

#### Cell culture and fixation

BS-C-1, HeLa and NIH-3T3 cells were purchased from the Korean Cell Line Bank and cultured in a Nunc Lab-Tek II chambered #1.5 coverglass. The BS-C-1 cells were cultured in Minimum Essential Medium (MEM) supplemented with 10% fetal bovine serum (FBS), 1% penicillin-streptomycin and 1% sodium pyruvate. The HeLa cells were cultured in Dulbecco’s modified Eagles’ medium (DMEM) supplemented with 10% FBS and 1% penicillin-streptomycin. The NIH-3T3 cells were cultured in DMEM supplemented with 10% bovine calf serum (BCS) and 1% penicillin-streptomycin. All cells were incubated at 37 °C in 5% CO2. For the fixation, the cells were washed with 1× phosphate-buffered saline (PBS) three times and fixed with 4% paraformaldehyde (PFA) in 1× PBS for 10 min, then washed with 1× PBS three times. The cells were then incubated with 0.1 M glycine in 1x PBS for 10 min and washed with 1× PBS three times.

#### Mouse brain perfusion and slicing

All of the following procedures involving animals were approved by the Sungkyunkwan University Institutional Animal Care and Use Committee (SKKU-IACUC), and the Korea Advanced Institute of Science and Technology Institutional Animal Care and Use Committee (KAIST-IACUC). C57BL/6J and Thy1-YFP mice aged 8–14 weeks were used. The mice were anaesthetized with isoflurane and transcardially perfused with ice-cold 4% PFA in 1× PBS. Brains were harvested and stored in the same solution at 4 °C for 2 h before being sliced into 150 μm-thick slices on a vibratome (Leica VT1000S). The slices were stored in 0.1 M glycine and 0.01% sodium azide in 1× PBS at 4 °C until use.

#### Conjugation of Fab fragment antibodies with fluorophores

For the Alexa and CF fluorophores, 10 μL of 1 M sodium bicarbonate (pH 8.3) and a 9-fold molar excess of succinimidyl ester-fluorophore stock in dimethyl sulfoxide (DMSO) were added to 90 μL of unconjugated antibody solutions. For the ATTO fluorophores, 10 μL of 1 M sodium bicarbonate (pH 8.3) and a 3-fold molar excess of succinimidyl ester-fluorophore stock in dimethyl sulfoxide (DMSO) were added to 90 μL of unconjugated antibody solutions. The fluorophore-antibody solutions were incubated at RT for 1 h in darkness. To purify the labelled antibodies, NAP-5 gel filtration columns were used. The columns were equilibrated with 10 mL of 1× PBS. 100 μL of the reacted solutions were loaded into the columns. The eluates containing fluorophore-conjugated antibodies were collected after loading 900 μL of 1× PBS into the columns. Then, the eluates were concentrated using centrifugal filters with a molecular weight cut-off (MWCO) of 30,000.

#### Preparation of preformed antibody complexes

Preformed antibody complexes were prepared according to the original primary antibody-Fab complex formation protocol^19^. Briefly, 1× PBS, a solution containing a fluorophore-conjugated Fab fragment antibody, and a solution containing a primary antibody were mixed at a volume ratio of 10:2:1 and then incubated for 10 min at RT in darkness. A 5-fold excess volume of a blockng buffer (10% normal rabbit serum, 0.2% Triton X-100, 1× PBS) was then added to the solution and incubated for 10 min at RT in darkness with gentle shaking. Then, the preformed antibody complexes against different targets were mixed together, diluted in the blocking buffer (1:1000 to approximately 1 μg/mL) and then used for staining. The 28 fluorophores tested with the preformed antibody complex are listed in **Supplementary Table 2**.

#### Staining of cells and mouse brain slices with preformed antibody complexes

All of the following steps were performed at RT. For permeabilization and blocking, cells were incubated in a blocking buffer (10% normal rabbit serum, 0.2% Triton X-100, 1× PBS) for 30 min. The cells were then stained with a preformed antibody complex mixture for 30 min and then washed three times with the blocking buffer. For the staining of mouse brain slices, blocking, staining, and washing steps were identical to the cultured cell protocol except for the incubation time (blocking: 1.5 h, staining: overnight, washing: 30 min).

#### Validation of the staining using a preformed antibody complex

For the validation study shown in **Supplementary Fig. 5**, a mouse brain slices were blocked in a blocking buffer (10% normal rabbit serum, 0.2% Triton X-100, 1× PBS). The slice was then incubated overnight in a preformed rabbit antibody solution diluted at 1:500 at 4 °C and then washed three times with a blocking buffer. The slice was then stained overnight with a 1:500 chicken primary antibody in an NGS buffer (5% normal goat serum, 0.2% Triton X-100, 1× PBS) at 4 °C and then washed three times with the NGS buffer. The slice was then stained overnight with an anti-chicken secondary antibody diluted to 1:250 in the NGS buffer at 4 °C and then washed with the NGS buffer three times. For the staining of BS-C-1 cell, blocking, staining, and washing steps were identical to the cultured cell protocol except for the incubation time (blocking: 30 min, staining: 30 min, washing: immediately).

#### Study on the crosstalk between two preformed antibody complexes

For the study shown in **Supplementary Fig. 6a–c**, a Thy1-YFP mouse brain slice was stained with a mixture of a preformed rabbit anti-GFP antibody (not labelled) and preformed rabbit anti-neurofilament-H antibody (labelled with CF568). In **Supplementary Fig. 6d–f** another sample was stained with a mixture of a preformed rabbit anti-GFP antibody (not labelled) and preformed rabbit anti-calbindin antibody (labelled with CF633).

#### Validation of unmixing *via* MI minimization

For the validation of unmixing *via* MI minimization shown in **Fig. 3b–e**, a mouse brain slice was blocked in a blocking buffer (10% normal rabbit serum, 0.2% Triton X-100, 1× PBS) and then incubated overnight with a mixture of preformed rabbit anti-PV (labelled with CF488A), preformed rabbit anti-NeuN (labelled with ATTO514), preformed rabbit anti-GFAP (labelled with ATTO532), guinea pig anti-NeuN antibody, and mouse anti-GFAP antibody diluted to 1:1000 in the blocking buffer. The sample was washed with the blocking buffer three times and stained overnight with a CF405S-labelled anti-mouse secondary antibody, and CF660R-labelled anti-guinea pig secondary antibody diluted to 1:500 in the blocking buffer.

#### Staining of FFPE samples

The following experiments using human samples were all approved by the Korea Advanced Institute of Science and Technology Institutional Review Board (KAIST IRB). The normal human multi-organ tissue microarray (TMA) used in **Fig. 6** was purchased from Novus Biologicals (NBP2-30189). For the F**F**PE clinical samples, the sample slides were dried for 1 h in an oven at 60 °C. The slides were then deparaffinized in xylene twice for 5 min each time. For hydration, the slides were placed in a series of solutions – namely 100% ethanol (EtOH) twice, 95% EtOH, 80% EtOH, and deionized water at RT – for 3 min each. The slides used in this study were processed with a heat-induced epitope retrieval (HIER) procedure before the staining^59^. Briefly, the slides were placed in a 10 mM retrieval solution (10 mM sodium citrate, 0.05% Tween 20, pH 6.0) for 30 min at 95–100 °C and allowed to cool for 20 min in 1× PBS. The slides were blocked with a blocking buffer (10% normal rabbit serum, 0.2% Triton X-100, 1× PBS) for 3 h at RT, followed by overnight incubation with preformed antibody complexes at RT, before being washed three times with the blocking buffer at RT for 30 min each.

#### Hybridization chain reaction (HCR)

For the PICASSO imaging of two mRNA molecules shown in **Supplementary Fig. 22a–r**, HCR (Molecular Instruments) was conducted on NIH-3T3 cells as instructed by the manufacturer’s user manual. Briefly, fixed NIH-3T3 cells were permeabilized overnight with 70% EtOH and stored at –20 °C until their use. The permeabilized cells were rinsed with 2× saline sodium citrate (SSC) and incubated with an HCR probe hybridization buffer (Molecular Instruments) for 30 min at 37 °C. Then, the cells were incubated with a probe solution (HCR probes diluted to 4 nM in the HCR probe hybridization buffer) for 12–16 h at 37 °C. After the probe hybridization, the cells were washed four times with an HCR probe wash buffer (Molecular Instruments) for 5 min at 37 °C and then washed two more times with 5× SSCT (5× SSC, 0.1% Tween 20) for 5 min at RT. The cells were incubated with an HCR amplification buffer (Molecular Instruments) for 30 min at RT and then with an amplification solution for 12–16 h at RT in darkness. The amplification solution was prepared by the snap-cooling of HCR hairpin amplifiers H1 and H2, heating the amplifier hairpins at 95 °C for 90 sec and then cooling them down to RT in darkness for 30 min. The snap-cooled hairpins were then diluted to 6 nM in the HCR amplification buffer to make the amplification solution. After the amplification, the cells were washed five times with 5× SSCT for 5 min at RT. The cells were then stained with 4′,6-diamidino-2-phenylindole (DAPI), rinsed with 1× PBS.

#### Custom dye conjugation to HCR hairpin amplifiers

The unconjugated hairpin amplifier was purchased from Molecular Instruments. First, 75 μL of the labelling buffer (0.1 M sodium tetraborate, pH 8.5) and 5 μL of deionized water were added to a reaction vial. Then, 10 μL of 100 μM unlabelled hairpins (H1 and H2 were conjugated separately) and 10 μL of 10 mg/mL ATTORho101-NHS-ester were added to the reaction vial and allowed to react for 6 h. After the reaction, 10 μL of 3 M sodium chloride (NaCl) and 250 μL of –20 °C pure ethyl alcohol were added to the reaction vial and then stored at –20 °C for 30 min. The reaction vial was then centrifuged at 12,000 g for 30 min. After the centrifugation, the supernatant was carefully removed, and the DNA pallet on the bottom of the vial was rinsed with ice-cold 70% ethyl alcohol twice. The pellet was then dried in darkness overnight and re-dissolved in 1× PBS or storage buffer (150 mM NaCl and 100 mM sodium phosphate dibasic).

#### Simultaneous imaging of mRNA and protein

For the simultaneous imaging of the mRNA and protein shown in **Supplementary Fig. 22s–v**, RNAscope (Advanced Cell Diagnostics) was conducted on NIH-3T3 cells as instructed by the manufacturer’s user manual. After the RNAscope labelling of the *Gapdh* mRNA of the cultured cells, the cells were stained with a preformed rabbit anti-vimentin complex as described in ‘Staining of cells and mouse brain slices with preformed antibody complexes.’

#### Fluorophore inactivation (t-CyCIF)

For the first staining round, two preformed antibody complexes – one against PV (labelled with CF488A) and the other against NeuN (labelled with ATTO514) – were used to stain a 150 μm-thick mouse brain slice. DAPI was applied to the slice as a fiducial marker. After the first round of imaging, the fluorophores of the stained sample were inactivated in an inactivation buffer (4.5% H2O2 and 24 mM NaOH in 1× PBS) for 2 h at RT under a white light, according to the original t-CyCIF protocol^32^. The slice was then washed four times with 1× PBS for 1 h each, followed by a second round of staining, during which the sample was labelled with two preformed antibody complexes: one against calretinin (labelled with CF488A) and the other against GluT1 (labelled with ATTO514). DAPI was applied again after the second antibody staining cycle.

#### Protein-retention expansion microscopy

Stained brain slices were incubated in acryloyl-X, SE ((6-((acryloyl)amino)hexanoic acid, succinimidyl ester, AcX) diluted to 0.1 mg/ml in 1× PBS overnight at RT and then washed three times for 30 min with 1× PBS. The samples were then incubated twice with a monomer solution (7.5% (w/w) sodium acrylate, 2.5% (w/w) acrylamide, 0.15% (w/w) N,N′-methylenebisacrylamide (BIS), 1× PBS, 2 M NaCl) at 4 °C for 30 min each time. After incubation, the samples were placed between two coverglasses filled with a gelation buffer (monomer solution, 0.2% (w/w) ammonium persulfate (APS), 0.2% (w/w) tetramethylethylenediamine (TEMED), 0.01% (w/w) 4-hydroxy-2,2,6,6-tetramethylpiperidin-1-oxyl (H-TEMPO)) and incubated at 37 °C for 1.5 h. The gels were treated with proteinase K diluted at 1:100 in a digestion buffer (25 mM ethylendiaminetetraacetic acid (EDTA), 50 mM Tris-HCl (pH 8), 0.5% Triton X-100, 1 M NaCl) at 37 °C overnight with gentle shaking. After digestion, the digested gels were placed in deionized water (DI) with gentle shaking.

#### SHIELD

SHIELD (LifeCanvas Technologies) was conducted as instructed by the manufacturer’s user manual. Briefly, 500 μm-thick mouse brain slices were incubated in a SHIELD-OFF solution (a mixture of DI water, SHIELD-Buffer solution, and SHIELD-Epoxy solution at a ratio of 1:1:2) at 4 °C with gentle shaking for 1 day. The samples were transferred to a mixture of SHIELD-ON buffer and SHIELD-Epoxy solution with a ratio of 7:1 and then incubated at 37 °C with gentle shaking for 6 h. Samples were incubated in the SHIELD-ON buffer at 37 °C with gentle shaking overnight. For tissue clearing, samples were incubated in a clearing solution (300 mM sodium dodecyl sulfate, 10 mM boric acid, 100 mM sodium sulfite, pH 9.0) at 37 °C with gentle shaking for 1 day. Cleared samples were imaged in 1× PBS.

#### Imaging

In this work, three microscopy systems were used. The first two were point-scanning confocal microscopy systems: Leica SP8 equipped with a white light laser and Nikon C2 Plus equipped with four excitation lasers (405, 488, 557, 640 nm). Even though the Leica microscope had a white light laser and its excitation wavelength was adjustable, we used the same excitation wavelengths as the Nikon system to test the multiplexed imaging ability of PICASSO with two different typical microscopy systems. For the images shown in **Fig. 4**, 633 nm or 660 nm was used as an excitation wavelength instead of 640 nm to image red fluorophores with a higher SNR; however, 640 nm can be used to image two proteins labelled with spectrally overlapping fluorophores, as shown in **Fig. 1e–t**. The third system was a spinning-disk confocal microscopy system, which was Andor Dragonfly equipped with four excitation lasers (405, 488, 561, 637 nm). All the images were acquired by using a 40× 1.15 NA water-immersion objective. Detailed information about the imaging is listed in **Supplementary Table 6**.

#### Pre-processing of images for unmixing

We performed pre-processing of images before unmixing the images. We first resized the input images to 50% of their images by using the ‘resize’ function of the Fiji image-processing software, with the option ‘average when downsizing’ checked. The resizing might have made the estimation of α more accurate by averaging out the shot noises in the nearby pixels of the input images. For mosaic images, the shading and background variations were corrected using BaSiC^60^ and then stitched. The images from the spinning-disk confocal microscopy were further processed. First, vignetting in the images was corrected using the image of a fluorescent slide. Second, ring-shaped artefacts in the images, possibly due to the interference of the excitation laser with internal optics of the spinning-disk microscopy system (e.g. two rotating disks with an array of microlenses), were removed by subtracting an image taken without specimens. Third, the pixel shift due to the chromatic aberration was handled *via* image registration.

#### Unmixing

For PICASSO unmixing, we first initialized the solution *X*_(0)_ as the acquired images *D*. Then, we construct a progressive unmixing matrix as follows:

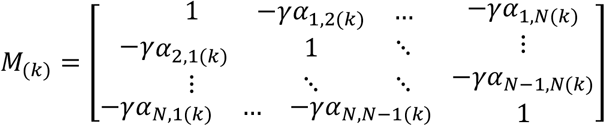

where the *k* and *γ* denote the iteration number and the update step size, respectively; α _*i,jj*(*k*)_ was calculated as 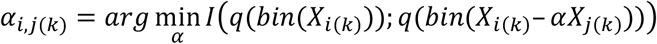 using the Nelder-Mead simplex algorithm where *q* (.)and bin (.) denote the quantization and pixel binning functions. The parameters α _*i,jj*(*k*)_ were calculated using binned and quantized images, to improve the robustness against noise and the speed of the joint histogram calculation, whereas the progressive unmixing, *X*_(*k+1*)= *M*_(*k*)_*X*(*k*)_, was performed on the full-resolution images without quantization (see **Supplementary Note 3** for details).

For linear unmixing with reference spectra, the linear matrix equation *D* = *M* × *F* was directly set using the acquired 32-channel image *D* and the reference spectra matrix *M*. The equation was solved using QR factorization. For blind unmixing with NMF, the acquired 32-channel image *D* ∈ ℝ^32×*m*^ was unmixed by directly factorizing as *D* = *W* × *H* (*W* ∈ ℝ^32×*k*^, *H* ∈ ℝ^*k*×*m*^), where *m* is the number of pixels of the image and *k* is the number of fluorophores, using NMF with alternating least squares method. All image processing was conducted using custom-written MATLAB codes.

### Data availability

The data that support the findings of this study are available from the corresponding author upon request.

## ACKNOWLEDGEMENTS

This work was supported by the Samsung Research Funding & Incubation Center for Future Technology (SRFC-IT1702-09). Spectral imaging by using a Leica SP8 microscope was performed in the Korea Basic Science Institute (KBSI) Western Seoul Center. We acknowledge Uhtaek Oh and Jinhyun Kim for their helpful discussions on brain imaging. We acknowledge Seok-Gyu Kwon and Hyunsu Jung for their comments on the preparation of mouse brains. We also acknowledge Seong Gi Kim, Minah Suh, Myunghwan Choi, and Chun Gwon Park for their help.

## AUTHOR CONTRIBUTIONS

J.-B.C., Y.-G.Y., J.S., Y.S., and J.K. conceived the main idea. J.S.,Y.S., and H.K. performed immunostaining and imaging. Y.-G.Y. and J.K. wrote all unmixing and imaging processing programs. I.C. performed mRNA-related experiments. All authors wrote the paper and contributed to the editing of the paper. J.-B.C. and Y.-G.Y. supervised this work.

## COMPETING FINANCIAL INTERESTS

J.-B.C., Y.-G.Y., J.S., Y.S. H.K., J.K., have applied for patents on PICASSO (KR patent application 10-2020-0088091, 10-2020-0106838, 10-2020-0109519, and US patent application 17/132,628).

