## Supplementary Information for "PICASSO: Ultra-multiplexed fluorescence imaging of biomolecules through single-round imaging and blind source unmixing"

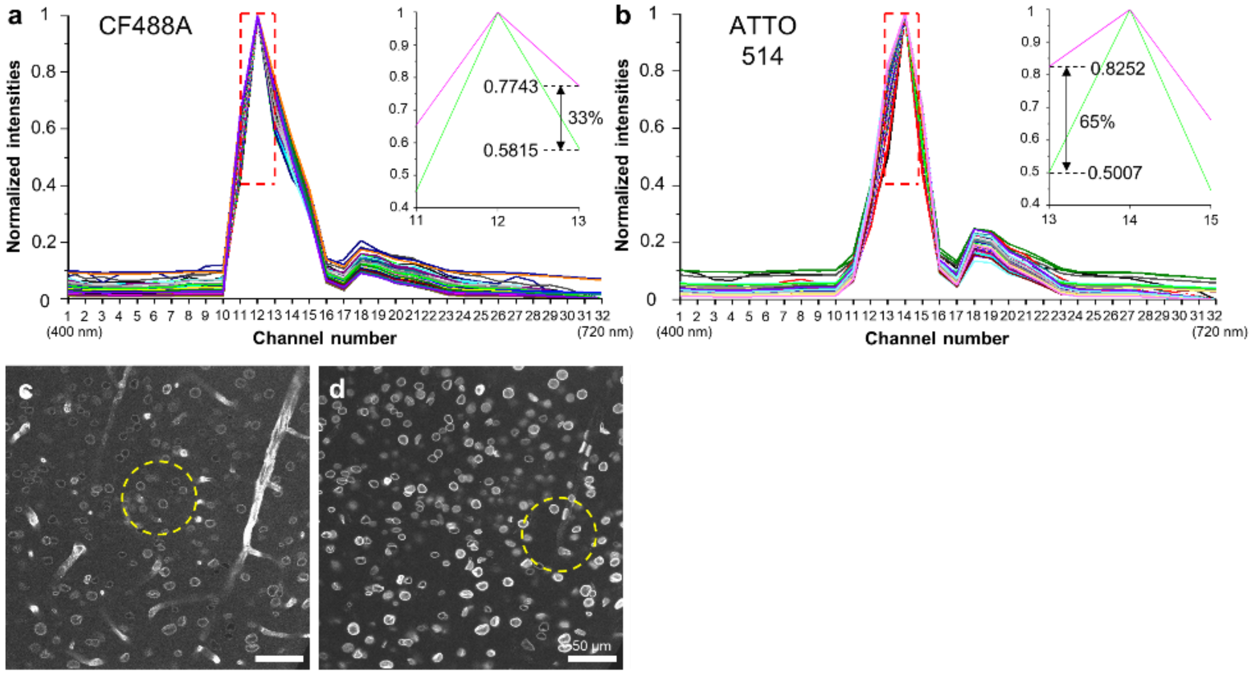


**Supplementary Figure 1**. **The variation in the emission spectra of fluorophores and their effects on the unmixing performance of linear unmixing**. (**a–b**) Normalized emission spectra of CF488A and ATTO514 measured from mouse brain slices. Brain slices were stained with an antibody conjugated with either CF488A or ATTO514 and then imaged at 32 detection channels with a bandwidth of 10 nm. Multiple brain slices labelled with different antibodies were used for the spectrum measurement. Images were acquired from various brain regions under various imaging conditions (i.e. laser intensity and camera gain). To precisely measure the emission spectra, excluding the effect of autofluorescence or background signals from non-specifically bound antibodies, only pixels with the top 1% brightness were selected and used for the emission spectrum measurements. (**a**) Normalized emission spectra of CF488A. (**b**) Normalized emission spectra of ATTO514. In **a** and **b**, each line is an emission spectrum acquired from one image. The antibodies used were antibodies against NeuN, GFAP, PV ZNF3, GluT1, or MAP2. Insets are the magnified views of the red boxed regions, showing the uppermost spectrum (magenta) and lowermost spectrum (green) among the acquired spectra. Note that the CF488A and ATTO514 spectra show a significant amount of variations. (**c–f**) Effects of the variations in the emission spectra on the performance of reference-based unmixing. A mouse brain slice was stained with a CF488A-conjugated antibody against GluT1 and an ATTO514-conjugated antibody against lamin B1 and then imaged using the same linear detector. The acquired 32-channel images were unmixed *via* either (**c**, **d**) linear unmixing. Reference spectra shown in **a** and **b** were used. Due to the variation in the spectra, two fluorophore signals were not completely unmixed.

In **c**, only GluT1 should be visible, but lamin B1 is also visible (yellow dotted circle). In **d**, only lamin B1 should be visible, but GluT1 is also visible (yellow dotted circle).

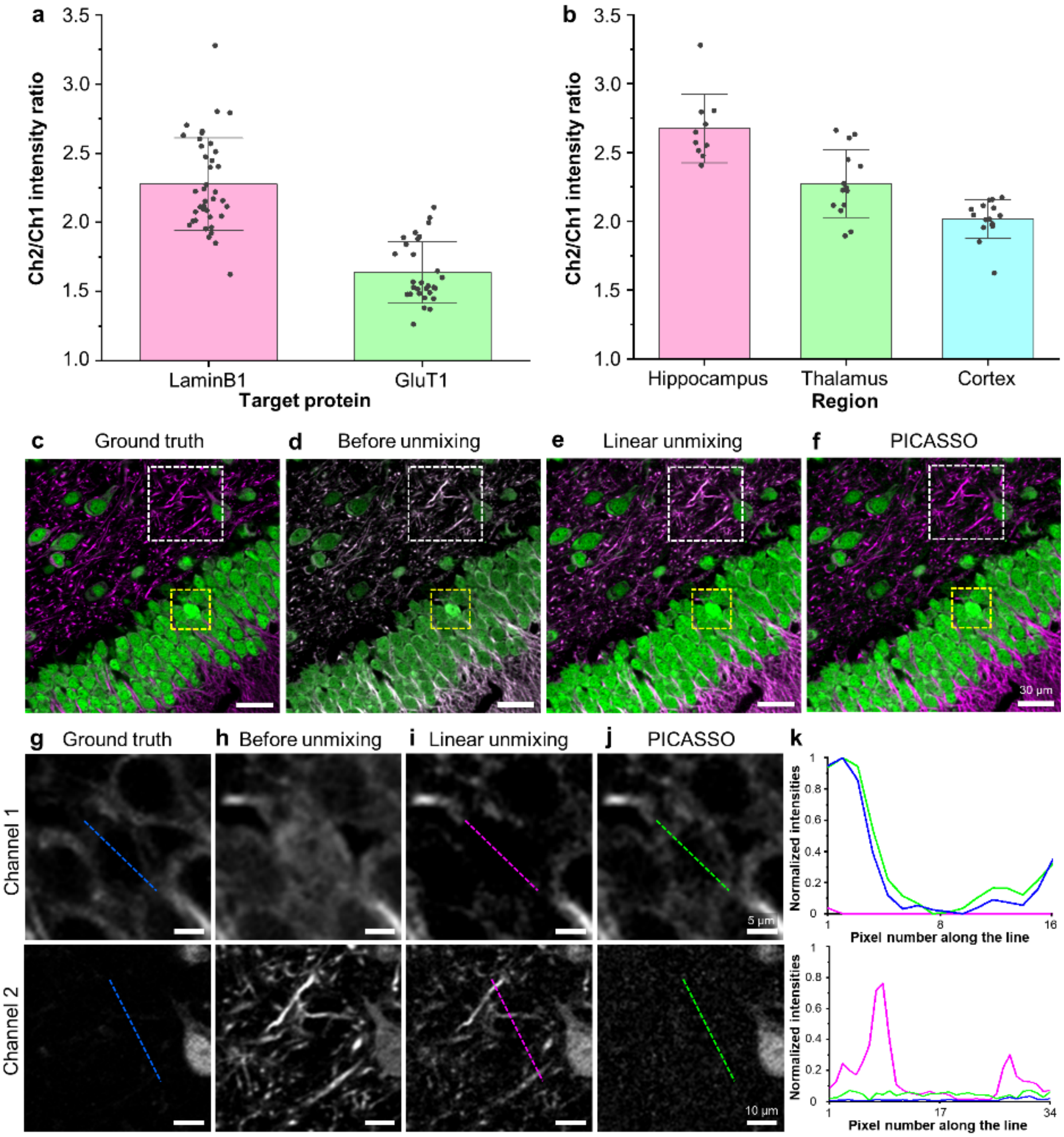


**Supplementary Figure 2**. **The variation in the emission spectra of fluorophores depending on the target proteins and imaging regions and their effects on the unmixing performance of linear unmixing.** Two mouse brain slices were labelled with an ATTO514-conjugated preformed antibody complex against lamin B1 or GluT1. Two-channel images were acquired in various brain regions (hippocampus, thalamus, and cortex) at two detection ranges (channel 1: 490–525 nm and channel 2: 525–560 nm). Due to the variation in the protein expression levels in different brain regions, laser intensities were manually adjusted when acquiring images. However, the ratio between the laser intensities used to acquire the first and second images was kept constant. The fluorescence intensity ratio in the two channels (channel 2/channel 1) was measured only from pixels with the top 1% brightness, in the same way as in **Supplementary Fig**. **1**. Unlike Supplementary Fig. 1, where the emission spectra of fluorophores were acquired over 32 narrow detection channels, we measured the fluorescence intensity ratio only in two 35 nm-wide detection channels to maximize the signal intensity and to minimize the effect of autofluorescence and noise. (**a**) The fluorescence intensity ratios measured using lamin B1 or GluT1. Lamin B1: 2.28 ± 0.33 (mean ± standard deviation). GluT1: 1.64 ± 0.22. (**b**) The fluorescence intensity ratios measured by using lamin B1 from different brain regions. Hippocampus: 2.67 ± 0.25. Thalamus: 2.27 ± 0.25. Cortex: 2.02 ± 0.14. Note that the fluorescence intensity ratio varied depending on the target protein and the subregion of the brain. It also showed a significant level of variation within the same subregion. (**c–k**) Linear unmixing *vs*. PICASSO. The unmixing performances of linear unmixing and PICASSO were compared by using two mixed images acquired at two detection channels. The fluorescence intensity ratios (Ch2/Ch1) of the two fluorophores used were measured from separate brain slices and then used as reference spectra. To minimize the specimen-to-specimen variation, two subsequent brain slices of the same mouse brain were used in this experiment. The first slice was stained with two antibodies, one against MAP2 and the other one against NeuN. The antibody against MAP2 bore CF488A, and the antibody against NeuN bore ATTO514. Two-channel images were acquired at two detection ranges (channel 1: 500–525 nm and channel 2: 525–550 nm) in the dentate gyrus of the hippocampal region. This slice was also stained with other antibodies, one against MAP2 bearing CF568 and the other one against NeuN bearing CF633. The CF568 and CF633 signals were used as ground truth. The second brain slice was divided into two pieces, and each piece was stained either with an antibody against MAP2 (CF488A) or NeuN (ATTO514). The Ch2/Ch1 ratios of CF488A and ATTO514 were measured from the hippocampal regions of these pieces. (**c–f**) Merged images of channel 1 and channel 2. MAP2; magenta, NeuN; green. (**c**) Ground-truth images of MAP2 and NeuN generated by merging the CF568 and CF633 signals. (**d**) Mixed image acquired at the two detection channels. (**e**) Unmixing of the image shown in **d** *via* linear unmixing. (**f**) Unmixing of the image shown in **d** *via* PICASSO. (**g–j**) Magnified view of boxed region in **c**–**f**. The first row shows the magnified view of channel 1 of the yellow boxed region in **c**–**f**. The second row shows the magnified view of channel 2 of the white boxed region in **c**–**f**. (**g**) Ground-truth image shown in **c**. (**h**) Mixed image shown in **d**. (**i**) Unmixing *via* linear unmixing shown in **e**. (**j**) Unmixing *via* PICASSO shown in **f**. (**k**) Intensity profiles along the dotted lines in **g–j**. Blue line: ground truth, magenta line: linear unmixing, green line: PICASSO. Note that linear unmixing over-subtracted the NeuN signal from channel 1 of the mixed image, indicated by the discrepancy between the magenta line (linear unmixing) and blue line (ground truth) in the upper graph in **k**. The unmixing result of PICASSO matched well with the ground truth, indicated by the match between the green line (PICASSO) and blue line (ground truth) in the same graph. Linear unmixing also under-subtracted the MAP2 signal from channel 2, as indicated by the discrepancy between the magenta line (linear unmixing) and blue line (ground truth) in the lower graph in **k**. PICASSO successfully removed such filamentous structures from channel 2, as indicated by the match between the green line (PICASSO) and blue line (ground truth) in the same graph. Even the reference spectra were measured from the same brain region by using the same antibody, linear unmixing sometimes resulted in such incomplete unmixing, possibly due to the variation in the emission spectra shown in **b**.

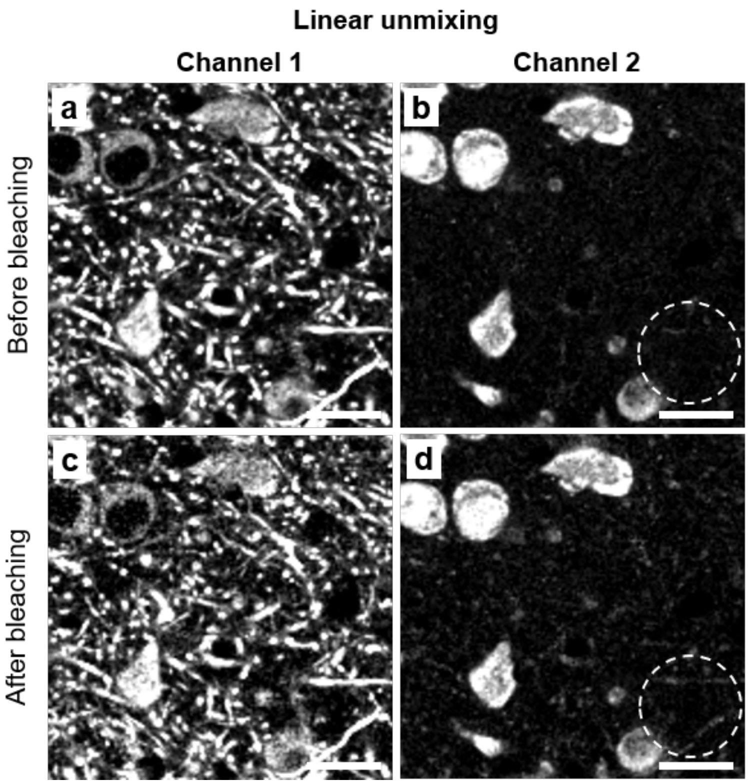


**Supplementary Figure 3**. **The effect of photobleaching on the performance of reference-based unmixing**. A mouse brain slice was labelled with two preformed antibody complexes against MAP2 (CF488A) and NeuN (ATTO514) and then imaged at 32 detection channels with a bandwidth of 10 nm multiple times. Two mixed images, which were acquired at the beginning and end of the repeated imaging process, were unmixed *via* linear unmixing to study the effect of photobleaching on the unmixing performance. The fluorescence intensity of the mixed imaged acquired at the end was around half of that of the mixed images acquired at the beginning of the repeated imaging process. (**a**,**b**) Unmixing of the mixed images acquired at the beginning. The fluorophores were not photobleached when the images were acquired. (**a**,**b**) Ch1 and Ch2 after unmixing *via* linear unmixing. In **a**, only MAP2 should be visible. In **b**, only NeuN should be visible. In **b**, MAP2 structures were visible (dotted circle), but at a low signal intensity. (**c**,**d**) Unmixing of the mixed images acquired at the end. The fluorophores were half-bleached when the images were acquired due to the repeated imaging. In **d**, the fluorescence intensity of the MAP2 structures (dotted circle) became stronger due to the incomplete unmixing. Such incomplete unmixing may be attributed to the change in the emission spectra of the fluorophores after photobleaching. Scale bars: 20 μm.

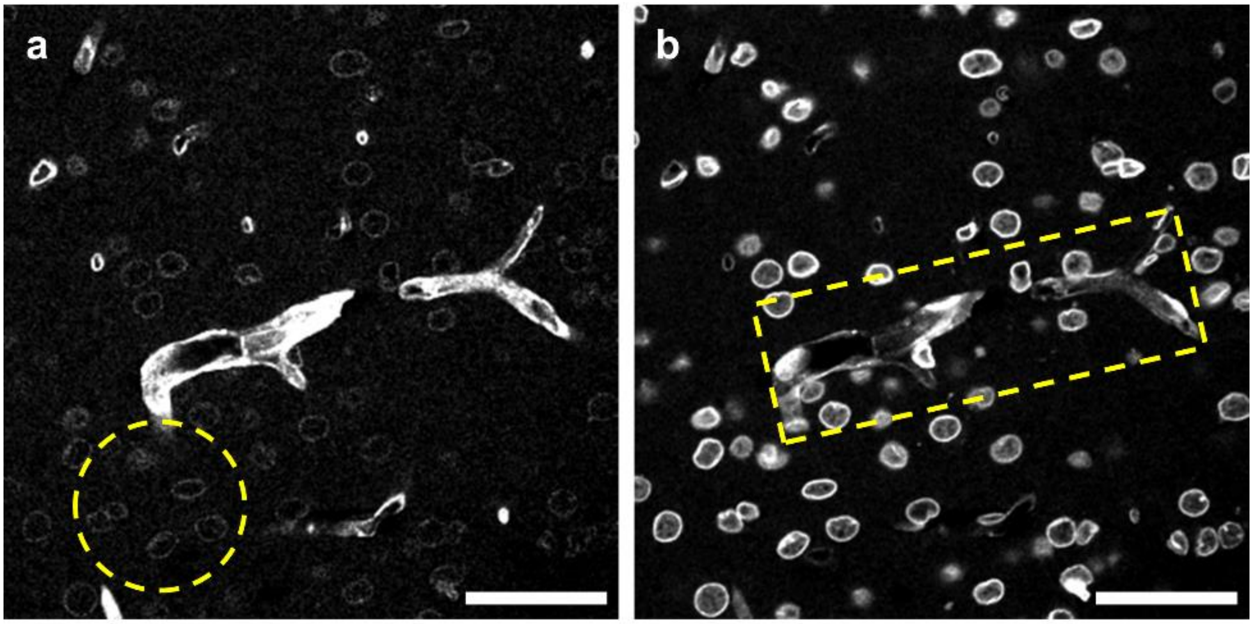


**Supplementary Figure 4**. **Unmixing *via* NMF**. The mouse brain slice was stained with two preformed antibody complexes against GluT1 (CF488A) and lamin B1 (ATTO514), then imaged at 32 detection channels with a 10-nm bandwidth from 400 nm to 720 nm. The 32-channel images were unmixed *via* NMF. (**a**) Ch1 of the unmixed images. Only GluT1 signals should be visible, but the lamin B1 signal (yellow circled region) is visible. (**b**) Ch2 of the unmixed images. Only lamin B1 signal should be visible, but the GluT1 signal (yellow boxed region) is visible.

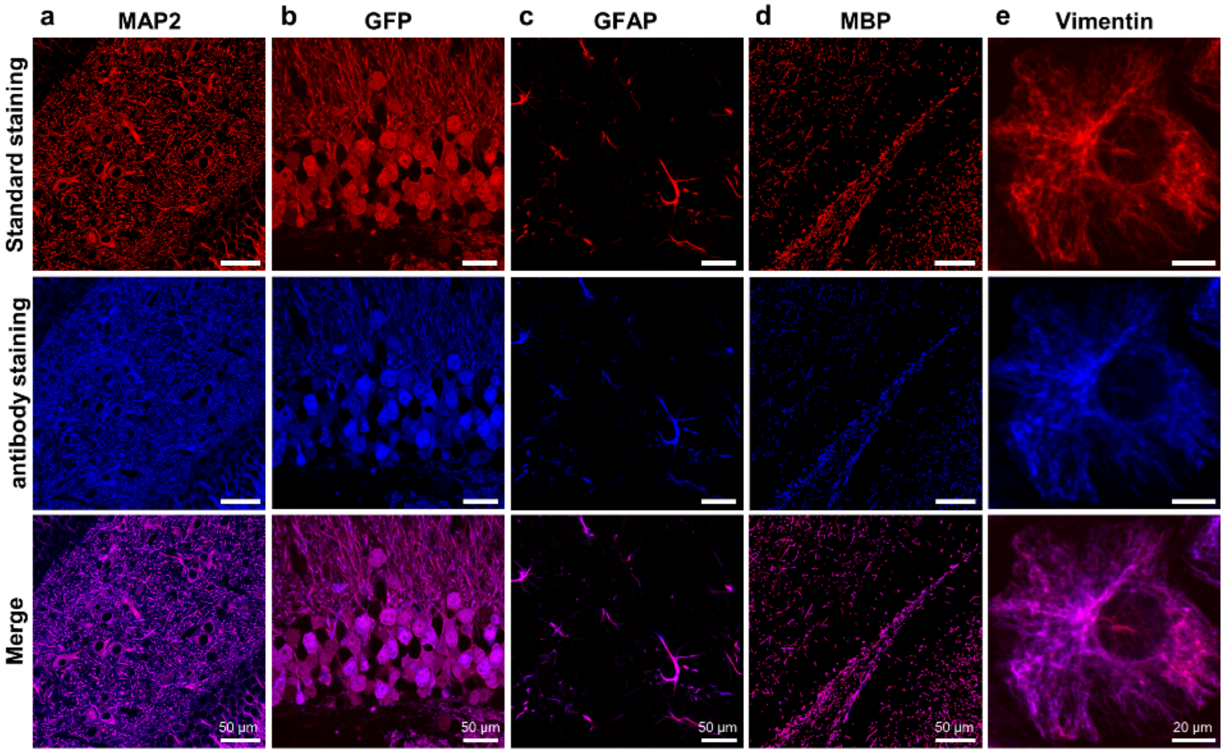


**Supplementary Figure 5**. **Validation of the staining with preformed antibody complexes**. Confocal microscopy images of mouse brain slices or cultured cells, each of which was stained with a conventional antibody and a preformed antibody complex against the same target proteins. The target proteins were (**a**) MAP2, (**b**) GFP, (**c**) GFAP, (**d**) MBP, and (**e**) vimentin. For **a**,**c**,**d**, wildtype mouse brain slices were used. For **b**, a Thy1-YFP mouse brain slice was used. For **e**, cultured BS-C-1 cells were used. For all tested targets, the conventional antibody staining and preformed antibody staining showed identical staining patterns, as shown in the 3rd row.

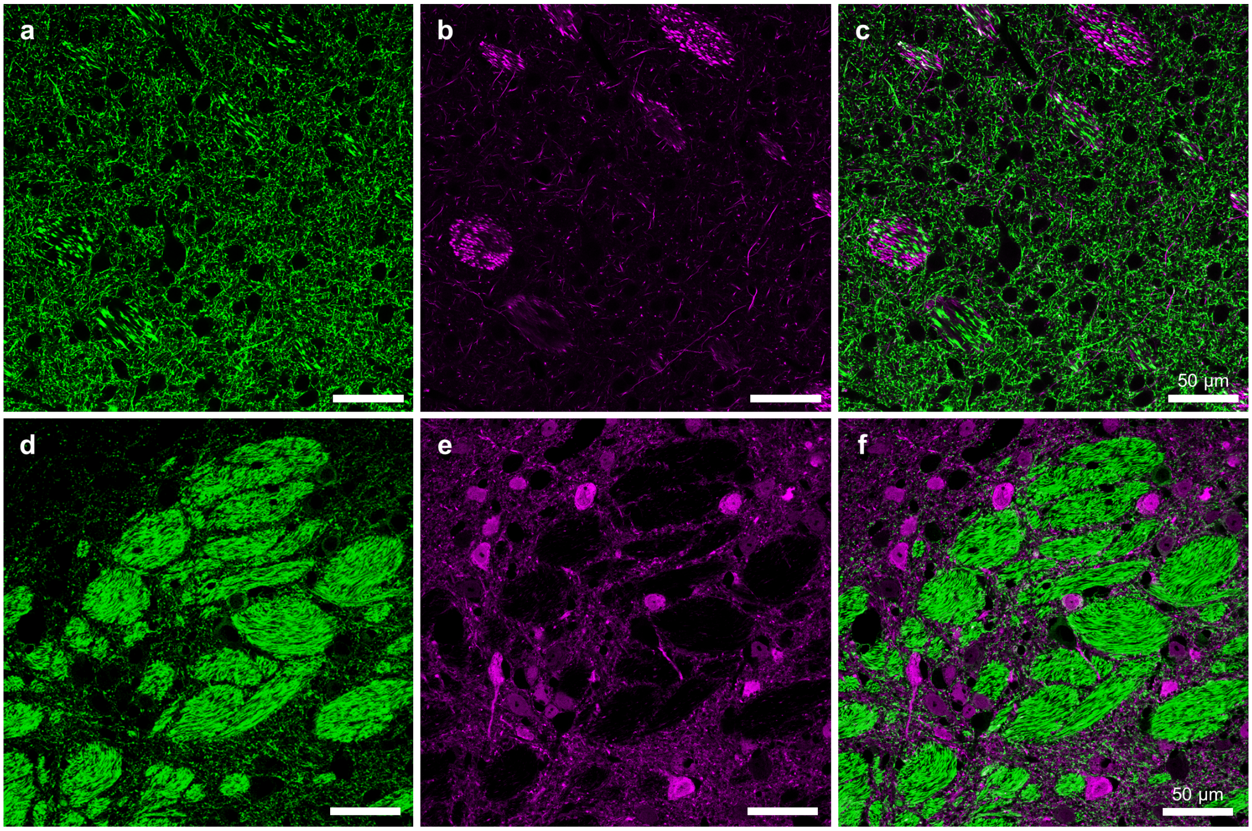


**Supplementary Figure 6**. **Validation of the absence of crosstalk between two preformed antibody complexes**. A Thy1-YFP mouse brain slice was stained with two preformed rabbit antibody complexes, one against YFP (unlabelled) and the other against neurofilament-H (NF-H, labelled with CF568). (**a**) YFP fluorescence. (**b**) CF568 fluorescence, showing NF-H. (**c**) Merged image of **a** and **b.** No crosstalk between the signals of the endogenous YFP and the signals of the antibody-labelled NF-H is visible. (**d**) YFP fluorescence. (**e**) CF633 fluorescence, showing calbindin. (**f**) Merged image of **d** and **e.** No crosstalk between the signals of the endogenous YFP and the signals of the antibody-labelled calbindin is visible.

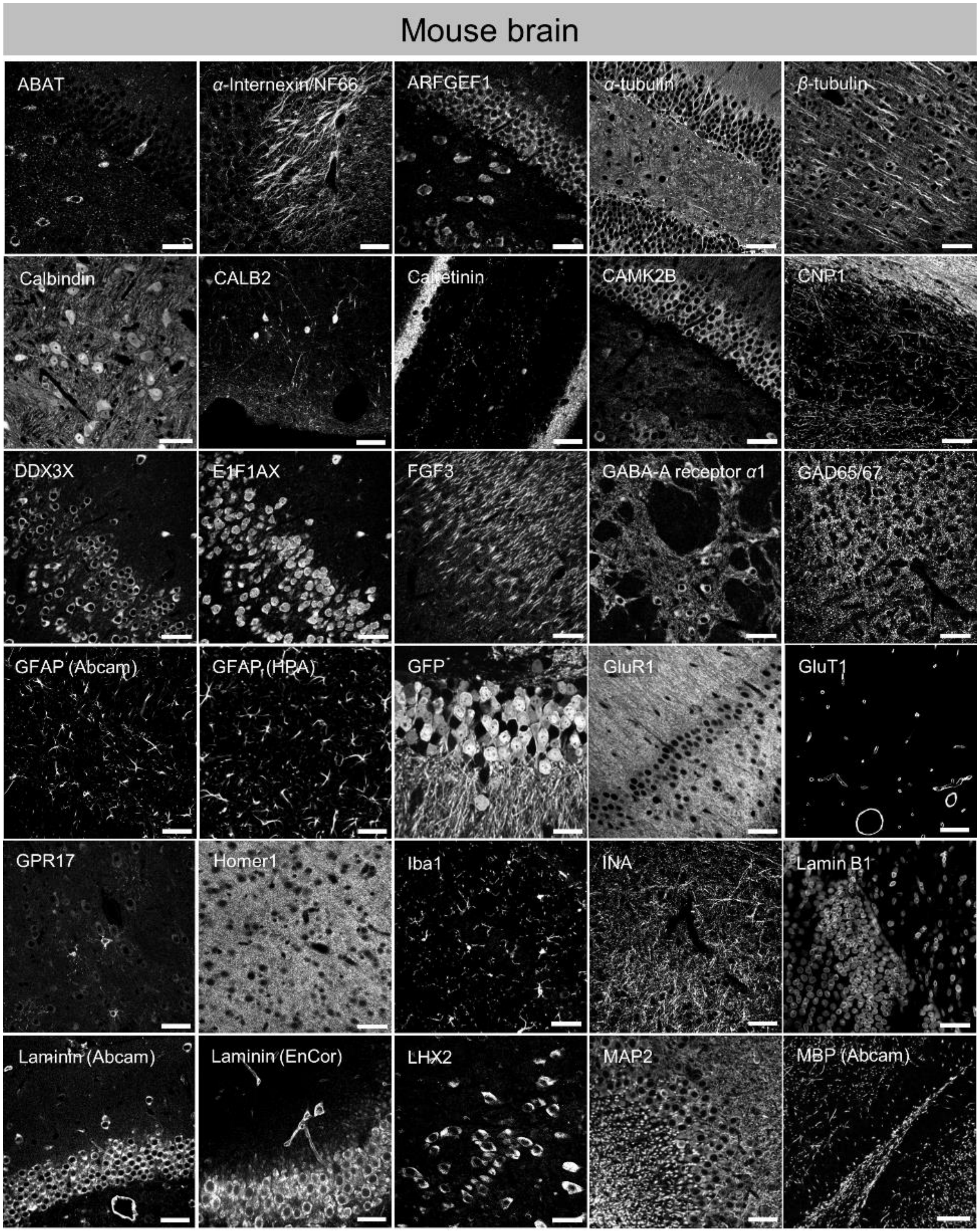
Mouse brain
ABAT
α-Internexin/NF66
ARFGEF1
α-tubulin
β-tubulin
Calbindin
CALB2
Calretinin
CAMK2B
CNP1
DDX3X
E1F1AX
FGF3
GABA-A receptor α1
GAD65/67
GFAP (Abcam)
GFAP (HPA)
GFP
GluR1
GluT1
GPR17
Homer1
Iba1
INA
Lamin B1
Laminin (Abcam)
Laminin (EnCor)
LHX2
MAP2
MBP (Abcam)

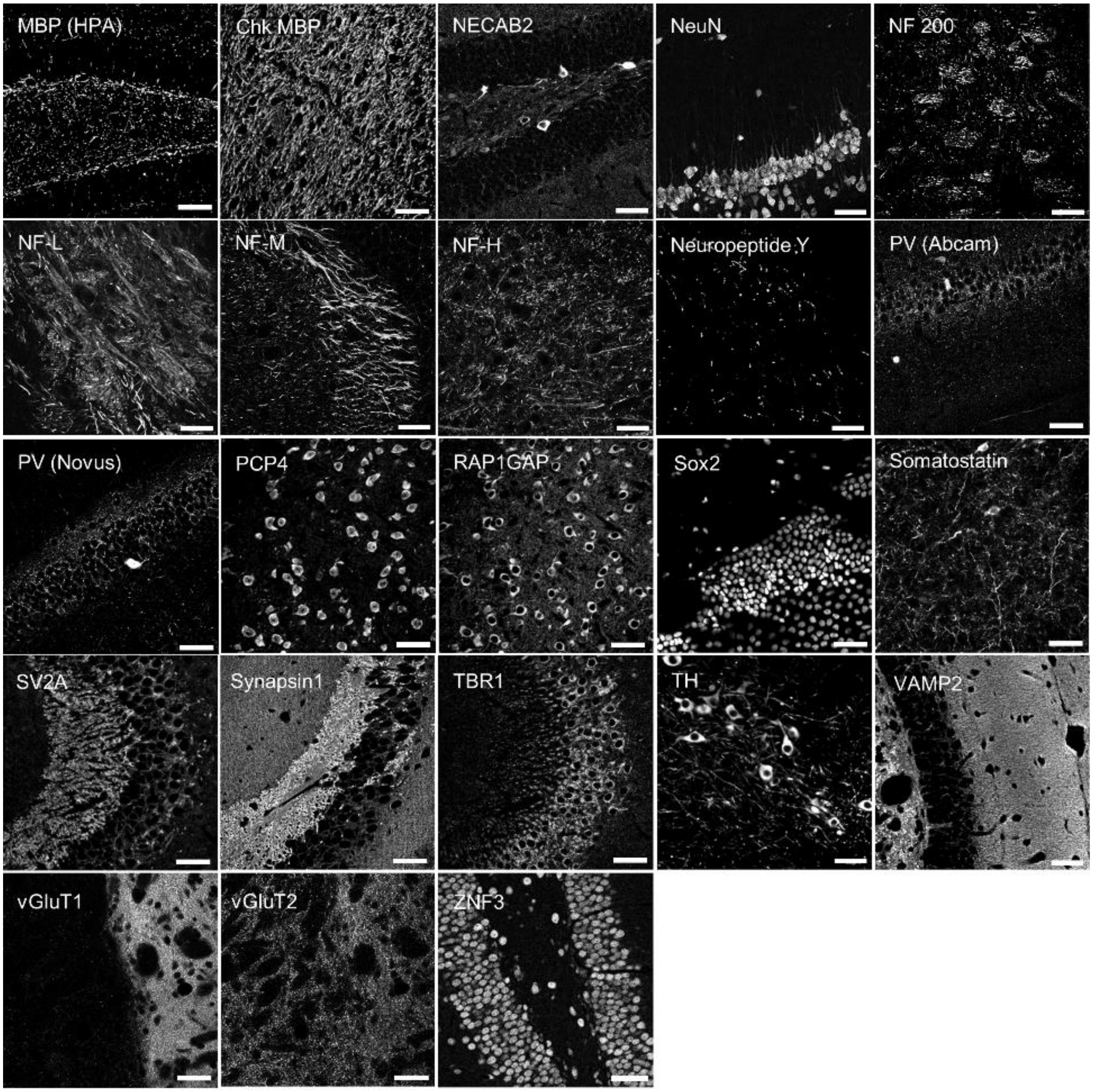
MBP (HPA)
Chk MBP
NECAB2
NeuN
NF 200
NF-L
NF-M
NF-H
Neuropeptide Y
PV (Abcam)
PV (Novus)
PCP4
RAP1GAP
Sox2
Somatostatin
SV2A
Synapsin1
TBR1
TH
VAMP2
vGluT1
vGluT2
ZNF3

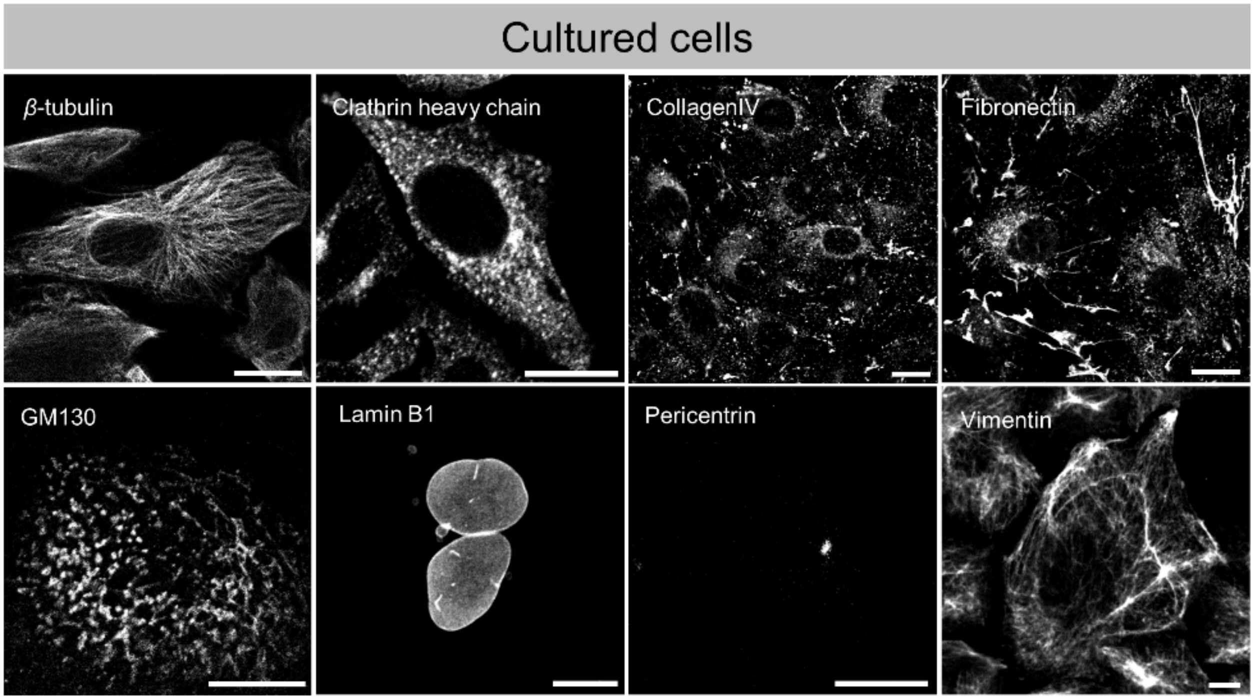


**Supplementary Figure 7**. **Confocal microscopy images of various proteins labelled with preformed antibody complexes**. Fifty-nine different antibodies were used, targeting diverse types of proteins, such as cell type-specific proteins, receptor proteins, transcription factor proteins, neurofilament-related proteins, and synaptic vesicle proteins. Detailed information about the antibodies is given in **Supplementary Table 1**. Scale bars: brain, 50 μm; cell, 20 μm.

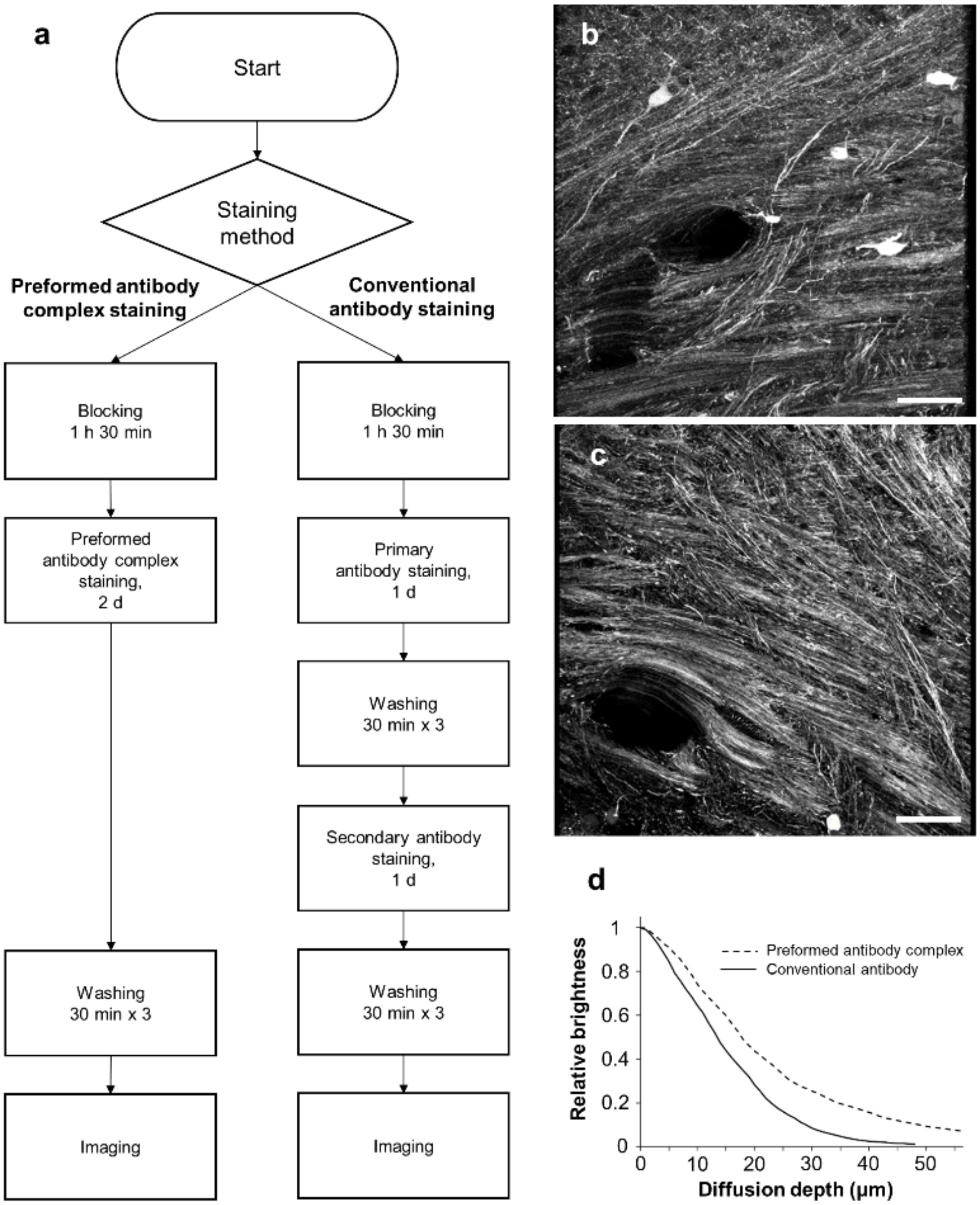


**Supplementary Figure 8**. **Comparison of the diffusion depths between staining with a preformed antibody complex and a conventional antibody**. (**a**) Experiment workflow. (**b–c**) Maximum intensity projections of z-stack images of the thalamus from mouse brain slices stained with either a preformed antibody complex or a regular antibody against calbindin. (**b**) Result of a preformed antibody complex. (**c**) Result of a conventional antibody. (**d**) Diffusion depth profiles of the specimens shown in **b** and **c**. Scale bars: 50 μm.

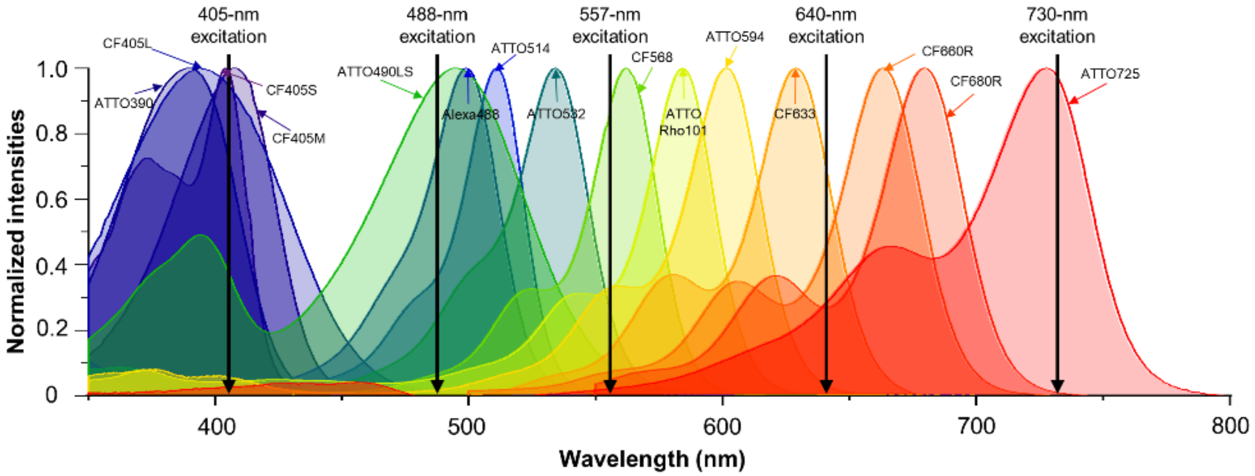


**Supplementary Figure 9**. **Excitation spectra of the 15 fluorophores used in Fig. 1d**. Black arrows indicate the wavelengths of the five standard excitation lasers.

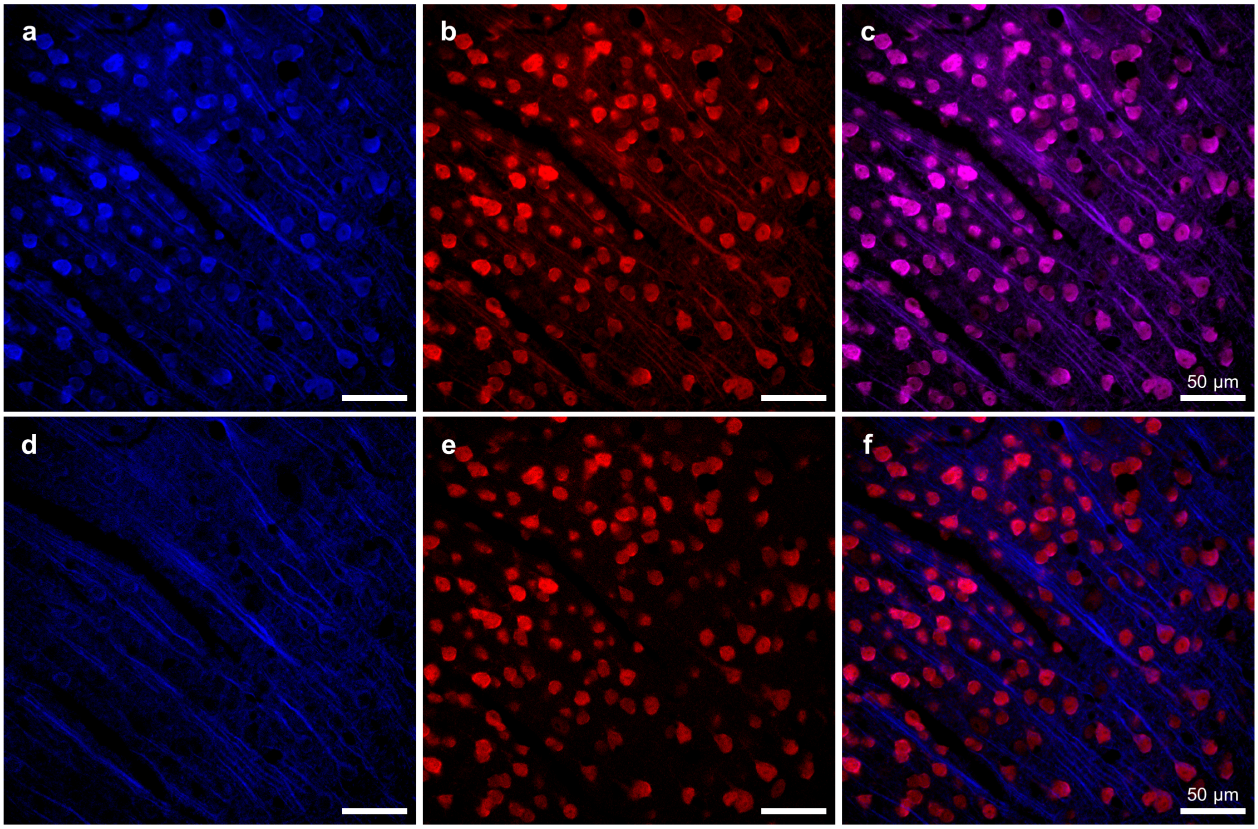


**Supplementary Figure 10**. **Unmixing of two spectrally overlapping fluorophores with an 8-nm separation between their emission peaks *via* PICASSO**. (**a**–**c**) Before unmixing. (**d**–**f**) After unmixing. (**a**,**d**) Ch1. (**b**,**e**) Ch2. (**c**,**f**) Merged image of Ch1 and Ch2. A mouse brain slice was stained with antibodies against MAP2 (ATTO488) and NeuN (ATTO514) and then imaged at two detection channels. The first detection channel was 500–530 nm, and the second detection channel was 530–560 nm. The two images were then unmixed *via* PICASSO.

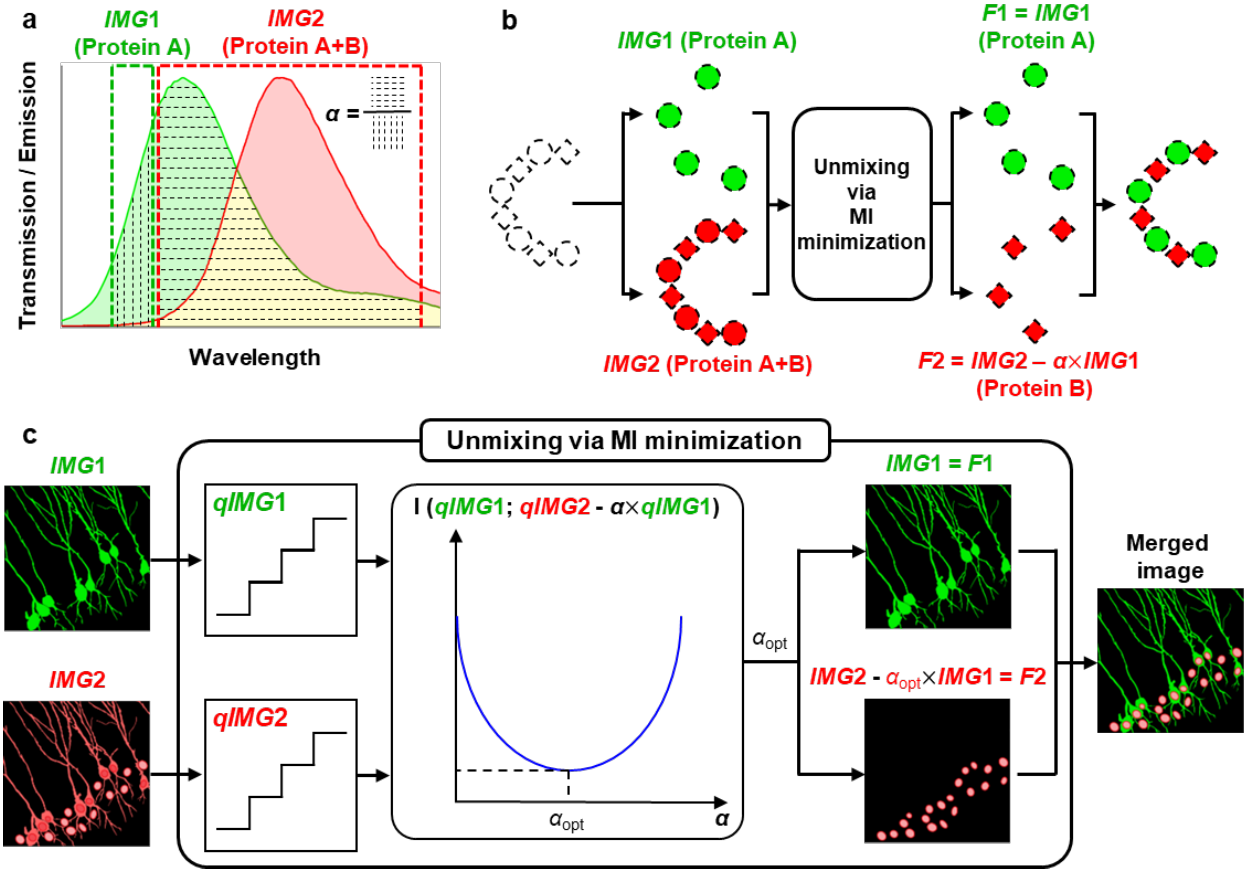


**Supplementary Figure 11**. **Experimental design to validate the use of MI minimization as a means of unmixing**. (**a**) Emission spectra of two spectrally overlapping fluorophores (green and red solid lines and coloured regions) and detection channels (green and red dotted rectangles) used in this validation study. Here, $\alpha$ is the ratio of the area with horizontal dotted lines to that with vertical dotted lines. $IMG1,2$: images acquired at the first (green dotted rectangle) and second (red dotted rectangle) detection channels. (**b**) Example of PICASSO images before and after unmixing. Dotted circles and squares: two target proteins to be imaged in a specimen. Coloured circles and squares: structures shown in images. $F1, F2$: images after unmixing. (**c**) Unmixing *via* MI minimization. $qIMG1,2$: quantized $IMG1,2$. $I(qIMG1; qIMG2 - \alpha \times qIMG1)$: mutual information between $qIMG1$ and $qIMG2 - \alpha \times qIMG$. $\alpha_{\mathrm{opt}}$: $\alpha$ minimizing $I(qIMG1; qIMG2 - \alpha \times qIMG1)$.

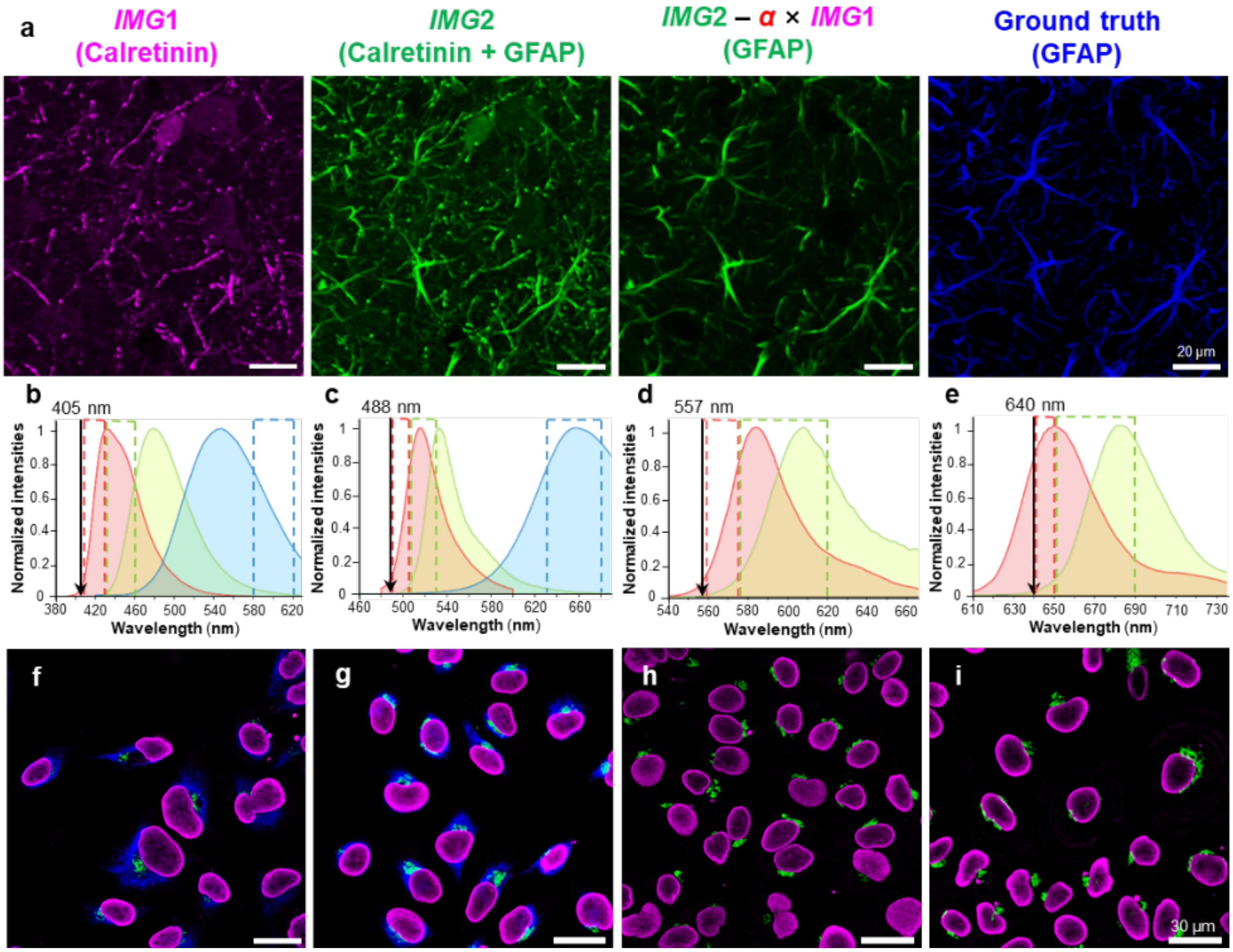


**Supplementary Figure 12**. **Experimental validation and demonstrations of unmixing *via* MI minimization**. (**a**) Experimental validation. A mouse brain slice was stained with two antibody complexes against calretinin (CF568) and GFAP (ATTORho101) along with one regular antibody against GFAP (CF633). The regular antibody staining was used as the ground-truth image of GFAP. From the left to the right, *IMG*1, *IMG*2, unmixed image (*IMG*2 – $\alpha \times$ *IMG*1), and the ground-truth image are shown respectively. *IMG*1 acquired at the first detection channel where only the calretinin (CF568) signal is visible. *IMG2* acquired at the second detection channel where both calretinin (CF568) and GFAP (ATTORho101) signals are visible. After unmixing, the calretinin signal was successfully subtracted from *IMG*2. The resulting unmixed GFAP image was identical to the ground-truth image. (**b–i**) Demonstrations of two- or three-colour multiplexed imaging with a single excitation laser. Two spectrally overlapping fluorophores were used for each excitation laser. An additional large Stokes shift fluorophore was used for a 405-nm and 488-nm excitation laser. (**b–e**) Normalized emission spectra of fluorophores (solid lines and coloured regions) and detection channels (coloured dotted rectangles) for each excitation laser (black arrow). (**b**) CF488A (red), ATTO514 (green), and ATTO490LS (blue) excited by a 488-nm laser. (**c**) CF405S (red), ATTO390 (green), and CF405L (blue) excited by a 405-nm laser. (**d**) CF568 (red) and ATTORho101 (green) excited by a 557-nm laser. (**e**) CF633 (red) and CF660R (green) excited by a 640-nm laser. (**f–i**) Two- or three-colour multiplexed immunofluorescence images of HeLa cells after unmixing *via* MI minimization. Magenta, lamin A/C; green, GM130; blue, vimentin. The fluorophores and detection channels shown in **b–e** were used. See the first paragraph of **Supplementary Note 2** for details.

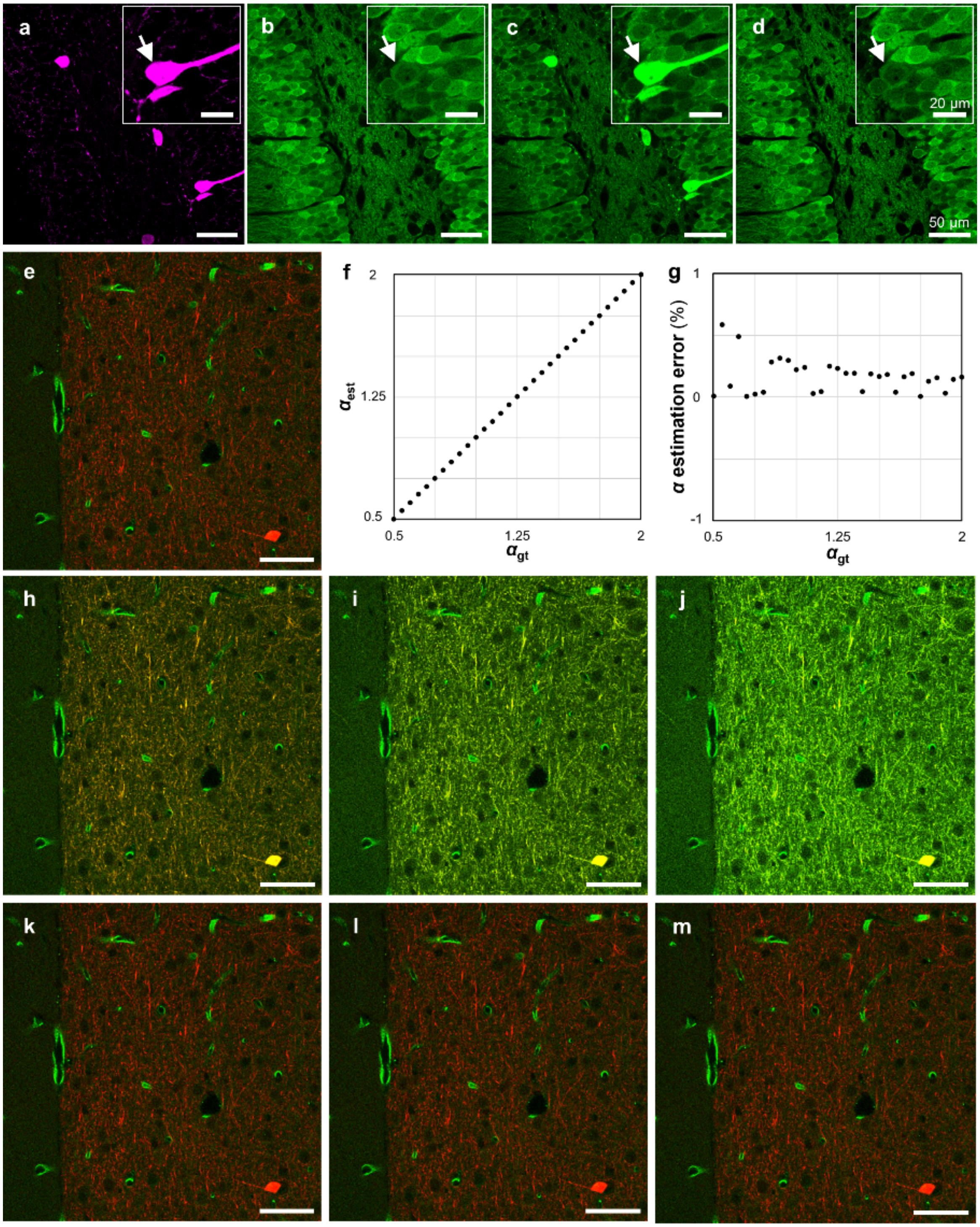


**Supplementary Figure 13**. **Validation of the accuracy of the unmixing *via* MI minimization**. (**a–d**) Validation of unmixing *via* MI minimization. (**a**,**b**) Immunofluorescence image of parvalbumin (**a**) and calbindin (**b**). (**c**) Synthetic mixed image (*SynIMG*) of **a** ($F1$) and **b** ($F2$) with an $\alpha$ value of 2 ($SynIMG = 2 \times F1 + F2$). Two mixed images, which are $F1$ and *SynIMG*, were unmixed *via* MI minimization. (**d**) The resulting unmixed image, showing only calbindin. Note that the image in **d** (after unmixing) is identical to the image in **b** (ground-truth image). (**e–m**) Quantitative analysis of the accuracy of unmixing *via* MI minimization. (**e**) Two-colour image of a mouse brain slice stained with antibodies against calbindin and GluT1. The proteins were imaged using different excitation lasers, resulting in non-mixed images. Red ($F1$), calbindin; green ($F2$), GluT1. (**f**) Ground-truth $\alpha_{\text{gt}}$ *vs*. estimated $\alpha$ ($= \alpha_{\text{est}}$). The two channels shown in **e** were used to generate a synthetic mixed image with a known $\alpha$ ($= \alpha_{\text{gt}}$). The synthetic mixed images were then unmixed *via* MI minimization, and the resulting estimated $\alpha_{\text{est}}$ was compared with the ground-truth $\alpha_{\text{gt}}$. (**g**) $\alpha$ estimation error. (**h–j**) Synthetic mixed images used for the simulation. Red, same as the calbindin channel (red) of the image shown in **e**; green, mixed image obtained by the linear addition of the calbindin channel (red) of the image shown in **e** multiplied by $\alpha_{\text{gt}}$ and the GluT1 channel (green) of the image shown in **e**. $\alpha_{\text{gt}}$ was (**h**) 0.5, (**i**) 1.25, (**j**) 2. (**k–m**) Unmixed two-channel images obtained by unmixing *via* MI minimization. The image shown in **k**, **l**, and **m** was obtained by unmixing the image shown in **h**, **i**, and **j**, respectively. Scale bars: 50 µm. See the first paragraph of **Supplementary Note 2** for details. Note that unmixed images shown in **k**,**l**, and **m** are identical to the ground-truth image shown in **e**.

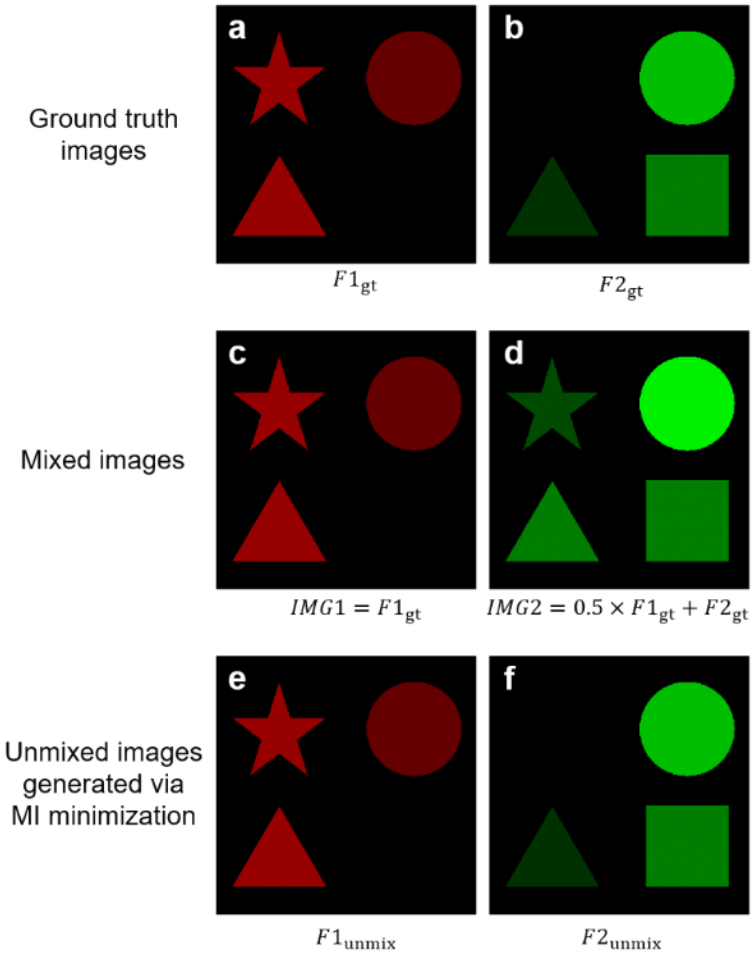


**Supplementary Figure 14**. **Unmixing of two images with mutual information**. (**a**,**b**) Ground-truth images with high mutual information. (**a**) First ground-truth image ($F1_{\mathrm{gt}}$). (**b**) Second ground-truth image ($F2_{\mathrm{gt}}$). Note that the two images have high mutual information as both contain a triangular and a circular structure. (**c**,**d**) Synthetic mixed images. (**c**) Image expected to be acquired at the first detection channel, where the signal of only the first fluorophore is collected. $IMG1 = F1_{\mathrm{gt}}$. (**d**) Image expected to be acquired at the second detection channel, where the signals of both fluorophores are collected. $IMG2 = 0.5 \times F1_{\mathrm{gt}} + F2_{\mathrm{gt}}$. (**e**,**f**) Images generated by unmixing *via* MI minimization. (**e**) The first channel of the unmixed image ($F1_{\mathrm{unmix}}$). (**f**) The second channel of the unmixed image ($F2_{\mathrm{unmix}}$). Note that the second channel of the unmixed image **f** is identical to the ground-truth image shown in **b**. See the first paragraph of **Supplementary Note 2** for details.

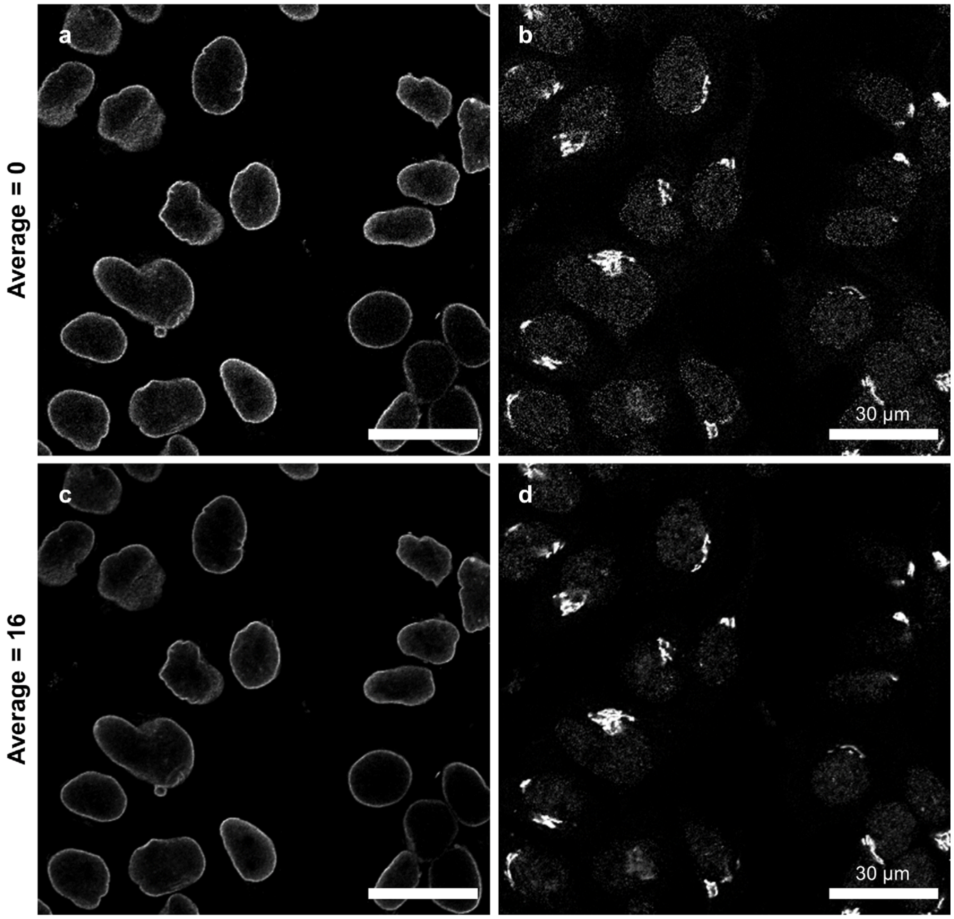


**Supplementary Figure 15**. **Reduction in the Poisson noise *via* image averaging**. Study on the Poisson noise level in the unmixed images depending on the number of averagings during the image acquisition step. Lamin B1 and GM130 of cultured cells were stained with two spectrally overlapping fluorophores and then imaged at two detection channels twice. At the first image acquisition, images were acquired without any averaging. At the second image acquisition, images were acquired 16 times and then averaged. The images were then unmixed *via* MI minimization. (**a**,**b**) Result of unmixing *via* PICASSO without image averaging. Considerably high noise values were observed in the unmixed images. (**c**,**d)** Result of unmixing *via* PICASSO with image averaging. A significant reduction in the noise was observed in the unmixed images.

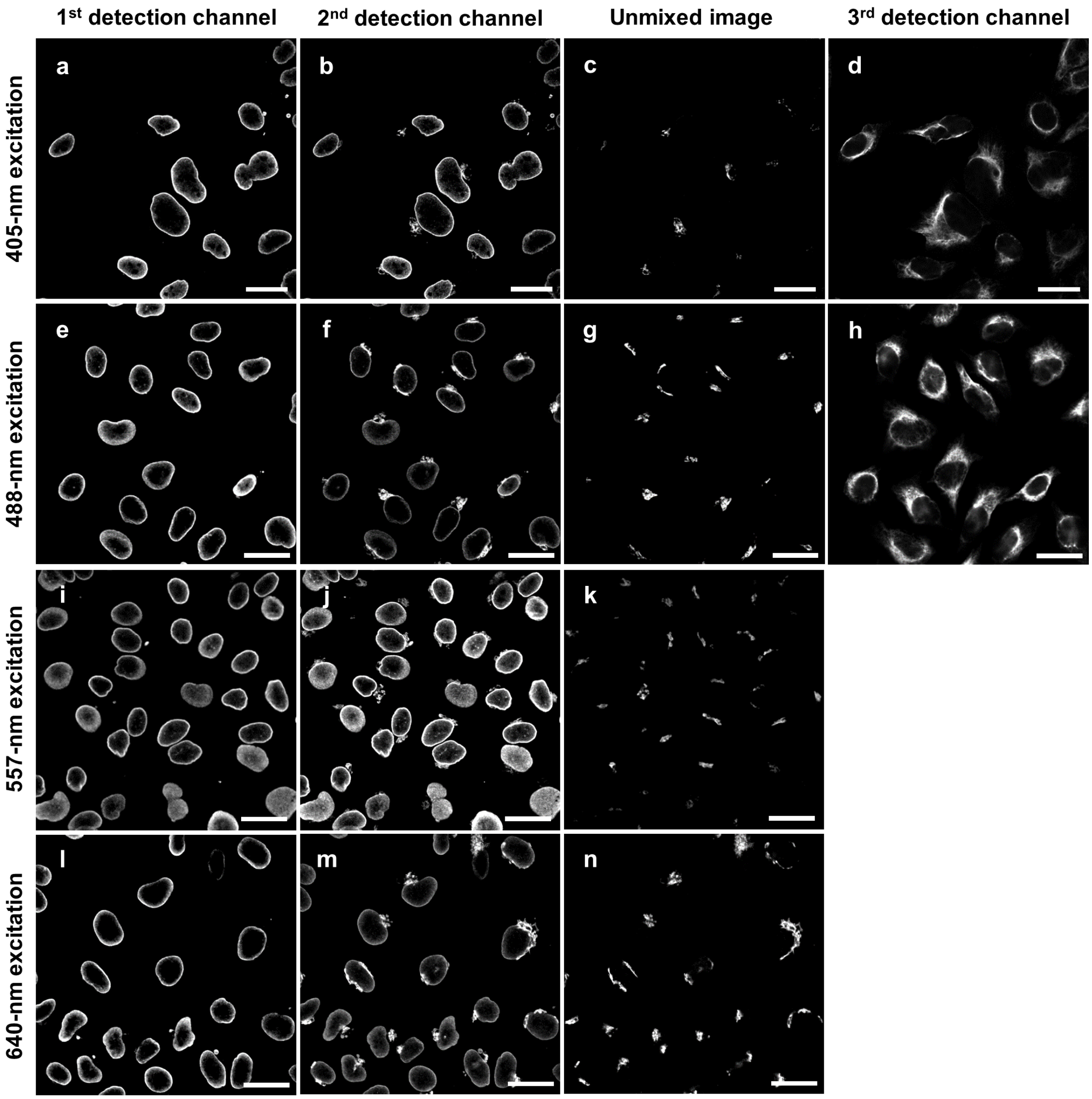


**Supplementary Figure 16**. **Unmixing *via* Gram-Schmidt orthogonalization**. Two- or three-colour multiplexed images acquired with a single excitation laser, enabled by Gram-Schmidt (GS) orthogonalization. The same mixed images used in **Supplementary Fig**. **12f–i** were unmixed *via* GS orthogonalization. Target proteins were lamin A/C (shown in the image of the 1st detection channel image), GM130 (shown in the unmixed image), and vimentin (shown in the image of the 3rd detection channel). Scale bars: 30 μm.

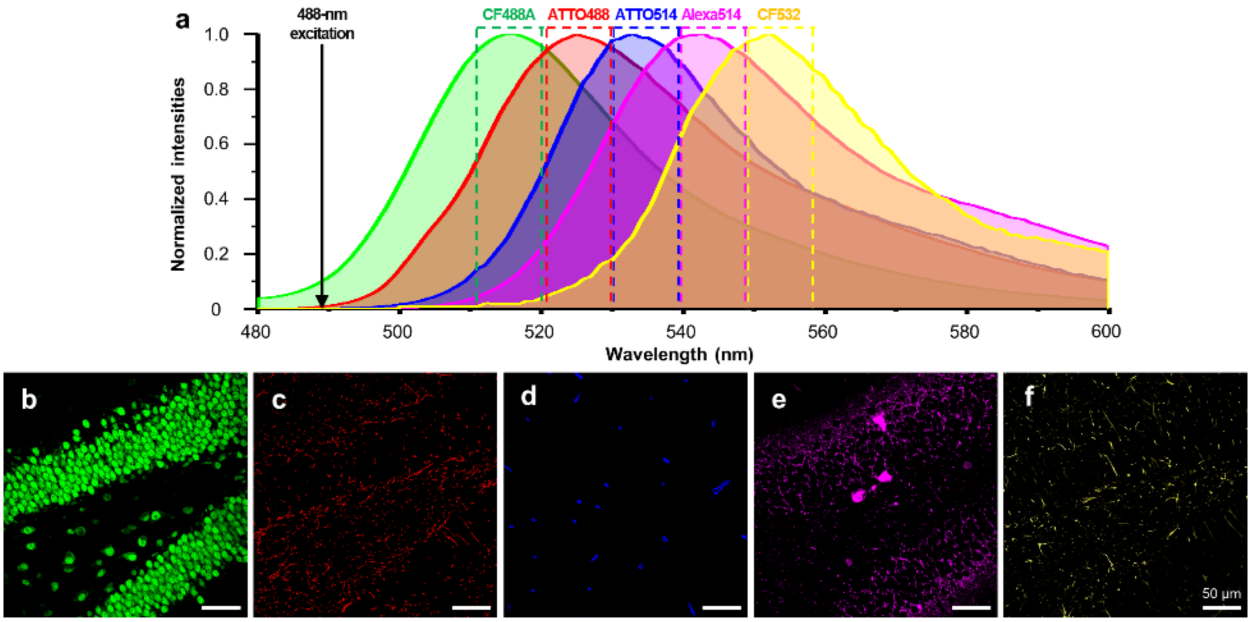


**Supplementary Figure 17**. **Unmixing of five highly overlapping fluorophores**. The ground-truth images shown in **Figure 2d** in the first row were used to generate synthetic mixed images and then unmixed *via* PICASSO. (**a**) Emission spectra of the five fluorophores used in the simulation. Emission peaks of the fluorophores were separated less than 10 nm (the emission peak of CF488A: 515.6 nm, ATTO488: 525 nm, ATTO514: 533 nm, Alexa Fluor 514: 543 nm, and CF532: 552 nm). The detection channels (dotted boxes) were defined from -5 to +5 nm of the emission peaks of the fluorophores. (**b–f**) Resulting unmixed images. (**b**) NeuN (CF488A). (**c**) CNP1 (ATTO488). (**d**) GluT1 (ATTO514). (**e**) PV (Alexa Fluor 514). (**f**) GFAP (CF532). The Pearson correlation coefficient of unmixing was 0.9938.

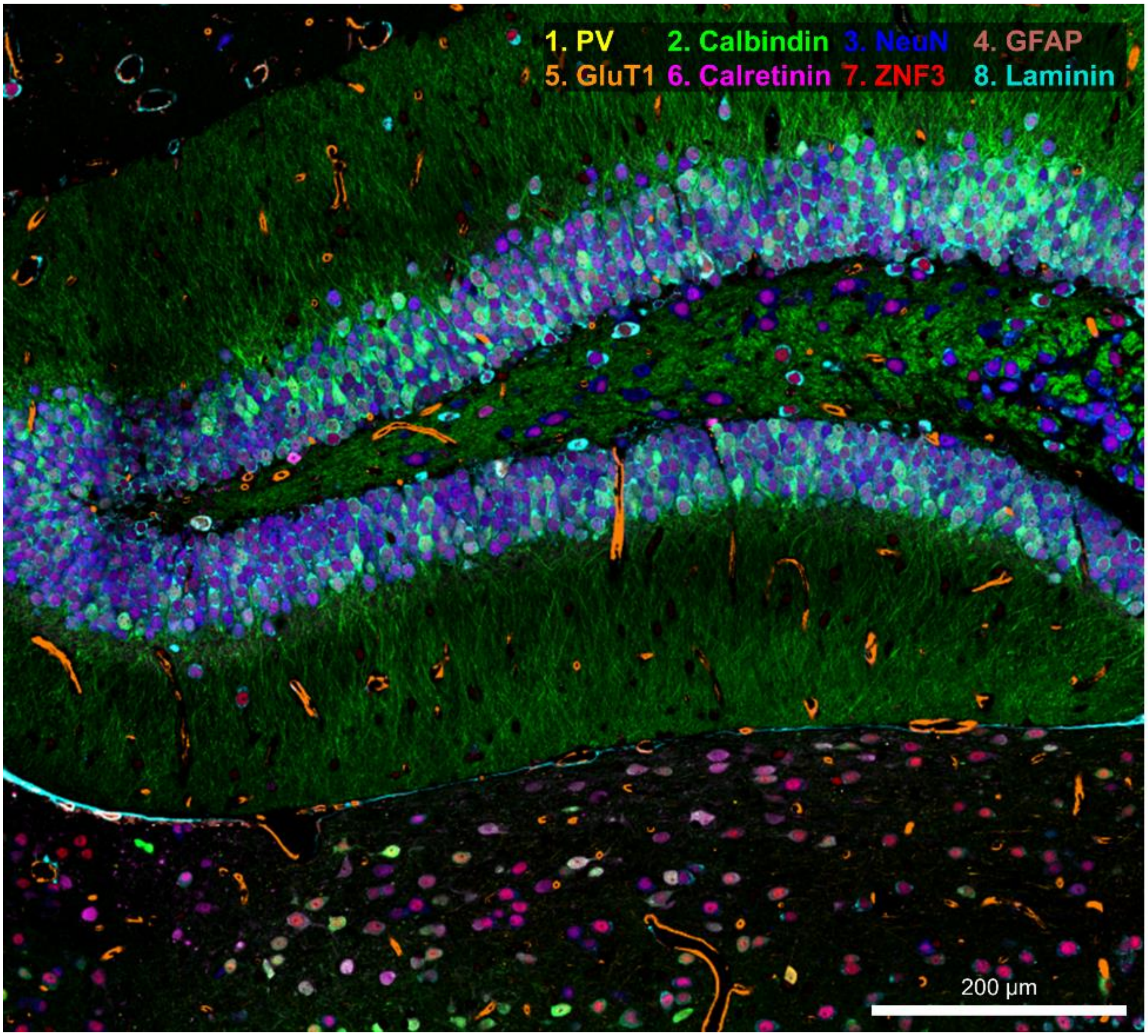


**Supplementary Figure 18**. **An enlarged image of Figure 4a**.

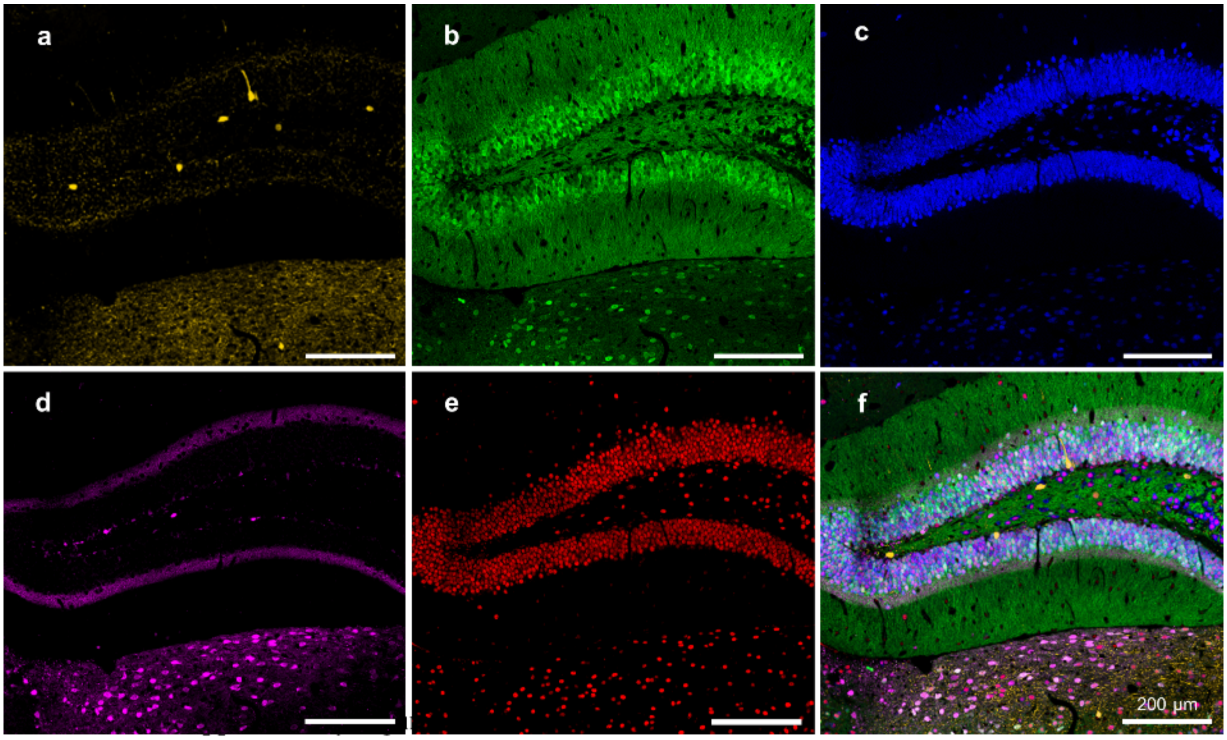


**Supplementary Figure 19**. **Five single-channel images of Figure 4a**. (**a–e**) Single-channel counterparts of the 8-colour multiplexed image shown in **Fig. 4a** clearly showing the spatial distribution of each protein in the mouse dentate gyrus and thalamus. (**a**) Yellow: parvalbumin (PV). (**b**) Green: calbindin. (**c**) Blue: NeuN. (**d**) Magenta: calretinin. (**e**) Red: zinc-finger protein 3 (ZNF3). (**f**) Merged image.

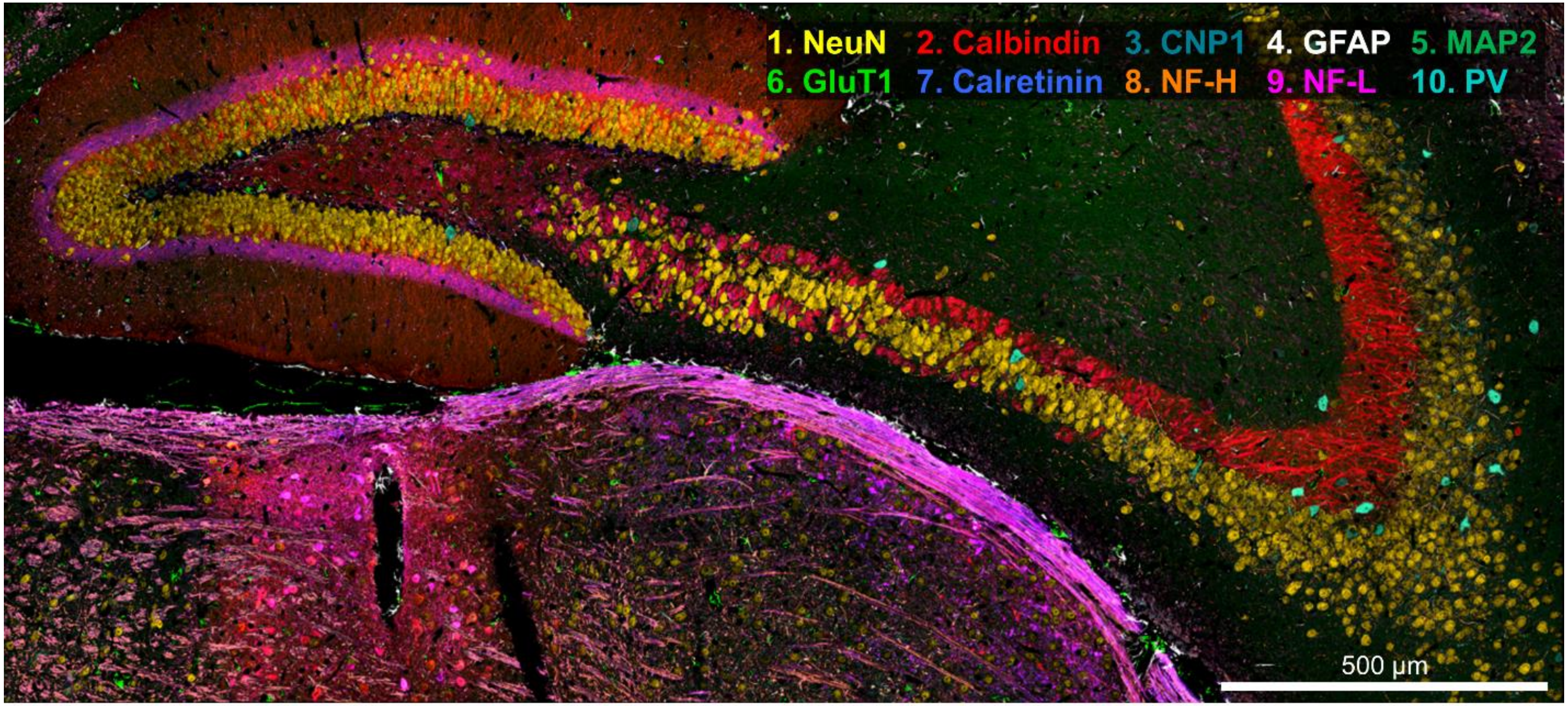


**Supplementary Figure 20**. **An enlarged image of Figure 4n**.

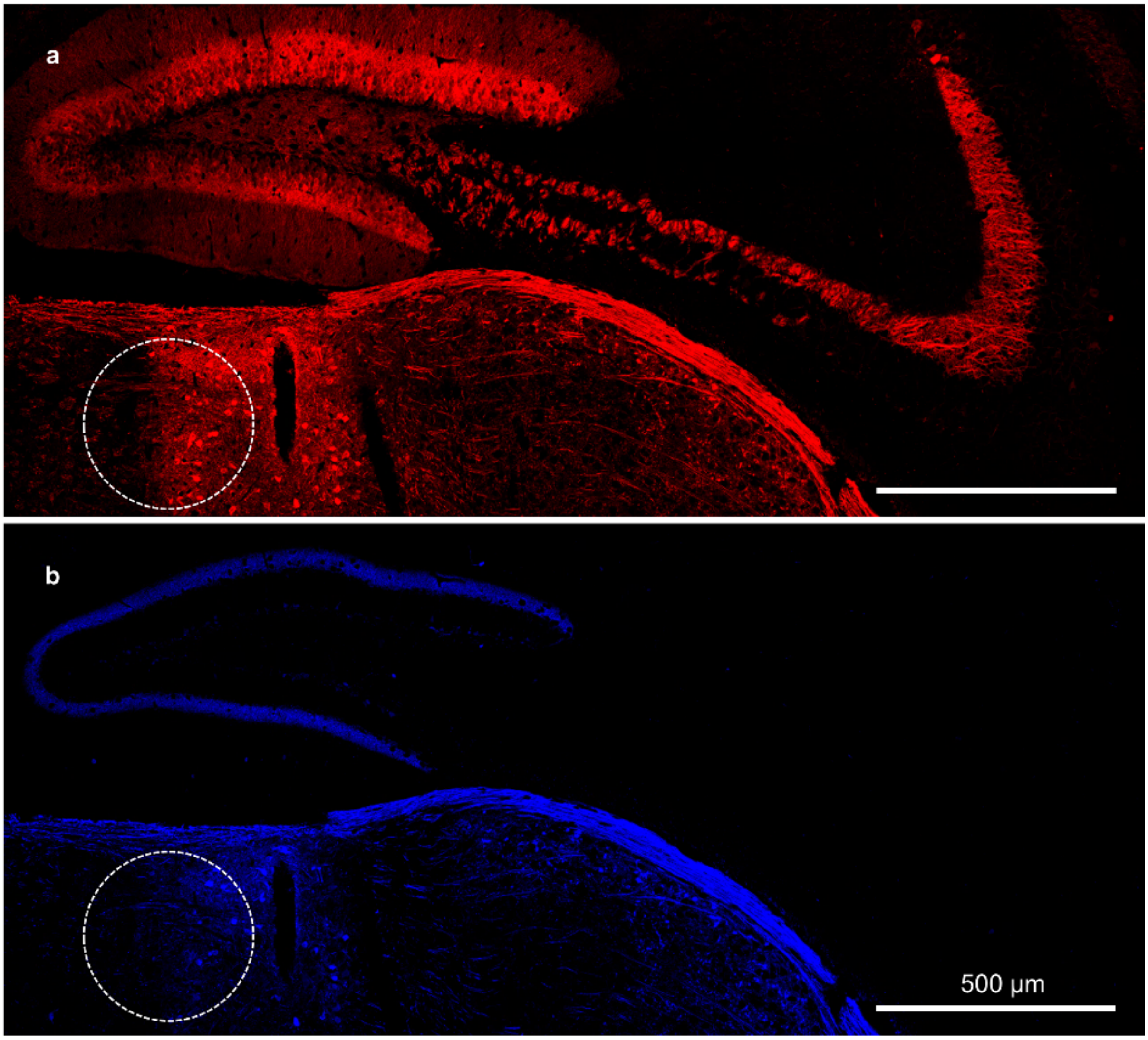


**Supplementary Figure 21**. **Two single-channel images of Figure 4n, showing calbindin and calretinin**. Dotted circles indicate the boundary between the lateral posterior nucleus of the thalamus and the anterior pretectal nucleus of the midbrain. Calbindin and calretinin were highly expressed in the lateral posterior nucleus. (**a**) Red: calbindin. (**b**) Blue: calretinin.

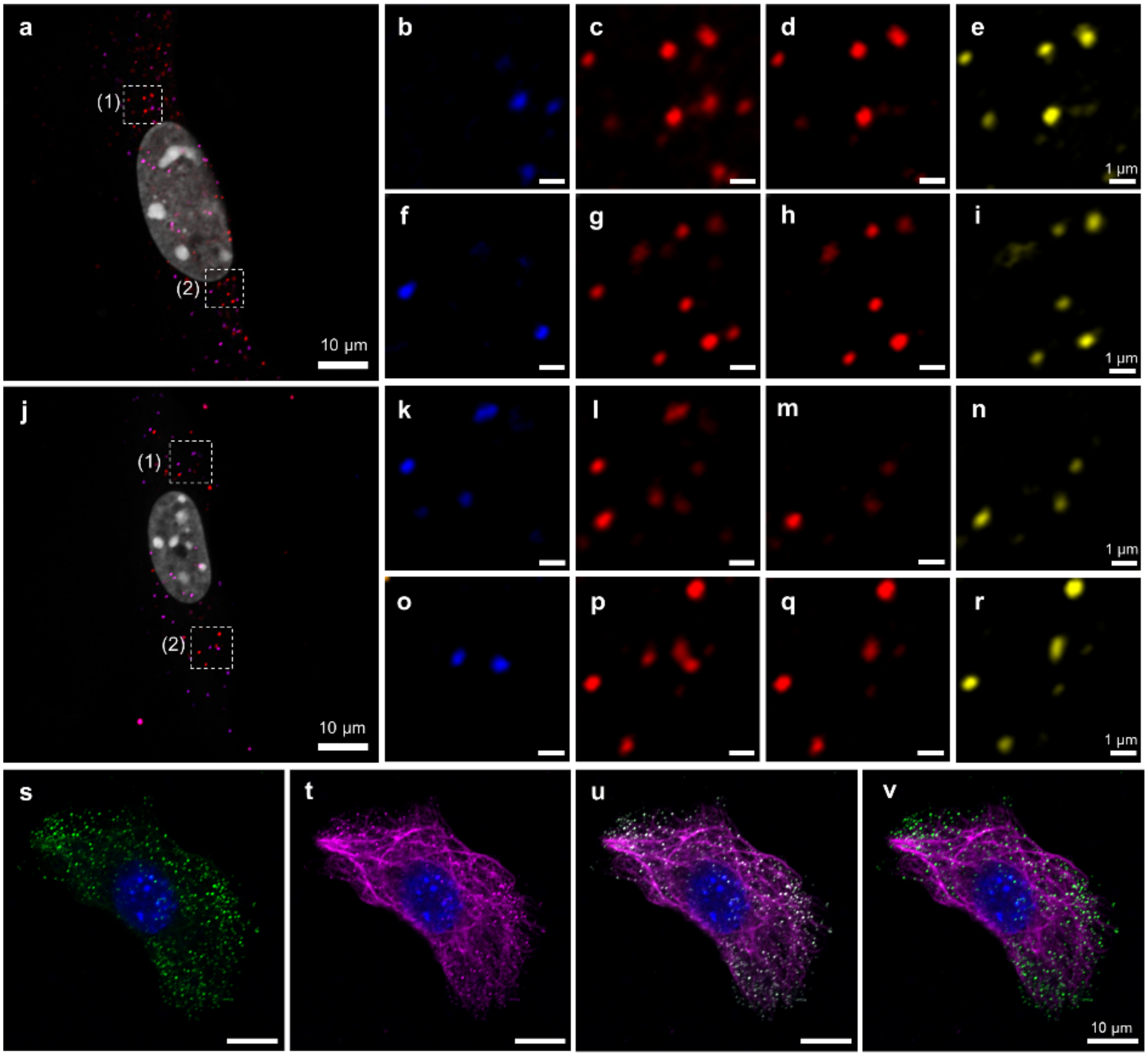


**Supplementary Figure 22**. **Application of PICASSO to multiplexed mRNA imaging and simultaneous imaging of mRNA and protein**. (**a–r**) Multiplexed imaging of two mRNAs with a single excitation laser *via* PICASSO and its validation. *Eif1a* and *Polr2a* mRNAs were labelled with hybridization chain reaction (HCR) probes conjugated to two spectrally overlapping fluorophores. *Polr2a* mRNA was labelled with another HCR probe conjugated to a spectrally distinct fluorophore, and its image was used as ground truth. (**a**) A merged image containing three channels, including two mixed channels (blue and red) and DAPI (white). (**b–e**) Magnified views of the white dotted box (1) in **a**. (**b**) Image acquired at the first detection channel, showing only *EiF1a* mRNA. (**c**) Image acquired at the second detection channel, showing both *Eif1a* and *Polr2a* mRNA before unmixing. (**d**) Second channel after unmixing *via* GS orthogonalization, showing only *Polr2a* mRNA. (**e**) Ground-truth image of *Polr2a* mRNA. Note that the *Polr2a* mRNA puncta shown in **d** (after unmixing) and **e** (ground truth) coincide. (**f–i**) As in (**b–e**) but for the white dotted box (2). (**j–r**) As in (**a–i**) but showing a different cell. (**s–v**) Simultaneous imaging of mRNA and protein with a single excitation laser *via* PICASSO. Cultured NIH-3T3 cells were stained with an antibody against vimentin and with an RNAscope fluorescent in-situ hybridization (FISH) probe against *Gapdh* mRNA. The anti-vimentin antibody and the FISH probe bore spectrally overlapping fluorophores. (**s**) Image acquired at the first detection channel, showing only *Gapdh* mRNA. (**t**) Image acquired at the second detection channel, showing both *Gapdh* mRNA and vimentin before unmixing. (**u**) A merged image combining **s** and **t**. (**v**) Merged image after unmixing *via* PICASSO. Magenta, vimentin; green, *Gapdh* mRNA.

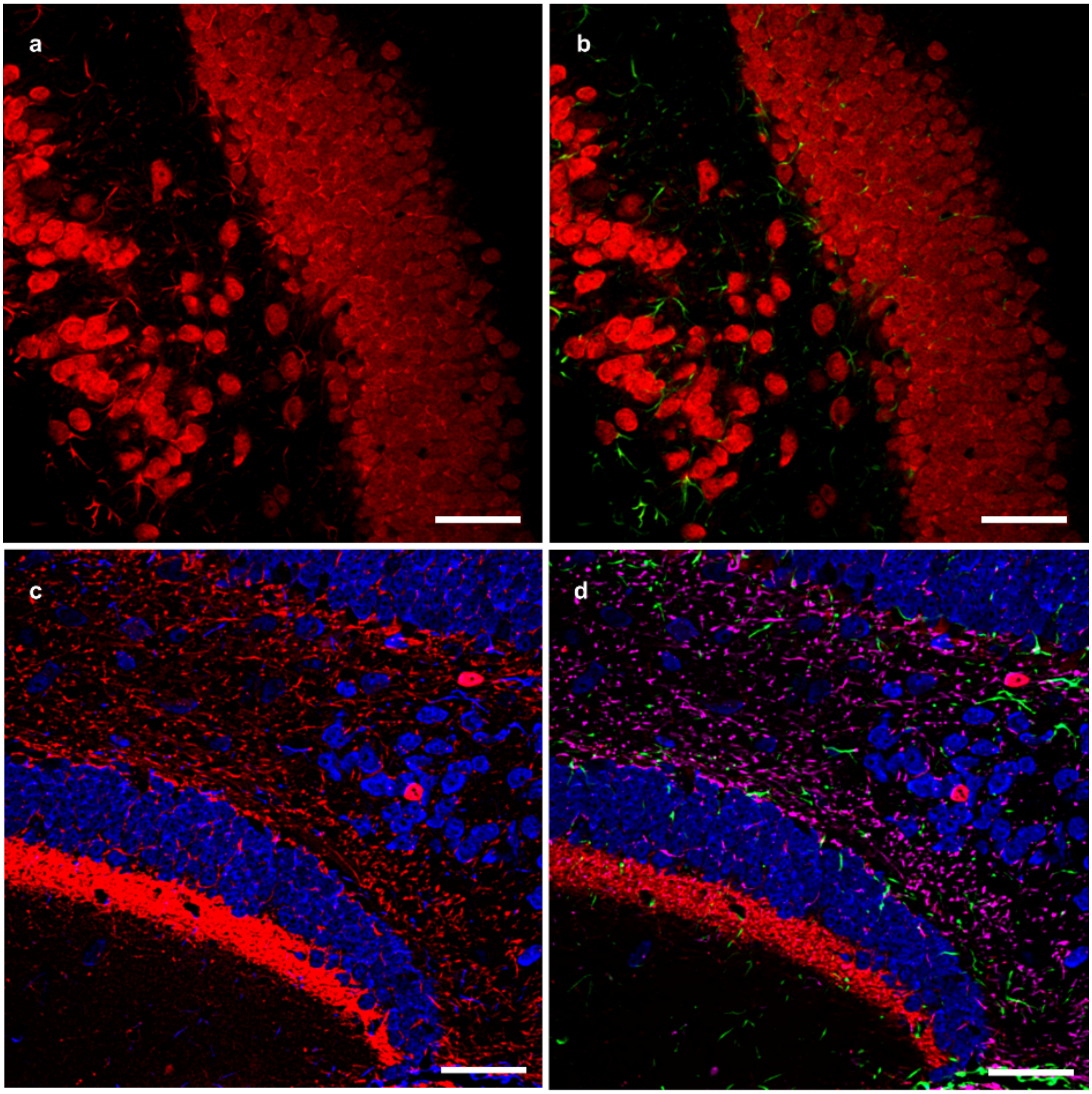


**Supplementary Figure 23**. **Application of PICASSO to expansion microscopy (ExM) and SHIELD**. (**a–b**) Two-colour multiplexed imaging of an ExM-processed mouse brain slice with one excitation laser *via* PICASSO. (**a**) Image acquired at the second detection channel, showing both NeuN and GFAP before unmixing. (**b**) Merged two-channel image after unmixing *via* MI minimization. Red, NeuN; green, GFAP. Two-colour multiplexed imaging of an expanded specimen was achieved *via* unmixing of two spectrally overlapping fluorophores via PICASSO. (**c**,**d**) Four-colour multiplexed imaging of a SHIELD-processed mouse brain slice imaged with two excitation lasers *via* PICASSO. (**c**) Merged image of two channels, each of which was acquired at the second detection channel under the illumination of different excitation lasers before unmixing. Blue, NeuN and GFAP, acquired with a 488-nm excitation laser; Red, calretinin and NF-H, acquired with a 557-nm laser. The images acquired at the first detection channel are not shown here. (**d**) After unmixing. Blue, NeuN; green, GFAP; red, calretinin; magenta, NF-H. Four-colour multiplexed imaging of a cleared brain was achieved *via* unmixing of four spectrally overlapping fluorophores via PICASSO. Scale bars: 50 µm.

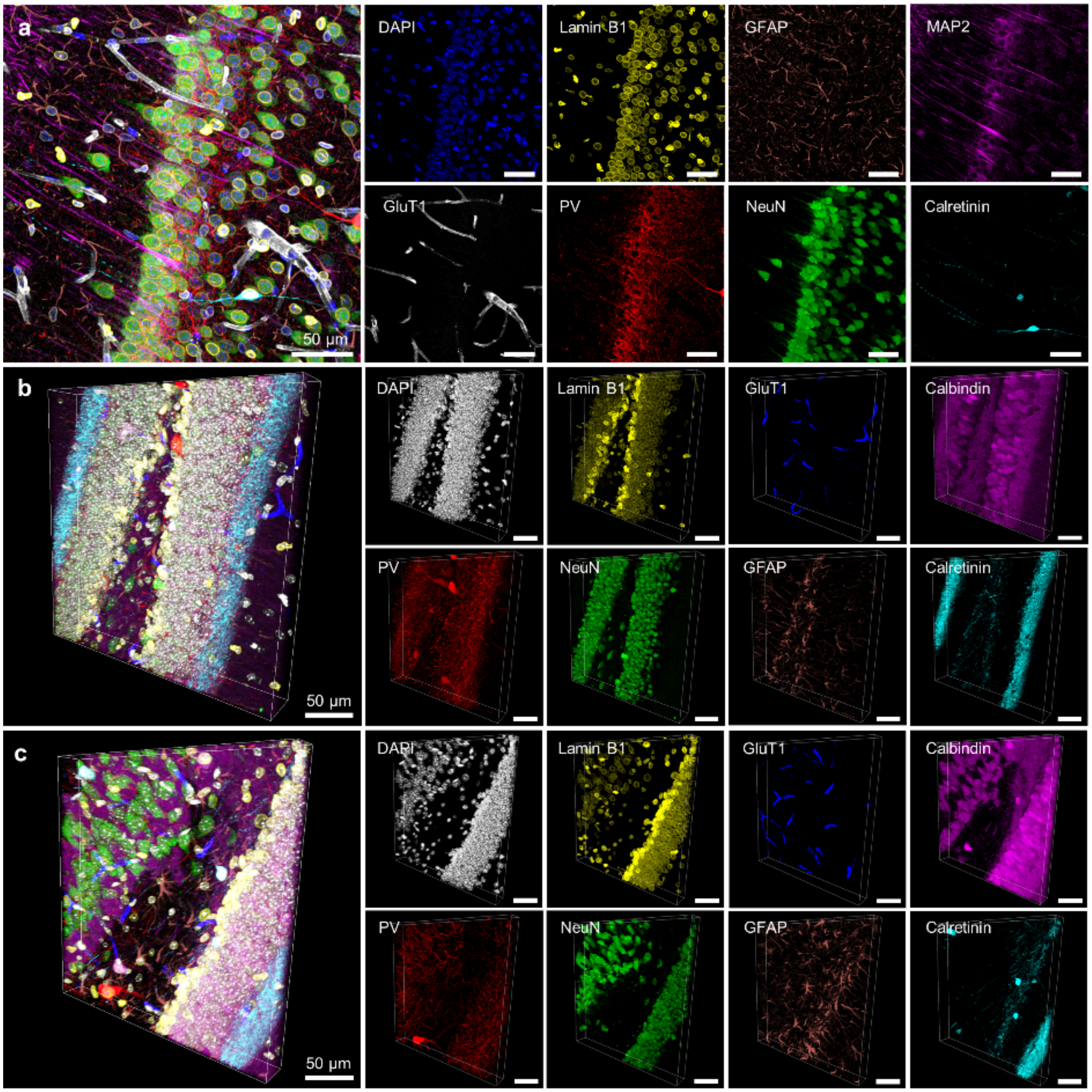


**Supplementary Figure 24**. **3D multiplexed imaging *via* MI minimization**. (**a–c**) 8-colour 3D multiplexed imaging of the mouse hippocampus. Seven preformed rabbit antibody complexes were used along with DAPI. (**a**) Maximum intensity projection (MIP) of the 8-colour multiplexed z-stack image from **Fig**. **5a**. Individual MIPs are presented on the right side. (**b–c**) The 8-colour 3D images acquired from the dentate gyrus in a different mouse brain slice. The z-stack images were acquired over a thickness of 40 μm with a step size of 0.5 μm.

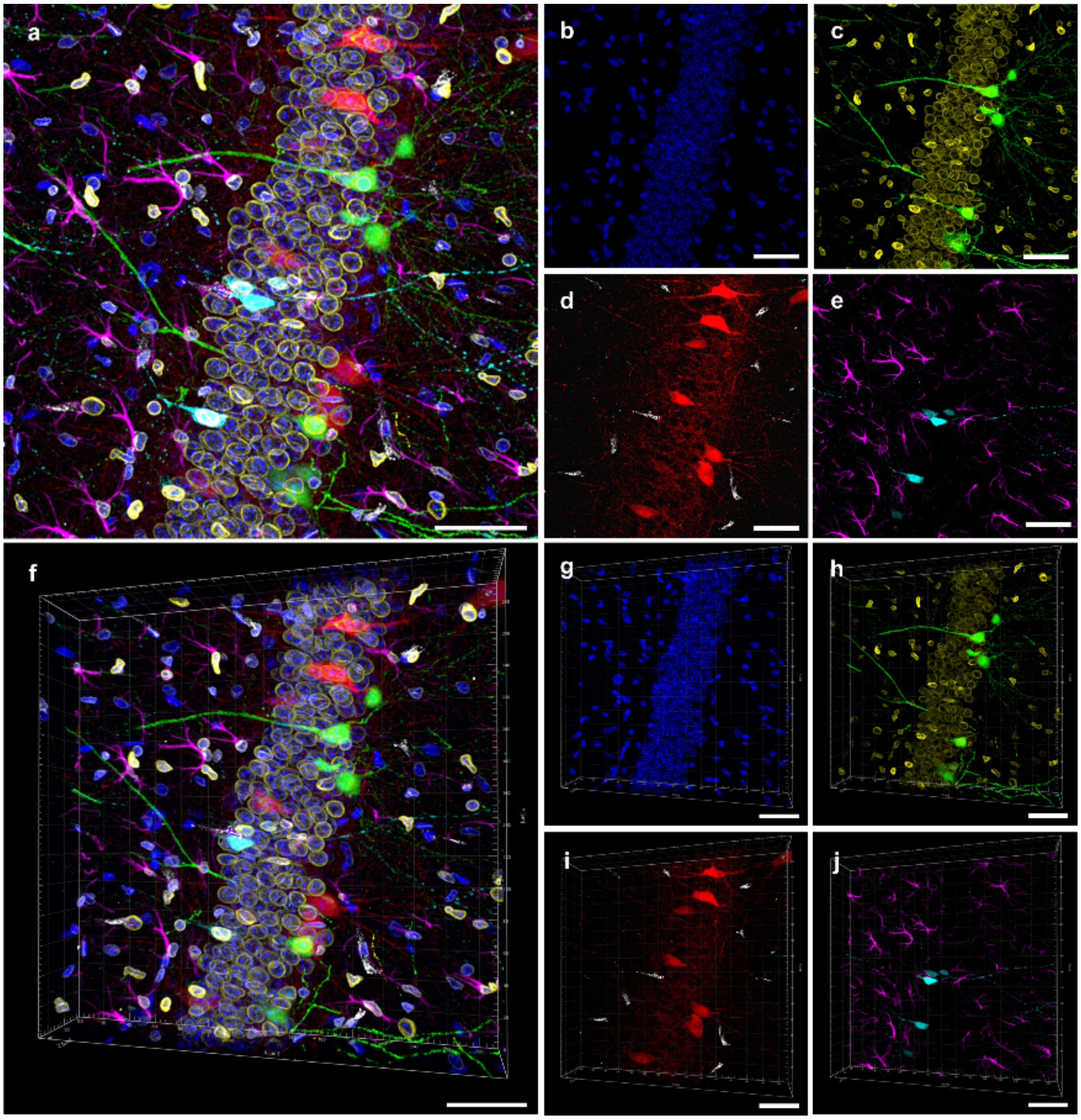


**Supplementary Figure 25**. **3D multiplexed imaging of a transgenic mouse brain slice *via* PICASSO**. Six proteins (including an endogenous YFP) and DAPI were imaged from a Thy1-YFP mouse brain slice at seven detection channels and unmixed *via* MI minimization. Five preformed antibody complexes were used. YFP was not labelled; instead, an innate YFP signal was imaged. (**a–e**) Maximum intensity projection of a z-stack image. (**a**) Merged image. (**b**) DAPI (blue), acquired using a 405-nm laser. (**c**) Lamin B1 (yellow) and YFP (green), acquired using a 488-nm laser. The signal from a CF488A-labelled preformed antibody complex against lamin B1 was acquired at the first detection channel and the CF488A + YFP signal at the second detection channel. The two images were unmixed *via* MI minimization. (**d**) PV (red) and GluT1 (white), acquired using a 557-nm laser. (**e**) GFAP (magenta) and calretinin (cyan), acquired using a 640-nm laser. (**f–j**) A 3D view of the z-stack image shown in **a–e**. Scale bars: 50 μm.

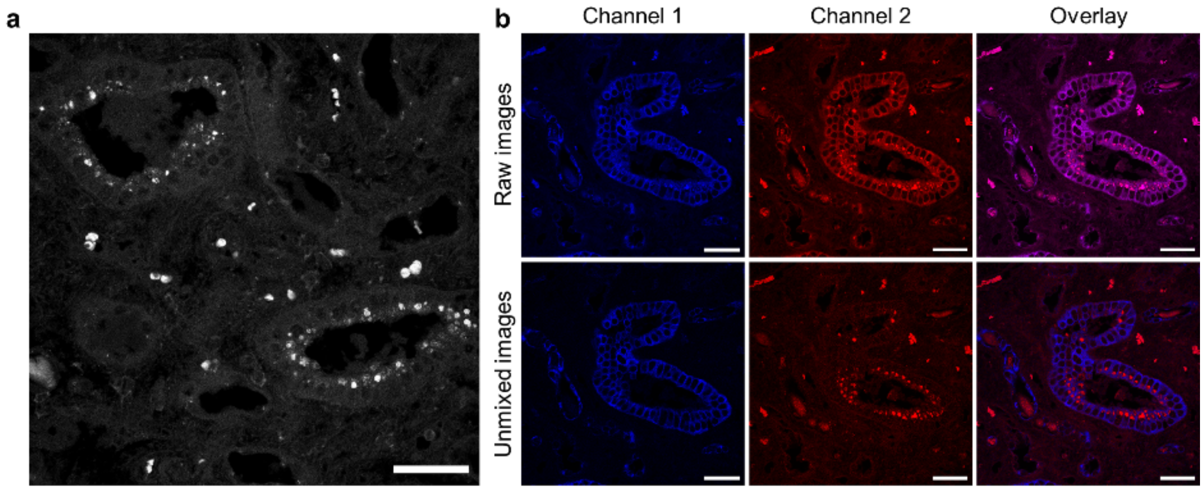


**Supplementary Figure 26**. **Removal of the autofluorescence from FFPE human kidney medulla *via* PICASSO**. (**a**) Image showing the autofluorescence of the specimen we used. (**b**) Removal of the autofluorescence *via* PICASSO. (**b**) FFPE sample was labelled with a preformed antibody complex against keratin (CF488A). Autofluorescence was considered as a separate fluorophore to remove it from the CF488A signal. Two images were acquired at two detection channels (490 – 525 nm and 525 – 560 nm) and then unmixed *via* PICASSO. Raw images were shown in the first row, and unmixed images were shown in the second row. In the second row, the blue signal is keratin (CF488A), and the red signal is the autofluorescence. Note that the unmixed autofluorescence signal is similar to the autofluorescence signal acquired in a separate specimen shown in **a**.

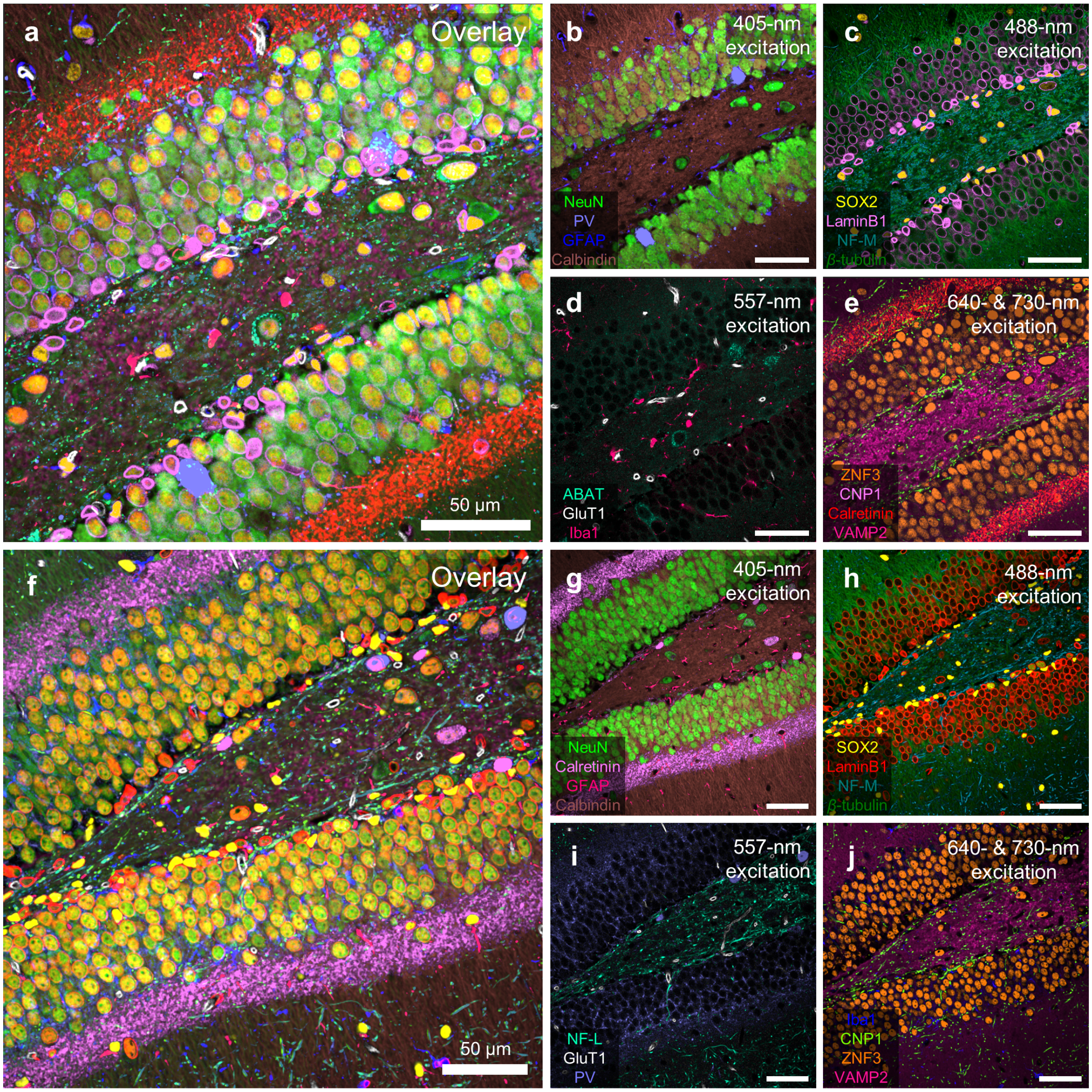


**Supplementary Figure 27**. **More demonstrations of 15-colour multiplexed imaging**. 15-colour multiplexed images acquired from the dentate gyrus of the mouse hippocampus of different mouse brain slices. To confirm that the 15-colour multiplexed imaging works reproducibly, different antibody combinations were used in this experiment from that used in **Fig. 1e**. Detailed information about the antibodies are shown in **Supplementary Table 6**. (**a**) Overlaid image. (**b–e**) Unmixed images acquired from each excitation laser. (**f–j**) Another 15-colour multiplexed imaging of a different antibody combination. (**f**) Overlaid image. (**g–j**) Unmixed images acquired from each excitation laser.

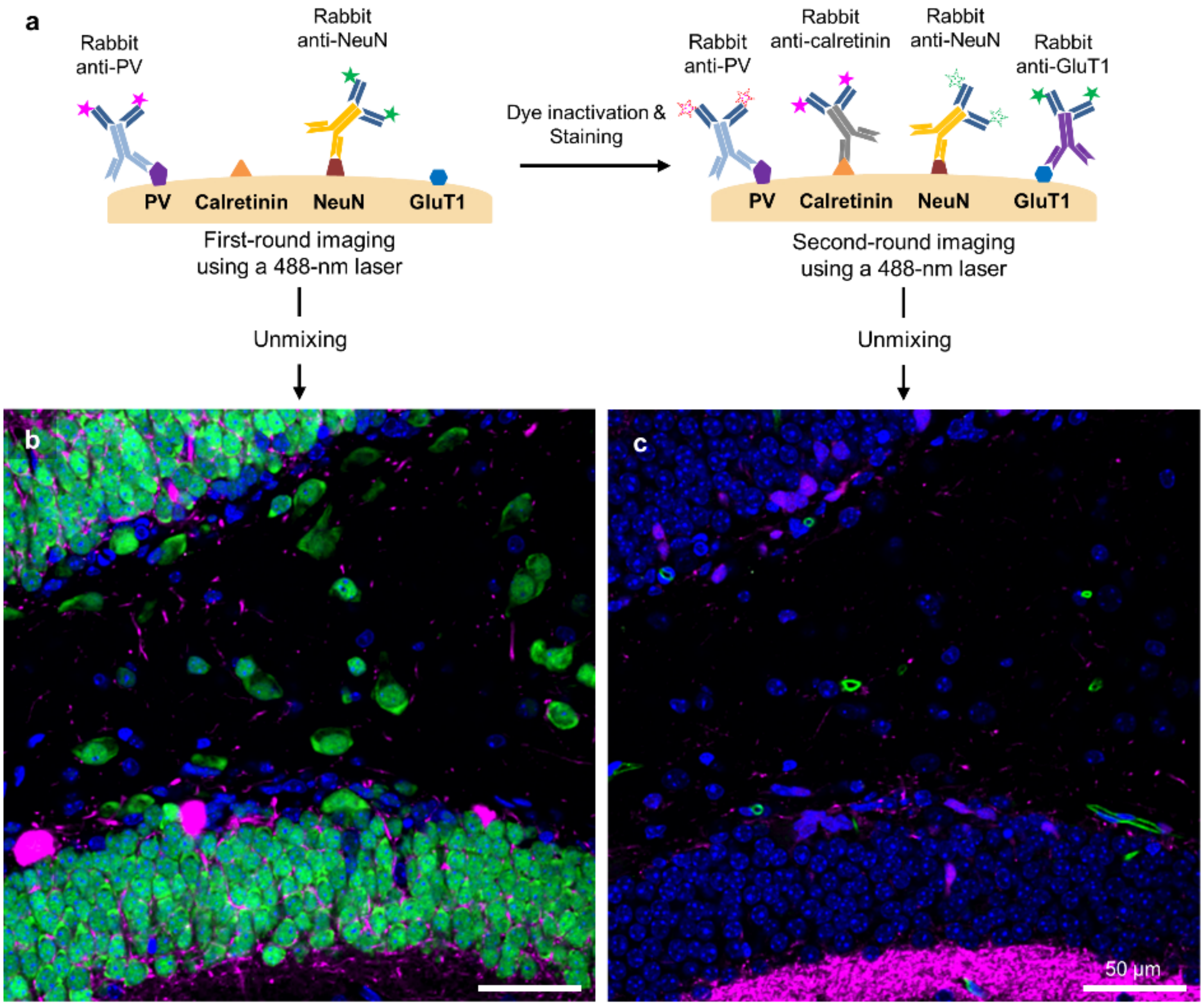


**Supplementary Figure 28**. **Demonstration of a higher multiplexing capability from combining PICASSO with a cyclic immunofluorescence technique (t-CyCIF)**. (**a**) Schematic showing how PICASSO can be combined with t-CyCIF. (**b–c**) Confocal microscopy of the hippocampal region of a 150 μm-thick mouse brain slice. (**b**) In the first cycle, PV and NeuN were labelled with spectrally overlapping fluorophores and imaged using a 488-nm excitation laser. The acquired images were then unmixed *via* MI minimization. Blue, DAPI; magenta, PV; green, NeuN. (**c**) After dye inactivation, calretinin and GluT1 were labelled with the same spectrally overlapping fluorophores and imaged using a 488-nm excitation laser. The acquired images were then unmixed *via* MI minimization. Blue, DAPI; magenta, calretinin; green, GluT1. Scale bars: 50 μm.

**Supplementary Note 1**. **Dependence of the emission spectra of fluorophores on optical, chemical, and environmental factors**. The internal optics of microscopes[1], such as the presence of a notch filter[2] and wavelength-dependent quantum efficiency of a microscope detector[3], strongly affect the emission spectra of fluorophores. In addition, some optical components are affected by environmental conditions; for example, the cut-off spectra of bandpass filters are sensitive to temperature changes due to the expansion or contraction of the coating materials[4]. Photomultiplier tube (PMT) detectors are also sensitive to temperature due to changes in the cathode sensitivity, especially at long wavelengths[5]. The chemical properties of solvents also affect emission spectra, such as the solvent polarity[6,7], pH[8,9], temperature[6], viscosity[6], internal charge transfer[6], and hydrogen bonding between fluorophore molecules and solvents[6]. Such solvent effects are becoming more important as diverse solvents such as dibenzyl ether, ethanol benzyl, iohexol, nicotinamide, antipyrine, sorbitol, N-methylacetamide, urea, DMSO, diatrizoic acid, n-methyl-d-glucamine, and deionized water have been introduced to recently developed tissue clearing, expansion, and shrinking techniques[10]. In addition, the emission spectra of fluorophores also depend on the local micro-environment inside cells or tissue slices[11–14]. The emission spectra also change when the fluorophores are conjugated to probes (e.g. antibodies)[15]. The mixing matrix, $M$, is the result of all of the above-mentioned optical, chemical, and environmental effects combined.

**Supplementary Note 2**. **Unmixing of two spectrally overlapping fluorophores *via* MI minimization**. The experimental design we used to validate the unmixing of two mixed images *via* MI minimization is shown in **Supplementary Fig**. **11**. We experimentally validated the unmixing of two spectrally overlapping fluorophores *via* MI minimization and demonstrated the two- or three-colour multiplexed imaging with a single excitation laser, as shown in **Supplementary Fig**. **12**. To quantitatively measure the accuracy of unmixing *via* MI minimization, we used a numerical simulator to generate synthetic mixed images with a known $\alpha_{gt}$ value and compared the $\alpha_{est}$ value estimated *via* MI minimization with the ground-truth $\alpha_{gt}$. The $\alpha$ estimation error was less than 1% for all values of $\alpha_{gt}$ (**Supplementary Fig. 13**). We also validated that unmixing *via* MI minimization can unmix two images even when there is a high level of mutual information between them, as shown in **Supplementary Fig**. **14**. Such high accuracy of unmixing *via* MI minimization would be attributed to the estimation of a single $\alpha$ value from more than millions of pixel values. A practical limiting factor of all spectral unmixing techniques, including PICASSO, is the shot noise, but it can be mitigated by averaging images multiple times or increasing the exposure time (**Supplementary Fig**. **15**).

In addition to the MI minimization, we demonstrated the second approach to estimate $\alpha$ based on Gram-Schmidt (GS) orthogonalization. GS orthogonalization, which is again a blind unmixing approach, decomposes the second image into two components: one parallel to the first image and the other perpendicular to the first image. In this approach, $\alpha_{\mathrm{GS}}$ was directly estimated by computing $\alpha_{\mathrm{GS}} = \frac{<\overrightarrow{IMG1},\overrightarrow{IMG2}>}{<\overrightarrow{IMG1},\overrightarrow{IMG1}>}$, where $<,>$ denotes a dot product, and $\overrightarrow{IMG1}$ and $\overrightarrow{IMG2}$ are $IMG1$ and $IMG2$ rearranged as vectors, respectively. Similarly, the image of the second fluorophore was obtained by calculating $IMG2 - \alpha_{\mathrm{GS}} \times IMG1$ (**Supplementary Fig. 16**). GS-based blind unmixing is more straightforward than MI minimization, but it tends to overestimate $\alpha$ when the expression patterns of two molecules highly overlap.

### Supplementary Note 3. PICASSO unmixing algorithm

---

**Algorithm 1**: PICASSO unmixing algorithm

---

**Input:** $D \in \mathbb{R}^{m\times n}$

**Output:** $X \in \mathbb{R}^{m\times n}$

**initialize** $X = D$

**while** *not converged* **do**

    **construct** M as $n \times n$ identity matrix

    **for all** $(i,j)$ such that $0 \le i < N, 0 \le j < N, i \ne j$ **do**

        **calculate** $\alpha_{i,j} = arg\min_{\alpha} I\left(q\left(bin(X_i)\right); q\left(bin(X_i - \alpha X_{ji})\right)\right)$

        $M[j,i] = \alpha_{i,j}$

    **end**

    **update** $X \leftarrow MX$

**end**

**return** $X$

*bin* denotes image binning function
*q* denotes image quantization function

---

Here, we find the condition to guarantee the strict increase of the relative portion of the dominant channel in each iteration. For the sake of simplicity, we will consider the first iteration of the unmixing process when $N = 2$. The image formation (linear mixing) process can be expressed as follows:

$$\begin{bmatrix} D_1 \\ D_2 \end{bmatrix} = \begin{bmatrix} 1 & \alpha_{1,2} \\ \alpha_{2,1} & 1 \end{bmatrix} \begin{bmatrix} F_1 \\ F_2 \end{bmatrix} = \mathrm{M} \begin{bmatrix} F_1 \\ F_2 \end{bmatrix},$$

where $\alpha_{i,j}$ refers to the relative leakage from the $j_{th}$ fluorophore to the $i_{th}$ image; $D_i$ and $F_i$ are the $i_{th}$ channels of the acquired and fluorophore images, respectively.

Without losing generality, let's consider $i = 1$ and write the unmixing equation:

$$X_{1(1)} = D_1 - \gamma \alpha_{1,2(0)} D_2.$$

By substitution ($\varepsilon = \gamma \alpha_{1,2(0)}$), we obtain the following equation, where $\alpha$ is a positive real number:

$$X_{1(1)} = D_1 - \alpha D_2.$$

Writing $X_{1(1)}$ as a linear summation of $F_i$ gives:

$$X_{1(1)} = \left(F_1 + \alpha_{1,2} F_2\right) - \varepsilon\left(F_2 + \alpha_{2,1} F_1\right).$$

We can rearrange the equation as:

$$X_{1(1)} = \left(1 - \varepsilon \alpha_{2,1}\right) F_1 + \left(\alpha_{1,2} - \varepsilon\right) F_2.$$

The condition for the strict increase of the relative portion of the dominant channel can be written as:

$$\frac{1}{\alpha_{1,2}} < \frac{1 - \varepsilon \alpha_{2,1}}{\alpha_{1,2} - \varepsilon}.$$

As the update parameter $\gamma$ can be chosen to be arbitrarily small (and hence $\varepsilon$ can be arbitrarily small), all terms are positive, and the inequality becomes as follows:

$$\alpha_{1,2} - \varepsilon < \alpha_{1,2}\left(1 - \varepsilon \alpha_{2,1}\right)$$

$$\therefore \varepsilon\left(1 - \alpha_{1,2} \alpha_{2,1}\right) > 0$$

$$\therefore 1 - \alpha_{1,2} \alpha_{2,1} > 0$$

$$\therefore \det(M) > 0.$$

Therefore, during the first iteration, the strict increase of the relative portion of the dominant channel is guaranteed if the determinant of the mixing matrix is positive. After the first iteration, $\alpha_{1,2}$ and $\alpha_{2,1}$ become smaller, so $1 - \alpha_{1,2} \alpha_{2,1} > 0$ is still met.

The same analogy can be extended to $N = 3$ and further:

$$\begin{bmatrix} D_1 \\ D_2 \\ D_3 \end{bmatrix} = \begin{bmatrix} 1 & \alpha_{1,2} & \alpha_{1,3} \\ \alpha_{2,1} & 1 & \alpha_{2,3} \\ \alpha_{3,1} & \alpha_{3,2} & 1 \end{bmatrix} \begin{bmatrix} F_1 \\ F_2 \\ F_3 \end{bmatrix} = \mathrm{M} \begin{bmatrix} F_1 \\ F_2 \\ F_3 \end{bmatrix}.$$

Again, without losing generality, let's write the unmixing equation for $i = 1$:

$$X_{1(1)} = X_{1(0)} - \sum_{i \neq j} \gamma \alpha_{1,j(0)} X_{j(0)} = D_1 - \sum_{i \neq j} \gamma \alpha_{1,j(0)} D_j = D_1 - \gamma \alpha_{1,2(0)} D_2 - \gamma \alpha_{1,3(0)} D_3.$$

By substitution ($\varepsilon = \gamma \alpha_{1,2(0)}$ and $\delta = \gamma \alpha_{1,3(0)}$), we can re-write as follows, where $\alpha$ and $\beta$ are both positive real numbers:

$$X_{1(1)} = D_1 - \varepsilon D_2 - \delta D_3.$$

If each subtraction increases the relative portion of the dominant channel, then performing all subtractions will also strictly increase the relative portion of the dominant channel. Therefore, it suffices to consider two cases separately: 1) $\varepsilon > 0$ and $\delta = 0$ and 2) $\varepsilon = 0$ and $\delta > 0$.

From 1), writing $X_{1(1)}$ as a linear summation of $F_i$ gives:

$$X_{1(1)} = \left(F_1 + \alpha_{1,2} F_2 + \alpha_{1,3} F_3\right) - \varepsilon\left(F_2 + \alpha_{2,1} F_1 + \alpha_{2,3} F_3\right)$$

$$\therefore X_{1(1)} = \left(1 - \varepsilon \alpha_{2,1}\right) F_1 + \left(\alpha_{1,2} - \varepsilon\right) F_2 + \left(\alpha_{1,3} - \varepsilon \alpha_{2,3}\right) F_3.$$

The condition for the strict increase of the relative portion of the dominant channel can be written as:

$$\frac{1}{\alpha_{1,2}} < \frac{1 - \varepsilon \alpha_{2,1}}{\alpha_{1,2} - \varepsilon} \text{ and } \frac{1}{\alpha_{1,3}} < \frac{1 - \varepsilon \alpha_{2,1}}{\alpha_{1,3} - \varepsilon \alpha_{2,3}}$$

$$\therefore \varepsilon\left(1 - \alpha_{1,2} \alpha_{2,1}\right) > 0 \text{ and } \varepsilon\left(\alpha_{2,3} - \alpha_{1,3} \alpha_{2,1}\right) > 0$$

$$\therefore \left(1 - \alpha_{1,2} \alpha_{2,1}\right) > 0 \text{ and } \left(\alpha_{2,3} - \alpha_{1,3} \alpha_{2,1}\right) > 0.$$

We find that both terms of the left-hand side are the determinants of the $2 \times 2$ sub-matrices of the original mixing matrix $M$.

From 2), similarly, we also obtain that the determinants of the other $2 \times 2$ sub-matrices of the original mixing matrix $M$ need to be positive. In extension, the determinants of all possible $2 \times 2$ sub-matrices of the mixing matrix need to be positive to guarantee the strict increase of the relative portion of the dominant channel.

**Supplementary Note 4. Biological contexts of Fig. 4a.** Calbindin is a calcium-binding protein, and its spatial expression level in the mouse brain was found to be high in the dentate gyrus (DG) along the mossy fiber projections and relatively low in the thalamus (TH) (**Fig. 4a** and **Supplementary Fig. 19b**), consistent with the literature[16]. Calretinin is a calcium-binding protein that is highly expressed in the mammalian brain[17]. Calretinin expression was found to be high in a thin layer of the molecular layer (DG-mo) and granule cell layer (DG-sg) of the DG and TH (**Fig. 4a** and **Supplementary Fig. 19d**) and matched well with the database (row 1 of **Supplementary Table 4**). ZNF3 is a transcription factor that is highly expressed in the mouse brain (row 2 of **Supplementary Table 4**). Consistent with the database, ZNF3 was found to be highly expressed in the DG-sg and polymorph layer (DG-po) of the DG and TH below the DG, as shown in **Fig. 4a** and **Supplementary Fig. 19e**. NeuN is a neuron marker that is highly expressed exclusively only in the brain (row 3 of **Supplementary Table 4**). As shown in **Fig. 4a** and **Supplementary Fig. 19c**, NeuN was found to be expressed in DG-sg, DG-po, and TH, but its expression was higher in DG than in TH, consistent with the literature[18]. Parvalbumin (PV) is a calcium-binding albumin protein that is highly expressed in the mouse brain (row 4 of **Supplementary Table 4**). PV was found to be widely expressed in DG-po, DG-sg, DG-mo, and TH (**Fig. 4a** and **Supplementary Fig. 19a**), again consistent with the literature[19].

**Supplementary Video 1. Sample preparation procedure for the 10-colour multiplexed imaging of a mouse brain slice.**

**Supplementary Video 2. 3D 8-colour multiplexed imaging of a mouse brain slice.** See **Fig. 5a** for details. Blue, DAPI; yellow, lamin B1; brown, GFAP; magenta, MAP2; white, GluT1; red, PV; green, NeuN; cyan, calretinin.

**Supplementary Video 3. Acquisition of the 10-colour multiplexed image.** A mouse brain slice stained with 10 preformed antibody complexes conjugated with 10 spectrally overlapping fluorophores was imaged by using a spinning-disk confocal microscopy system.

| # | Antibody | Vendor | Catalog# | Host | Clonality |
|---|---|---|---|---|---|
| **Mouse brain marker** | | | | | |
| 1 | ABAT | HPA | HPA041690 | Rb | Poly |
| 2 | $\alpha$-internexin/NF66 | EnCor | RPCA-$\alpha$-Int | Rb | Poly |
| 3 | ARFGEF1 | HPA | HPA023822 | Rb | Poly |
| 4 | $\alpha$-tubulin | Abcam | ab18251 | Rb | Poly |
| 5 | $\beta$-tubulin | Abcam | ab6046 | Rb | Poly |
| 6 | Calbindin | Abcam | ab11426 | Rb | Poly |
| 7 | CALB2 | HPA | HPA007305 | Rb | Poly |
| 8 | Calretinin | Abcam | ab702 | Rb | Poly |
| 9 | CAMK2B | HPA | HPA026307 | Rb | Poly |
| 10 | CNP1 | SYSY | 355 002 | Rb | Poly |
| 11 | DDX3X | HPA | HPA001648 | Rb | Poly |
| 12 | E1F1AX | HPA | HPA002561 | Rb | Poly |
| 13 | FGF3 | HPA | HPA012692 | Rb | Poly |
| 14 | GABA-A receptor $\alpha$1 | SYSY | 224 203 | Rb | Poly |
| 15 | GAD65/67 | Millipore | AB1511 | Rb | Poly |
| 16 | GFAP | Abcam | ab7260 | Rb | Poly |
| 17 | GFAP | HPA | HPA056030 | Rb | Poly |
| 18 | GFP | Abcam | ab290 | Rb | Poly |
| 29 | GluR1 | Abcam | ab31232 | Rb | Poly |
| 20 | GPR17 | HPA | HPA029766 | Rb | Poly |
| 21 | Homer1 | SYSY | 160 003 | Rb | Poly |
| 22 | Iba1 | SYSY | 234 003 | Rb | Poly |
| 23 | INA | HPA | HPA008057 | Rb | Poly |
| 24 | LaminB1 | Abcam | ab16048 | Rb | Poly |

| | | | | | |
|---|---|---|---|---|---|
| 25 | Laminin | Abcam | ab11575 | Rb | Poly |
| 26 | Laminin | EnCor | RPCA-Laminin | Rb | Poly |
| 27 | LHX2 | HPA | HPA000838 | Rb | Poly |
| 28 | MAP2 | Abcam | ab32454 | Rb | Poly |
| 29 | MBP | Abcam | ab40390 | Rb | Poly |
| 30 | MBP | HPA | HPA049222 | Rb | Poly |
| 31 | MBP | Aves | MBP | Chk | Poly |
| 32 | NECAB2 | HPA | HPA013998 | Rb | Poly |
| 33 | NeuN | Millipore | ABN78 | Rb | Poly |
| 34 | Neurofilament 200 | Sigma | N4142 | Rb | Poly |
| 35 | Neurofilament-L | EnCor | RPCA-NF-L | Rb | Poly |
| 36 | Neurofilament-M | EnCor | RPCA-NF-M | Rb | Poly |
| 37 | Neurofilament-H | EnCor | RPCA-NF-H | Rb | Poly |
| 38 | Neuropeptide Y | Immunostar | 22940 | Rb | Poly |
| 39 | Parvalbumin | Abcam | ab11427 | Rb | Poly |
| 40 | Parvalbumin | Novus | NB120-11427 | Rb | Poly |
| 41 | PCP4 | HPA | HPA005792 | Rb | Poly |
| 42 | RAP1GAP | HPA | HPA001922 | Rb | Poly |
| 43 | SLC2A1 (= GluT1) | HPA | HPA031345 | Rb | Poly |
| 44 | SOX2 | SYSY | 347 003 | Rb | Poly |
| 45 | Somatostatin | HPA | HPA019472 | Rb | Poly |
| 46 | SV2A | Abcam | ab32942 | Rb | Poly |
| 47 | Synapsin 1 | Novus | NB300-104 | Rb | Poly |
| 48 | TBR1 | Abcam | ab31940 | Rb | Poly |
| 49 | TH | Abcam | ab112 | Rb | Poly |
| 50 | VAMP2 | abcam | ab3347 | Rb | Poly |
| 51 | vGluT1 | SYSY | 135 303 | Rb | Poly |
| 52 | vGluT2 | SYSY | 135 403 | Rb | Poly |

| | | | | | |
|---|---|---|---|---|---|
| 53 | ZNF3 | HPA | HPA003719 | Rb | Poly |
| **Cell marker** | | | | | |
| 54 | $\beta$-tubulin | Abcam | ab6046 | Rb | Poly |
| 55 | Clathrin heavy chain | Abcam | ab21679 | Rb | Poly |
| 56 | CollagenIV | Abcam | ab6586 | Rb | Poly |
| 57 | Fibronectin | Abcam | ab2413 | Rb | Poly |
| 58 | GM130 | Abcam | ab52649 | Rb | Mono |
| 59 | LaminB1 | Abcam | ab16048 | Rb | Poly |
| 60 | Pericentrin | Abcam | ab4448 | Rb | Poly |
| 61 | Vimentin | Abcam | ab45939 | Rb | Poly |
| **Human tissue marker** | | | | | |
| 62 | KRT19 | HPA | HPA002465 | Rb | Poly |
| 63 | COX IV | Abcam | ab16056 | Rb | Poly |
| 64 | Histone H3 | Abcam | ab1791 | Rb | Poly |
| 65 | Vimentin | Abcam | ab45939 | Rb | Poly |

**Supplementary Table 1. List of the tested primary antibodies.**

| # | Dye | Vendor | Catalog# |
|---|---|---|---|
| 1 | ATTO390 | ATTO-TEC | AD 390-31 |
| 2 | CF405S | Biotium | #92110 |
| 3 | CF405M | Biotium | #92111 |
| 4 | CF405L | Biotium | #92112 |
| 5 | Dylight405 | Life technologies | 46400 |
| 6 | ATTO430LS | ATTO-TEC | AD 430LS-31 |
| 7 | CF488A | Biotium | #92120 |
| 8 | ATTO488 | ATTO-TEC | AD 488-31 |
| 9 | Alexa Fluor 488 | Life technologies | A20000 |
| 10 | ATTO490LS | ATTO-TEC | AD 490LS-31 |
| 11 | ATTO514 | ATTO-TEC | AD 514-31 |
| 12 | CF514 | Biotium | #92103 |
| 13 | ATTO532 | ATTO-TEC | AD 532-31 |
| 14 | Alexa Fluor 546 | Life technologies | A20002 |
| 15 | ATTO565 | ATTO-TEC | AD 565-31 |
| 16 | CF568 | Biotium | #92131 |
| 17 | ATTO Rho101 | ATTO-TEC | AD Rho101-31 |
| 18 | Alexa Fluor 594 | JacksonImmunoResearch | 111-587-008 |
| 19 | ATTO594 | ATTO-TEC | AD 594-31 |
| 20 | ATTO633 | ATTO-TEC | AD 633-31 |
| 21 | CF633 | Biotium | #92133 |
| 22 | Alexa Fluor 647 | JacksonImmunoResearch | 111-607-008 |
| 23 | ATTO647N | ATTO-TEC | AD 647N-31 |
| 24 | Alexa Fluor 680 | JacksonImmunoResearch | 111-627-008 |
| 25 | CF660R | Biotium | #92134 |
| 26 | CF680R | Biotium | #92107 |
| 27 | ATTO725 | ATTO-TEC | AD 725-31 |
| 28 | Alexa Fluor 790 | JacksonImmunoResearch | 111-657-008 |

**Supplementary Table 2**. **List of the fluorophores that worked with the primary antibody–Fab preformation technique**.

| Channel # | Fluorophore | Emission peak (nm) | Abbreviation |
|---|---|---|---|
| Ch1 (645–655 nm) | CF633 | 650 | CF633 |
| | Alexa Fluor 633 | 647 | AF633 |
| Ch2 (663–673 nm) | Alexa Fluor 647 | 668 | AF647 |
| | ATTO647N | 669 | AT647N |
| Ch3 (678–688 nm) | CF660R | 683 | CF660R |
| | ATTO665 | 685 | AT665 |
| Ch4 (696–706 nm) | CF680R | 701 | CF680R |
| | Alexa Fluor 680 | 702 | AF680 |
| Ch5 (714–724 nm) | ATTO700 | 719 | AT700 |
| | Alexa Fluor 700 | 719 | AF770 |

| # | Ch 1 | Ch 2 | Ch 3 | Ch 4 | Ch5 | Correlation w/ GT | Determinants of all 2×2 matrices |
|---|---|---|---|---|---|---|---|
| 1 | CF633 | AF647 | CF660R | CF680R | AT700 | 0.990 | Positive |
| 2 | CF633 | AF647 | CF660R | CF680R | AF770 | 0.990 | Positive |
| 3 | CF633 | AF647 | CF660R | AF680 | AT700 | 0.989 | Positive |
| 4 | CF633 | AF647 | CF660R | AF680 | AF770 | 0.989 | Positive |
| 5 | CF633 | AF647 | AT665 | CF680R | AT700 | 0.990 | Positive |
| 6 | CF633 | AF647 | AT665 | CF680R | AF770 | 0.990 | Positive |
| 7 | CF633 | AF647 | AT665 | AF680 | AT700 | 0.990 | Positive |
| 8 | CF633 | AF647 | AT665 | AF680 | AF770 | 0.990 | Positive |
| 9 | CF633 | AT647N | CF660R | CF680R | AT700 | 0.991 | Positive |
| 10 | CF633 | AT647N | CF660R | CF680R | AF770 | 0.990 | Positive |
| 11 | CF633 | AT647N | CF660R | AF680 | AT700 | 0.990 | Positive |
| 12 | CF633 | AT647N | CF660R | AF680 | AF770 | 0.991 | Positive |
| 13 | CF633 | AT647N | AT665 | CF680R | AT700 | 0.990 | Positive |
| 14 | CF633 | AT647N | AT665 | CF680R | AF770 | 0.991 | Positive |
| 15 | CF633 | AT647N | AT665 | AF680 | AT700 | 0.990 | Positive |
| 16 | CF633 | AT647N | AT665 | AF680 | AF770 | 0.990 | Positive |
| 17 | AF633 | AF647 | CF660R | CF680R | AT700 | 0.989 | Positive |
| 18 | AF633 | AF647 | CF660R | CF680R | AF770 | 0.990 | Positive |

| | | | | | | | |
|---|---|---|---|---|---|---|---|
| 19 | AF633 | AF647 | CF660R | AF680 | AT700 | 0.990 | Positive |
| 20 | AF633 | AF647 | CF660R | AF680 | AF770 | 0.990 | Positive |
| 21 | AF633 | AF647 | AT665 | CF680R | AT700 | 0.990 | Positive |
| 22 | AF633 | AF647 | AT665 | CF680R | AF770 | 0.990 | Positive |
| 23 | AF633 | AF647 | AT665 | AF680 | AT700 | 0.990 | Positive |
| 24 | AF633 | AF647 | AT665 | AF680 | AF770 | 0.990 | Positive |
| 25 | AF633 | AT647N | CF660R | CF680R | AT700 | 0.990 | Positive |
| 26 | AF633 | AT647N | CF660R | CF680R | AF770 | 0.990 | Positive |
| 27 | AF633 | AT647N | CF660R | AF680 | AT700 | 0.990 | Positive |
| 28 | AF633 | AT647N | CF660R | AF680 | AF770 | 0.991 | Positive |
| 29 | AF633 | AT647N | AT665 | CF680R | AT700 | 0.991 | Positive |
| 30 | AF633 | AT647N | AT665 | CF680R | AF770 | 0.991 | Positive |
| 31 | AF633 | AT647N | AT665 | AF680 | AT700 | 0.990 | Positive |
| 32 | AF633 | AT647N | AT665 | AF680 | AF770 | 0.990 | Positive |

**Supplementary Table 3**. **Unmixing of 32 different fluorophore combinations.** We tested whether PICASSO could unmix various fluorophore combinations even when the same detection channels are used. For each of the five fluorophores we used in **Fig. 2** (CF633, Alexa Fluor 647, CF660R, CF680R, and ATTO700), we chose an additional fluorophore with a similar emission spectrum. We chose Alexa Fluor 633 for CF633, ATTO647N for Alexa Fluor 647, ATTO665 for CF660R, Alexa Fluor 680 for CF680R, and Alexa Fluor 700 for ATTO700 (first table). For these five pairs of fluorophores, we generated 32 (=$2^5$) different 5-colour combinations and calculated mixing matrices of these 32 combinations based on the reference emission spectra of the fluorophores. When calculating the mixing matrices, the five detection channels shown in **Fig. 2b** were used. We then synthesized mixed imaging by using the calculated mixing matrices and single-channel images shown in **Fig. 2d** and unmixed them *via* PICASSO. The Pearson correlation coefficients of unmixing of all 32 fluorophore combinations were around 0.99, as shown in the second table. In addition, the determinants of all 2 × 2 sub-matrices of the mixing matrices of the 32 combinations met the non-negativity condition, as shown in the second table. This result indicates that PICASSO can unmix various fluorophore combinations with fixed detection channels. This result also suggests that PICASSO works for a given fluorophore set without changing or optimizing detection channels, even their emission spectra shift few nanometers due to the chemical or environmental factors shown in **Supplementary Note 1**.

| # | Name | Database | Link |
|---|---|---|---|
| 1 | Calretinin | The Human Protein Atlas | https://www.proteinatlas.org/ENSG00000172137-CALB2/brain |
| 2 | ZNF3 | The Human Protein Atlas | https://www.proteinatlas.org/ENSG00000166526-ZNF3 |
| 3 | NeuN | The Human Protein Atlas | https://www.proteinatlas.org/ENSG00000167281-RBFOX3 |
| 4 | PV | Allen Brain Atlas | http://mouse.brain-map.org/gene/show/19056 |
| 5 | Keratin 19 | The Human Protein Atlas | https://www.proteinatlas.org/ENSG00000171345-KRT19/tissue |
| 6 | H3F3A | The Human Protein Atlas | https://www.proteinatlas.org/ENSG00000163041-H3F3A/tissue |

**Supplementary Table 4**. **Databases used to cross-validate the protein expression patterns observed by PICASSO**.

| Filter number | CWL (nm) | FWHM (nm) | Vendor | Product number | Fluorophore (wavelength of the used excitation laser) |
|---|---|---|---|---|---|
| 1 | 425 | 25 | Edmund | #87-787 | CF405S (Exc: 405 nm) |
| 2 | 466 | 45.3 | Semrock | FF01-466/40-25 | ATTO390 (Exc: 405 nm) |
| 3 | 504 | 17 | Semrock | FF01-504/12-25 | CF488A (Exc: 488 nm) |
| 4 | 540 | 55.6 | Semrock | FF01-540/50-25 | CF405L (Exc: 405 nm)<br>ATTO514 (Exc: 488 nm) |
| 5 | 575 | 20.1 | Semrock | FF01-575/15-25 | CF568 (Exc: 561 nm) |
| 6 | 607 | 42 | Edmund | #84-102 | ATTORho101 (Exc: 561 nm) |
| 7 | 656 | 10 | Andover | 656HC10-25 | CF633 (Exc: 637 nm) |
| 8 | 680 | 47 | Semrock | FF01-680/42-25 | ATTO490LS (Exc: 488 nm)<br>CF660R (Exc: 637 nm) |

**Supplementary Table 5. List of the bandpass filters used with the confocal microscopy system.** Exc: wavelength of an excitation laser.

| # | Microscope | Figure | Used antibodies and fluorophores |
|---|---|---|---|
| 1 | Nikon C2 plus | Fig. 1e–t | NeuN (CF405S), CALB2 (CF405M), GFAP (ATTO390), CALB1 (CF405L), SOX2 (Alexa488), laminB1 (ATTO514), Neurofilament-M (ATTO532), beta-tubulin (ATTO490LS), ABAT (CF568), SLC2A1 (ATTORho101), parvalbumin (ATTO594), Iba1 (CF633), CNP1 (CF660R), ZNF3 (CF680R) and VAMP2 (ATTO725). |
| 2 | Nikon C2 plus | Fig 2c,d | NeuN (CF405S), CNP1 (CF488A), SLC2A1 (CF568), parvalbumin (CF633) and GFAP (CF405L). |
| 3 | Nikon C2 plus | Fig. 3b–e | Parvalbumin (CF488A), rabbit NeuN (ATTO514), rabbit GFAP (ATTO532), mouse GFAP (CF405S) and guinea pig NeuN (CF660R). |
| 4 | Leica SP8 | Fig. 4a–m | Parvalbumin (CF405S), CALB1 (CF405L), NeuN (CF488A), GFAP (ATTO514), CALB2 (ATTO490LS), SLC2A1 (CF568), ZNF3 (CF633) and laminin (CF660R). |
| 5 | Leica SP8 | Fig. 4n–r | NeuN (CF405S), CNP1 (ATTO390), CALB1 (CF405L), CALB2 (CF488A), GFAP (ATTO514), MAP2 (ATTO490LS), neurofilament-H (CF568), SLC2A1 (ATTORho101), neurofilament-L (CF633) and parvalbumin (CF660R). |
| 6 | Nikon C2 plus | Fig. 5a | DAPI, laminB1 (CF488A), GFAP (ATTO514), MAP2 (ATTO490LS), SLC2A1 (CF568), parvalbumin (ATTORho101), NeuN (CF633) and CALB2 (CF660R). |
| 7 | Andor dragonfly | Fig. 5b–i | LaminB1 (CF405S), CNP1 (ATTO390), GFAP (CF405L), SV2A (CF488A), SLC2A1 (ATTO514), MAP2 (ATTO490LS), parvalbumin (CF568), NeuN (ATTORho101), CALB2 (CF633) and ZNF3 (CF660R). |
| 8 | Nikon C2 plus | Fig. 6 | Keratin19 (CF488A), Histone H3 (ATTO514), COXIV (CF568) and vimentin (ATTORho101). |

| | | | |
|---|---|---|---|
| 9 | Nikon C2 plus | Supple. Fig. 2 | MAP2 (CF488A) and NeuN (ATTO514). |
| 10 | Nikon C2 plus | Supple. Fig. 10 | MAP2 (ATTO488) and NeuN (ATTO514) |
| 11 | Nikon C2 plus | Supple. Fig. 12a | CALB2 (CF568), rabbit GFAP (ATTORho101) and chicken GFAP (CF633). |
| 12 | Leica SP8 | Supple. Fig. 12f | Lamin A/C (CF405), GM130 (ATTO390) and vimentin (CF405L). |
| 13 | Nikon C2 plus | Supple. Fig. 12g | Lamin A/C (CF488A), GM130 (ATTO514) and vimentin (ATTO490LS). |
| 14 | Nikon C2 plus | Supple. Fig. 12h | Lamin A/C (CF568) and GM130 (ATTORho101). |
| 15 | Nikon C2 plus | Supple. Fig. 12i | Lamin A/C (CF633) and GM130 (CF660R). |
| 16 | Nikon C2 plus | Supple. Fig. 15 | Lamin A/C (CF488A) and GM130 (ATTO514). |
| 17 | Leica SP8 & Nikon C2 plus | Supple. Fig. 16 | Same as Supple. Fig. 12f–i. |
| 18 | Nikon C2 plus | Supple. Fig. 22a–r | DAPI, *Polr2a* (Alexa Fluor 488 and ATTORho101) and *Eif1a* (Alexa Fluor 546). |
| 19 | Nikon C2 plus | Supple. Fig. 22s–v | DAPI, *Gapdh* (Alexa Fluor 488) and vimentin (ATTO514). |
| 20 | Nikon C2 plus | Supple. Fig. 23a,b | NeuN (CF488A) and GFAP (ATTO514). |
| 21 | Nikon C2 plus | Supple. Fig. 23c,d | NeuN (CF488A), GFAP (ATTO514), CALB2 (CF568) and neurofilament-H (ATTORho101). |

| | | | |
|---|---|---|---|
| 22 | Leica SP8 | Supple. Fig. 24b,c | DAPI, laminB1 (CF488A), SLC2A1 (ATTO514), CALB1 (ATTO490LS), parvalbumin (CF568), NeuN (ATTORho101), GFAP (CF633) and CALB2 (CF660R). |
| 23 | Leica SP8 | Supple. Fig. 25 | DAPI, laminB1 (CF488A), YFP, parvalbumin (CF568), SLC2A1 (ATTORho101), GFAP (CF633) and CALB2 (CF660R). |
| 24 | Nikon C2 plus | Supple. Fig. 26 | Keratin 19 (CF488A) |
| 25 | Nikon C2 plus | Supple. Fig. 27a–e | NeuN (CF405S), parvalbumin (CF405M), GFAP (ATTO390), CALB1 (CF405L), SOX2 (Alexa Fluor 488), laminB1 (ATTO514), neurofilament-M (ATTO532), beta-tubulin (ATTO490LS), ABAT (CF568), SLC2A1 (ATTORho101), guinea pig Iba1 (ATTO594), ZNF3 (CF633), CNP1 (CF660R), CALB2 (CF680R) and VAMP2 (Alexa Fluor 790) |
| 26 | Nikon C2 plus | Supple. Fig. 27f–j | NeuN (CF405S), CALB2 (CF405M), GFAP (ATTO390), CALB1 (CF405L), SOX2 (Alexa Fluor 488), laminB1 (ATTO514), neurofilament-L (Alexa Fluor 532), beta-tubulin (ATTO490LS), neurofilament-M (CF568), SLC2A1 (ATTORho101), parvalbumin (ATTO594), Iba1 (CF633), CNP1 (CF660R), ZNF3 (CF680R) and VAMP2 (ATTO725). |
| 27 | Nikon C2 plus | Supple. Fig. 28 | 1$^{st}$ round: parvalbumin (CF488A) and NeuN (ATTO514), 2$^{nd}$ round: CALB2 (CF488A) and SLC2A1 (ATTO514). |

**Supplementary Table 6. Microscopy, antibodies, and fluorophores used for the PICASSO imaging.**

| Product name | Vendor | Product number |
|---|---|---|
| **Cell culture** | | |
| Minimum essential medium (MEM) | Thermofisher | 11095114 |
| Dulbecco's modified eagles' medium (DMEM) | Thermofisher | 11995065 |
| Penicillin-streptomycin | Thermofisher | 15140122 |
| Sodium pyruvate | Thermofisher | 11360070 |
| Fetal bovine serum (FBS) | Thermofisher | 10082147 |
| Bovine calf serum (BCS) | Thermofisher | 26170043 |
| Nunc Lab-Tek chambered coverglass | Thermofisher | 155380PK |
| **FFPE sample preparation** | | |
| Xylenes | Sigma | 534056 |
| Ethyl alcohol, Pure | Sigma | E7023 |
| Sodium citrate tribasic dihydrate | Sigma | C8532 |
| Tween 20 | Sigma | P1379 |
| **Fixation and staining** | | |
| 16% paraformaldehyde (PFA) | Electron Microscopy Science | 15710 |
| Glycine | Sigma | 50046 |
| Sodium azide | Sigma | 71289 |
| Triton X-100 | Sigma | X100 |
| 10× PBS | Invitrogen | AM9625 |
| Normal goat serum | Jackson Immunoresearch | 055-000-121 |
| Normal donkey serum | Jackson Immunoresearch | 017-000-121 |
| Normal rabbit serum | Jackson Immunoresearch | 011-000-120 |
| Normal chicken serum | Jackson Immunoresearch | 003-000-120 |
| Guinea pig anti-NeuN antibody | Synaptic systems | 266 004 |
| Guinea pig anti-Iba1 antibody | Synaptic systems | 234 004 |
| Mouse anti-GFAP antibody | Synaptic systems | 173 011 |
| Chicken anti-GFAP antibody | Aves labs | GFAP |
| Chicken anti-MAP2 | Abcam | ab5392 |
| Chicken anti-MBP | Aves labs | MBP |
| Chicken anti-GFP | Aves labs | GFP-1020 |
| Chicken anti-vimentin | Merck Millipore | AB5733 |
| Goat anti-Chicken IgY (H+L) Secondary Antibody, Alexa Fluor 488 | Thermofisher | A-11039 |
| Goat Anti-Chicken IgY (H+L), Highly Cross-Adsorbed, CF633 | Biotium | 20126-1mg |

| | | |
|---|---|---|
| Goat Anti-Guinea Pig IgG (H+L), Highly Cross-Adsorbed, CF660R | Biotium | 20496-1mg |
| Goat Anti-Mouse IgG (H+L), CF405S | Biotium | 20080-1mg |
| **Fluorophore conjugation** | | |
| Sodium bicarbonate | Sigma | S6297 |
| NAP-5 columns | Cytiva | 17-0853-02 |
| Amicon® Ultra-0.5, 30K MWCO | Merck Millipore | Z740174 |
| AffiniPure Fab Fragment Goat Anti-Rabbit IgG, Fc fragment specific | JacksonImmunoReserach | 111-007-008 |
| AffiniPure Fab Fragment Goat Anti-Chicken IgY (IgG), Fc Fragment Specific | JacksonImmunoReserach | 103-007-008 |
| AffiniPure Goat Anti-Chicken IgY (IgG) (H+L) | JacksonImmunoReserach | 103-005-155 |
| AffiniPure Goat Anti-Guinea Pig IgG (H+L) | JacksonImmunoReserach | 106-005-003 |
| AffiniPure Goat Anti-Rabbit IgG (H+L) | JacksonImmunoReserach | 111-005-144 |
| AffiniPure Donkey Anti-Mouse IgG (H+L) | JacksonImmunoReserach | 715-005-151 |
| **RNAscope** | | |
| RNAscope Fluorescent Multiplex Detection Reagents | ACD | 320851 |
| Probe-mm-*Gapdh* | ACD | 314091 |
| Ethyl alcohol, Pure | Sigma | E7023 |
| **HCR** | | |
| Ethyl alcohol, Pure | Sigma | E7023 |
| 20x saline sodium citrate (SSC) | Thermofisher | AM9763 |
| Tween 20 | Sigma | P1379 |
| HCR probe hybridization buffer | Molecular Instruments | |
| HCR probe | Molecular Instruments | |
| HCR probe wash buffer | Molecular Instruments | |
| HCR amplification buffer | Molecular Instruments | |
| HCR hairpin amplifier h1 | Molecular Instruments | |
| HCR hairpin amplifier h2 | Molecular Instruments | |
| DAPI | Sigma | D9542 |
| **Expansion microscopy** | | |
| Sodium acrylate | Sigma | 408220 |
| Acrylamide | Sigma | A9099 |
| N, N'-methylenebisacrylamide (BIS) | Sigma | M7279 |
| 4-hydroxy-TEMPO (H-TEMPO) | Sigma | 176141 |
| Ammonium persulfate (APS) | Sigma | A3678 |
| N,N,N',N'-tetramethylethylenediamide | Sigma | T7024 |

| | | |
|---|---|---|
| (TEMED) | | |
| Acryloyl-X SE (AcX) | Thermofisher | A-20770 |
| Proteinase K | New England Biolabs | P8107S |
| Trizma hydrochloride (Tris-HCl), pH8.0, 1 M | Sigma | T3038 |
| Ethylenediaminetetraacetic acid (EDTA) | Sigma | EDS |
| Sodium chloride | Sigma | 71376 |
| Triton X-100 | Sigma | X100 |
| **SHIELD** | | |
| SHIELD buffer solution | Lifecanvas technologies | SH-BS |
| SHIELD-ON buffer | Lifecanvas technologies | SH-ON |
| SHIELD-Epoxy solution | Lifecanvas technologies | SH-ES |
| Sodium dodecyl sulfate | Sigma | L3771 |
| Boric acid | Sigma | B6768 |
| Sodium sulfite | Sigma | S0505 |
| Sodium hydroxide | Sigma | S8045 |
| **Fluorophore inactivation (t-CyCIF)** | | |
| Hydrogen peroxide solution | Sigma | 216763 |
| Sodium hydroxide | Sigma | S8045 |
| **Direct staining** | | |
| Sox2 antibody [Alexa 488 conjugated] | Thermofisher | 53-9811-82 |
| NF-L Antibody (7D1) [Alexa Fluor® 532] | Novus Biologicals | NBP2-50612AF532 |

**Supplementary Table 7. List of the materials used in this study.**
